# Cooperative CTCF-nucleosome oligomerization stabilizes chromatin loop anchors

**DOI:** 10.64898/2026.02.06.704447

**Authors:** Manuel Osorio Valeriano, Alexander C. Stone, Masahiro Nagano, Bonnie Su, Laura Caccianini, Anders S. Hansen, Lucas Farnung, Seychelle M. Vos

## Abstract

Eukaryotic DNA is organized across multiple scales to support genome compaction, appropriate gene expression, and DNA recombination. A central player in these processes is the CCCTC binding factor (CTCF), which defines specific chromatin loop structures and insulates enhancer elements from promoters. Chromatin is organized in a distinct pattern around CTCF-bound sites; however, the role of this patterning remains unclear. Here, we report cryo-electron microscopy structures of reconstituted CTCF-nucleosome complexes, revealing that CTCF dimerization promotes the oligomerization of nucleosomes into defined higher-order assemblies involving specific histone-histone and CTCF-CTCF interactions. Notably, CTCF does not oligomerize efficiently on non-chromatinized DNA substrates. Disruption of identified CTCF interaction interfaces in cells results in a marked decrease in chromatin looping and functionally impairs cellular differentiation. These results indicate that chromatin structure at CTCF sites plays an important role in supporting higher-order interactions between distal regions of the genome and that these interactions are important for supporting cell-type-specific gene expression.

## Introduction

The CCCTC binding factor (CTCF) is an essential DNA-binding protein with central roles in gene expression regulation, DNA repair and recombination, and development^1–7^. Functionally, CTCF is responsible for the formation of chromatin loops that demarcate regulatory domains, enhancer insulation, and long-range chromatin interactions that are critical for proper transcriptional control^4,6–13^. Chromatin loops are thought to be formed largely through the action of cohesin, which can actively extrude DNA loops^14^. CTCF binds DNA in a directional manner through its recognition of an asymmetric DNA binding motif via 11 zinc fingers (ZnFs) that are flanked by flexible N- and C-termini (Figure 1a)^15–17^. This directional binding is responsible for delimiting how far cohesin extrudes DNA loops^18–21^. Finally, CTCF has been reported to associate with itself; however, the functional role for this association has not been established ^22–25^. Thus, the directional association of CTCF with DNA has profound consequences on both local chromatin structure and 3D genome organization, but the role of CTCF-CTCF interactions in these functions remains unknown.

**Figure 1:**
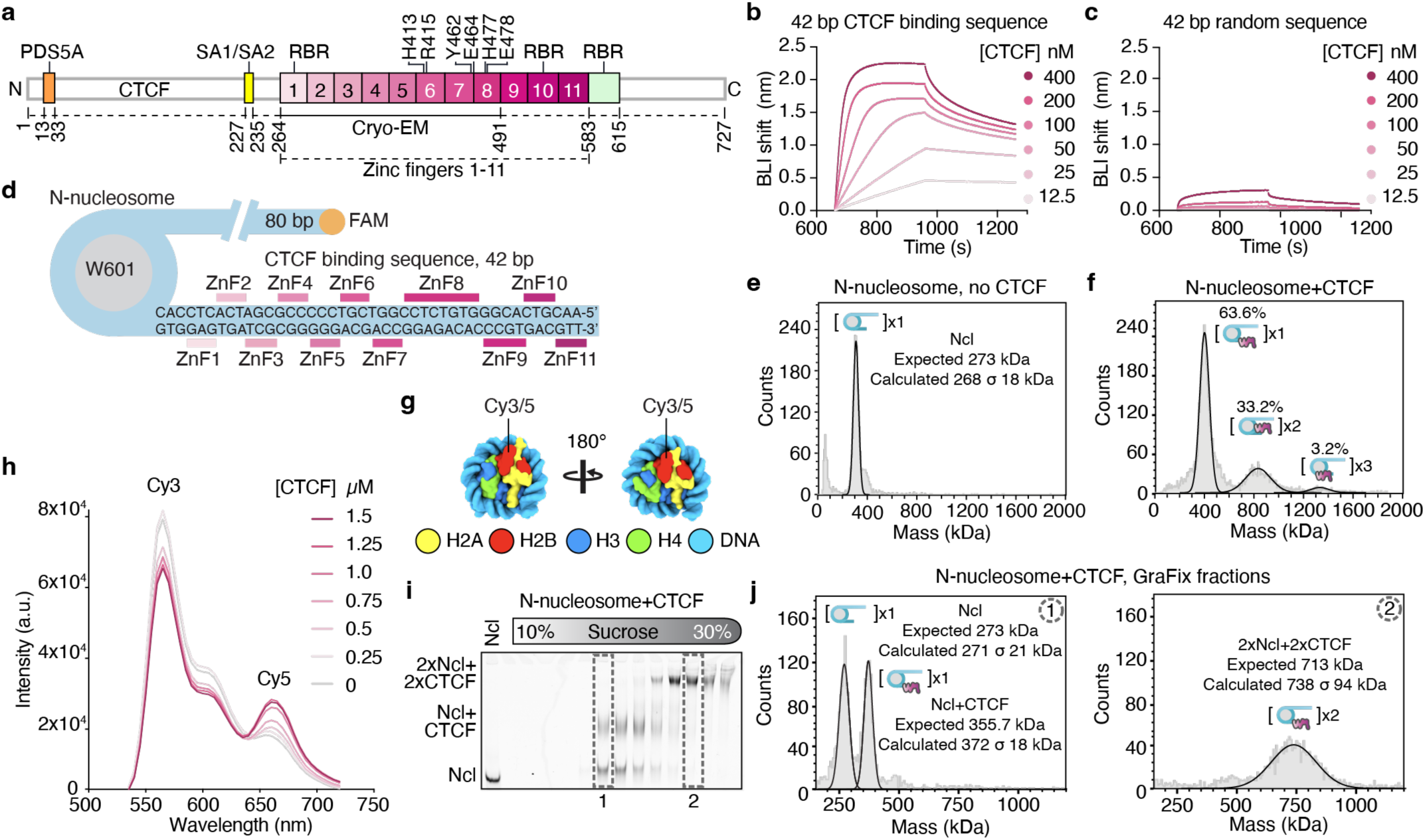
CTCF promotes nucleosome oligomerization. **a,** Domain architecture of full-length human CTCF. Amino acid boundaries are indicated below and residues mutated in this study are shown above the schematic. Solid line indicates the protein region modeled in cryo-EM reconstructions reported here. Regions of CTCF previously described to interact with proteins SA1/SA2, PDS5A, and RNA-binding regions (RBR) are demarcated. **b,c,** Full-length CTCF binding to linear DNA as detected by Biolayer interferometry (BLI). Sensorgrams are shown for increasing concentrations of CTCF. Consensus 42 base pairs (bp) CTCF binding sequence (**b**) or a random 42 bp sequence (**c**). **d,** Schematic of the reconstituted mononucleosome containing the consensus 42 bp CTCF binding sequence to position CTCF with its N-terminus (N-nucleosome) facing the Widom 601 (W601) nucleosome positioning sequence. The sequence of the CTCF binding motif and the relative position of the individual zinc finger contacts (ZnF1-11) are shown. The orange circle represents the FAM fluorophore attached to the nucleosomes for detection. **e,f,** Mass photometry analysis of N-nucleosome alone (**e**) and in the presence of CTCF (**f**) performed under native, non-crosslinked conditions using the rapid dilution MassFluidix HC system. The expected and calculated masses of the free nucleosome are indicated in (**e**). Distinct populations with masses corresponding to monomeric (x1), dimeric (x2), and trimeric (x3) nucleosome-CTCF species are present in (**f**), with the percentage of each species indicated. **g,** Schematic of the Cy3- and Cy5-labeled nucleosomes used for the FRET assay shown in (**h**). Fluorophores were conjugated to a cysteine substituted at residue 112 of histone H2B (H2B Thr112Cys) via maleimide-based labeling chemistry. **h,** Fluorescence emission spectra of Cy3/Cy5-labeled nucleosomes upon titration with increasing concentrations of wild-type CTCF (0-1.5 µM). The donor (Cy3, ∼565 nm) and acceptor (Cy5, ∼660 nm) emission peaks are indicated. Addition of increasing CTCF concentrations results in a progressive increase in Cy5 and decrease in Cy3 emission, consistent with increased FRET efficiency between donor- and acceptor-labeled nucleosomes. **i,** Native gel analysis of fractions collected from gradient fixation (GraFix) of FAM-labeled N-nucleosome after incubation with CTCF. Numbered lanes correspond to fractions used in mass photometry analysis. Gradient fractions were analyzed on 3.5% acrylamide Tris-Borate-EDTA (TBE) native gel run in 0.2x TBE buffer; a control nucleosome (Ncl) was included in the first lane. Positions corresponding to free nucleosome (Ncl), CTCF-bound nucleosome (Ncl+CTCF), and higher-order species corresponding to a dimeric complex with 2 nucleosomes and 2 CTCF molecules (2×Ncl+2×CTCF) are indicated. **j,** Mass photometry analysis of GraFix-purified N-nucleosome-CTCF complexes. The left and right panels show fractions 1 and 2 from (**i**), respectively. Expected and calculated masses are indicated.

Upon initial binding of CTCF to its cognate sequence motif, CTCF is thought to recruit chromatin remodelers, including ISWI, SWI/SNF, and CHD family members^26–30^. This results in the establishment of an asymmetric nucleosome-free region flanking the CTCF binding site, with 80-110 base pairs (bps) on the N-terminal side and 40-80 bp on the C-terminal side of CTCF, and the formation of phased nucleosome arrays with up to 10 well-positioned nucleosomes on each side of CTCF^29–33^. The disruption of positioned nucleosomes abutting CTCF alters 3D genome interactions and is manifested in reduced looping and may affect genomic insulation^29,33,34^. CTCF interacts with cohesin via its N-terminus, and the larger nucleosome free region on the N-terminal side of CTCF may provide sufficient space to accommodate and stall cohesin^20,21,35^. This suggests that CTCF influences 3D genome organization both through its direct association with cohesin and its direct influence on nucleosome organization.

Recent work has begun to elucidate how cohesin associates with CTCF^20,21,35^, but it remains unclear how nucleosome phasing at CTCF sites and CTCF self-interactions influence 3D genome organization. Here, we have systematically addressed these questions using biochemical, structural, and 3D genomic approaches in cells. Together, our results suggest that CTCF and nucleosomes work cooperatively to induce distinct chromatin conformations that enhance looping interactions genome-wide.

## Results

### CTCF forms higher-order assemblies with nucleosomes

To address how CTCF functions within a chromatin context, we first purified human CTCF and assessed its association with linear DNA fragments bearing either the complete CTCF-binding sequence motif or a random sequence. As previously observed, CTCF readily associated and slowly dis-associated from linear DNA containing its binding motif^18,19,36^. In contrast, binding to random DNA sequences was negligible across the same concentration range (Figure 1b, c and Extended Data Figure 1a, b). Together, these results confirm that our purified full-length CTCF rapidly and specifically associates with DNA containing its sequence motif.

We next investigated CTCF association with mononucleosomal substrates. DNA sequences containing 80 bp DNA overhangs, followed by a Widom 601 nucleosome positioning sequence and either a 42 bp CTCF consensus motif or a random 42 bp sequence, were used to reconstitute nucleosomes. The CTCF motif was provided to position CTCF with its N-terminus (N-nucleosome) facing the nucleosome (Figure 1d, Methods). This positioning was chosen to mimic initial binding of CTCF to DNA prior to nucleosome remodeling as has been reported in cells^30^. We used mass photometry to assess CTCF-nucleosome complexes in solution. In the absence of CTCF, nucleosomes were detected exclusively as monomers (Figure 1e). Notably, upon addition of CTCF, new populations consistent with nucleosome-CTCF complexes and higher-molecular weight species corresponding to CTCF-bound nucleosome dimers (two nucleosomes and two CTCF molecules) and trimers (three nucleosomes and three CTCF molecules) were observed (Figure 1f). Finally, a FRET assay using Cy3- and Cy5-labeled H2B(Thr112Cys) N-nucleosomes showed that increasing concentrations of CTCF led to a progressive increase in FRET (Figure 1g, h), demonstrating that CTCF induces oligomerization of mononucleosomes in solution.

To isolate distinct CTCF-nucleosome complexes, we used sucrose density gradient ultracentrifugation (Extended Data Figure 1c). Glutaraldehyde was either included in the sucrose gradient (<u>Gra</u>dient <u>Fix</u>ation (GraFix)) or added after the gradient was run to prevent complex disassociation^37^. Native PAGE and mass photometry analysis confirmed the presence of free nucleosomes, CTCF-bound nucleosome monomers, and dimers. Similar results were obtained for both crosslinking schemes (Figure 1i, j and Extended Data Figure 1d, e; Methods). As expected, in the absence of CTCF, only a small fraction of higher-molecular weight species was observed (Extended Data Figure 1f, g). Similar observations were made for a nucleosome lacking the 80 bp extranucleosomal DNA (0-W601-N-nucleosome) (Extended Data Figure 1h-j). Consistent with our experiments on linear DNA, CTCF did not associate with nucleosomes containing a random DNA sequence (Extended Data Figure 1k). Finally, GraFix experiments using non-chromatinized, motif-containing 42 bp DNA displayed only monomeric CTCF-DNA complexes and no detectable higher-order populations (Extended Data Figure 1l-n). Together, these results show that CTCF in complex with mononucleosomes forms higher-order assemblies.

### Cryo-EM structure of CTCF-nucleosome complex

We next visualized CTCF on the N-nucleosome template using cryogenic-electron microscopy (cryo-EM). We formed the CTCF-mononucleosome complex and separated oligomeric states using GraFix. Fractions containing dimeric species were pooled, buffer exchanged, concentrated, and used for the preparation of single-particle cryo-EM samples (Extended Data Figure 2a). Initial inspection of aligned micrographs and 2D reconstructions revealed stacked nucleosome dimers, with each extended extranucleosomal DNA showing density for CTCF (Figure 2a and Extended Data Figure 2b, c). Extensive 3D classification resulted in a final map with a resolution of 3.7 Å (0.143 gold standard Fourier Shell Correlation (FSC)), with core nucleosome densities corresponding to 3.2-6.3 Å and CTCF densities to >6.5 Å. A map focused on the nucleosome dimer stack was resolved to 2.8 Å (Figure 2b, Extended Data Figure 2d-f and Extended Data Figure 3). Zinc fingers (ZnFs) 1-8 from one CTCF molecule and 1-7 from the second were observed. The structure was modeled using a high-resolution structure of a *Xenopus laevis* nucleosome and previous X-ray crystal structures of CTCF bound to its cognate DNA sequence^15–17,38^. Although the CTCF density is of lower resolution, the ZnF density is consistent with high-resolution structures of motif-bound CTCF and was modeled accordingly. The final model was manually adjusted, refined, and showed good stereochemistry at the observed resolutions (Extended Data Table 1 and Extended Data Figure 2g-p).

**Figure 2:**
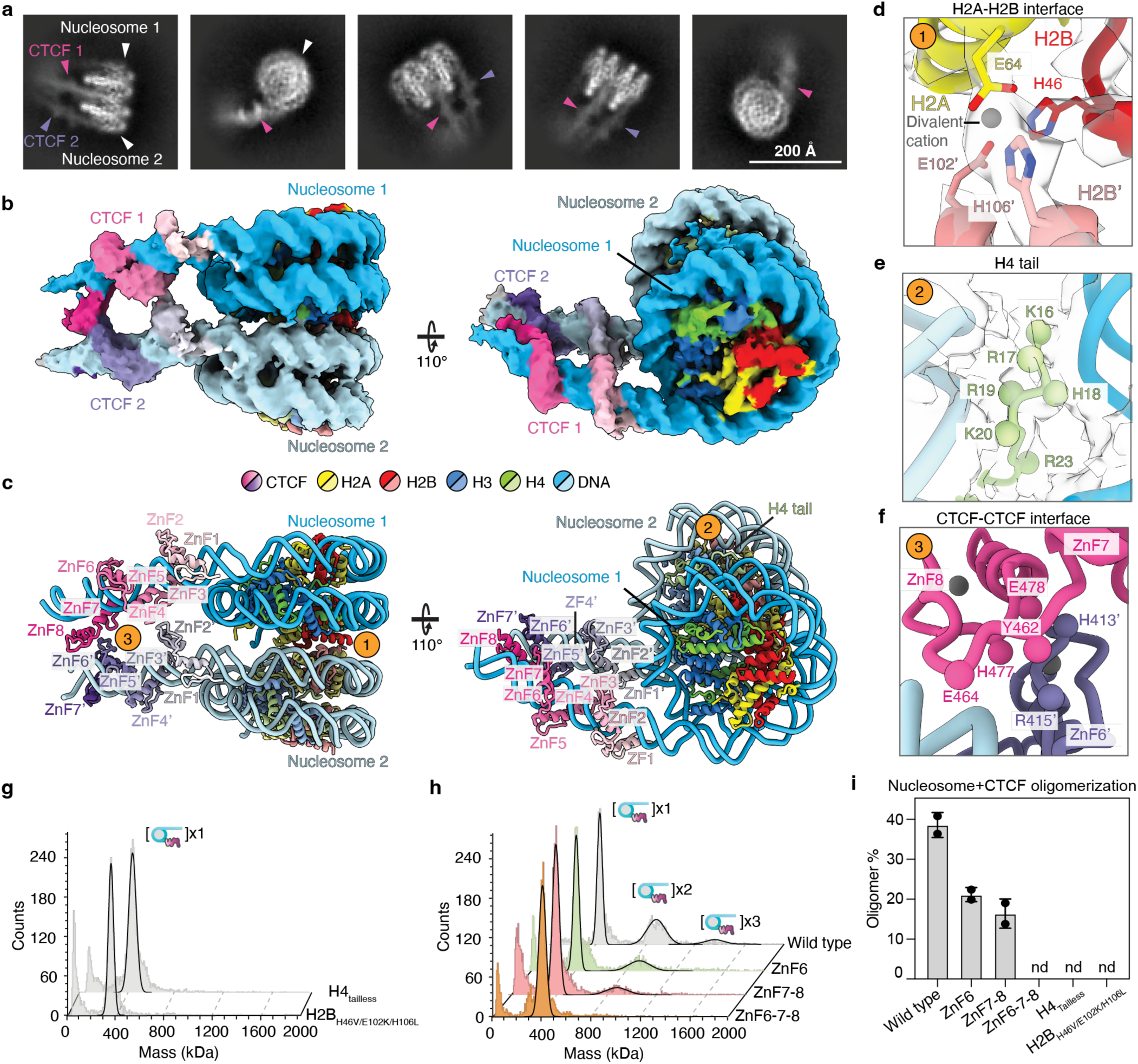
Cryo-EM structure of a CTCF-nucleosome dimer. **a,** Representative two-dimensional (2D) classes of CTCF-nucleosome dimers. **b,** Cryo-EM density map of CTCF-nucleosome dimer at 3.7 Å resolution (Map B, Extended Data Fig. 2d, e). Color code is shown at the bottom. **c,** Model of CTCF-nucleosome dimer. Regions mediating dimerization are indicated: (1) the interface between H2A and H2B from Nucleosomes 1 and 2, (2) the H4 tail from Nucleosome 2, (3) the interface between ZnF7-8 from CTCF 1 and ZnF6’ from CTCF 2. Colors as in panel **b**. **d,** H2A-H2B interaction interface. Density from Map A is shown as a transparent surface. **e,** H4 N-terminal tail from Nucleosome 2 sitting between the stacked nucleosomes. Density from Map A is shown as a transparent surface. **f,** CTCF-CTCF interaction interface between ZnF6’ from CTCF 2 and ZnF7-8 from CTCF 1. Putative residues mediating this interaction are indicated. Due to limited resolution of our structure in this region, these residues were selected based on their relative positioning to the C*a* backbone. **g,h,** Native, non-crosslinked mass photometry analysis of N-nucleosomes containing the histone H2B mutant (His46Val, Glu102Lys, His106Leu) or the H4 tailless mutant (Δ1-19) incubated with wild-type CTCF (**g**) or N-nucleosomes incubated with wild-type CTCF or the indicated CTCF zinc finger mutants (**h**). Peaks corresponding to monomeric (×1), dimeric (×2), and trimeric (×3) nucleosome-CTCF species are indicated. **i,** Quantification of CTCF-induced nucleosome oligomerization for wild-type CTCF, the indicated CTCF zinc finger mutants, and the H2B and H4 tailless nucleosome mutants, based on native, non-crosslinked mass photometry shown in (**g,h**). Bars represent mean ± standard deviation from two replicates; nd, no oligomeric species detected. Individual data points are shown.

The structure showed two CTCF-bound mononucleosomes forming a pseudo-symmetric dimer stack (Figure 2c). This type I histone stacking interaction has been observed in previous structures containing nucleosomal arrays or nucleosomes with linker histone H1, respectively^39–41^. Pseudosymmetric contacts are made across histone H2A-H2B dimers (Figure 2c,d). Interestingly, we observed a small additional density that appears to be coordinated by residues H2A Glu64 and H2B His46 on the same nucleosome, and residues H2B His106 and Glu102 on the opposing nucleosome (Figure 2d). We have tentatively assigned the additional density to a Zn^2+^ ion, as zinc was present in our reactions and has been implicated to mediate a similar interaction between yeast nucleosomes^42^. Present cryo-EM methods do not allow for the experimental identification of ions. Finally, one of the H4 N-terminal tails intercalated between the DNA duplexes of the stacked nucleosomes but did not interact with the H2A-H2B acidic patch (Figure 2c, e)^41,43,44^.

Our model suggests that CTCF dimerizes in a parallel orientation, with the two CTCF molecules asymmetrically interacting with each other using a binding interface between ZnF6-7-8. ZnF6 residues His413 and Arg415 from one CTCF molecule appear to be positioned near ZnF7 residues Tyr462 and Glu464, and ZnF8 residues His477 and Glu478 on the opposing CTCF molecule (Figure 2c, f). As modeled here and observed in high resolution X-ray crystal structures, CTCF recognizes its sequence motif through interactions between the ZnFs and bases in the major groove of the DNA^15–17^. The only exception to this binding mode is ZnF8, which interacts with the phosphate backbone, bridging across the DNA minor groove. No visible direct interactions between CTCF and the histone proteins were observed. CTCF dimerization has been observed in X-ray crystal structures. These interactions are distinct from those observed here regarding the involved ZnFs (Extended Data Figure 4), and their biological significance has not been experimentally tested^15–17^. Our results show that CTCF dimerizes effectively when bound to a mononucleosomal substrate by promoting type I nucleosome stacking and using an interaction interface comprising ZnF6-7-8.

### Disruption of dimerization interfaces reduces higher-order assemblies

To assess the importance of the observed interactions for dimerization, we generated a series of histone and CTCF mutants. CTCF-nucleosome oligomerization was assessed by mass photometry and GraFix experiments. Mutation of H2B interacting residues (His46Val, Glu102Lys, His106Leu) or deletion of the N-terminal H4 tail (Δ1-19) resulted in a severe oligomerization defect in the presence of CTCF (Figure 2g, i and Extended Data Figure 5a,b,g). Next, we generated point-substitution CTCF mutants designed to disrupt the dimerization interface while retaining appropriate ZnF folding (Figure 2f, Methods). Three constructs containing point mutations were cloned and purified (ZnF6, His413Gly and Arg415Ala; ZnF7-8, Tyr462Gly, Glu464Val, His477Val, and Glu478Arg; and ZnF6-7-8, combining all substitutions present in the ZnF6 and ZnF7-8 constructs) (Extended Data Figure 1a). Each mutant bound linear DNA similarly to wild-type CTCF (Extended Data Figure 5c). In CTCF-nucleosome oligomerization experiments, ZnF6 and ZnF7-8 displayed an oligomerization defect, whereas ZnF6-7-8 almost completely lost its ability to support oligomerization (Figure 2h, i and Extended Data Figure 5d-g). Supporting these observations, ZnF6-7-8 showed a strong nucleosome FRET defect compared to wild-type CTCF (Extended Data Figure 5h-j). These data indicate that CTCF-nucleosome oligomerization is dependent on both histone-histone and CTCF-CTCF interactions.

The flexible CTCF N- and C-termini are not observed in our cryo-EM maps and have been attributed significant functional roles. Specifically, the CTCF N-terminus is critical for the directional stalling of cohesin, mediates interactions with cohesin-associated proteins including PDS5A and SA1/SA2, and is reported to dimerize on its own^18,20,21,25,35^ (Figure 1a). The C-terminus of CTCF, in addition to ZnFs 1 and 10, can associate with RNA^24,45,46^. We thus next tested whether the CTCF N- or C-termini played a role in CTCF-nucleosome oligomerization. We purified a CTCF construct containing only ZnFs 1-11 (CTCF residues 263-581) to investigate whether the ZnFs alone could mediate dimerization (Extended Data Figure 1a). ZnF1-11 were sufficient to mediate dimerization and higher-order interactions with mononucleosomes (Extended Data Figure 5k, l). Furthermore, neither CTCF terminus associated with DNA under our assay conditions (Extended Data Figure 5m). These results show that the CTCF ZnFs are necessary and sufficient for CTCF-mononucleosome oligomerization.

Finally, our structure suggested that the orientation of CTCF relative to the nucleosome could influence dimerization efficiency. Inverting the CTCF motif so that the CTCF

C-terminus faced the nucleosome (C-nucleosome_register 0) was predicted to abolish the dimerization interface (Extended Data Figure 6a, b). Indeed, this nucleosome was impaired in forming CTCF-nucleosome dimers (Extended Data Figure 6c). Because motif inversion also alters the rotational phasing of CTCF relative to the nucleosome, we asked whether dimerization could be restored by adjusting motif spacing to recover a permissive helical register. Systematically varying the motif position across a complete DNA helical turn predicted that the ZnF6-7-8-mediated interface could form when the motif was placed 6 or 7 bp from the nucleosome (Extended Data Figure 6a,b). As predicted, positioning the motif at either distance restored dimerization to levels observed for the N-nucleosome (Extended Data Figure 6d-g). Thus, dimerization requires a defined geometric arrangement of CTCF relative to the nucleosome, consistent with a specific ZnF6-7-8 interface rather than nonspecific association.

To test whether CTCF-induced nucleosome oligomerization is retained when CTCF is positioned at a greater distance from the nucleosome, as is commonly observed in cells, we generated sequences that mimic native spacing between nucleosomes at CTCF sites^30,32,47^. Specifically, we used substrates containing a Widom 601 sequence and a 42 bp CTCF binding motif separated by either 80 bp or 110 bp of linker DNA (Extended Data Figure 7a). We again observed that CTCF enhanced dimerization of these constructs, indicating that longer linker length does not affect the ability of CTCF to induce nucleosome dimerization (Extended Data Figure 7b-e). It is likely that the increased flexibility in the linker permits dimerization over more rotational states than what we observed when CTCF sites were placed in close proximity to the nucleosome.

### CTCF forms higher-order assemblies with dinucleosomal substrates

In cells, nucleosomes are phased around CTCF-engaged sites. Up to 10 regularly spaced nucleosomes are observed on each side of CTCF^30–32^. We thus evaluated whether CTCF-nucleosome oligomerization is retained on DNA substrates where CTCF is flanked by two nucleosomes. We first placed a CTCF binding motif of 34 or 42 bp between Widom 601 and Widom 603 nucleosome positioning sequences (Dinucleosome 34 and Dinucleosome 42) (Figure 3a and Extended Data Figure 7f). Mass photometry analysis of non-cross-linked and GraFix-purified samples showed that both dinucleosome constructs self-associate to some extent in the absence of CTCF (Figure 3b, d, f and Extended Data Figure 7g, i). The addition of CTCF greatly enhanced multimerization of these substrates (Figure 3c, e, g and Extended Data Figure 7h, i).

**Figure 3:**
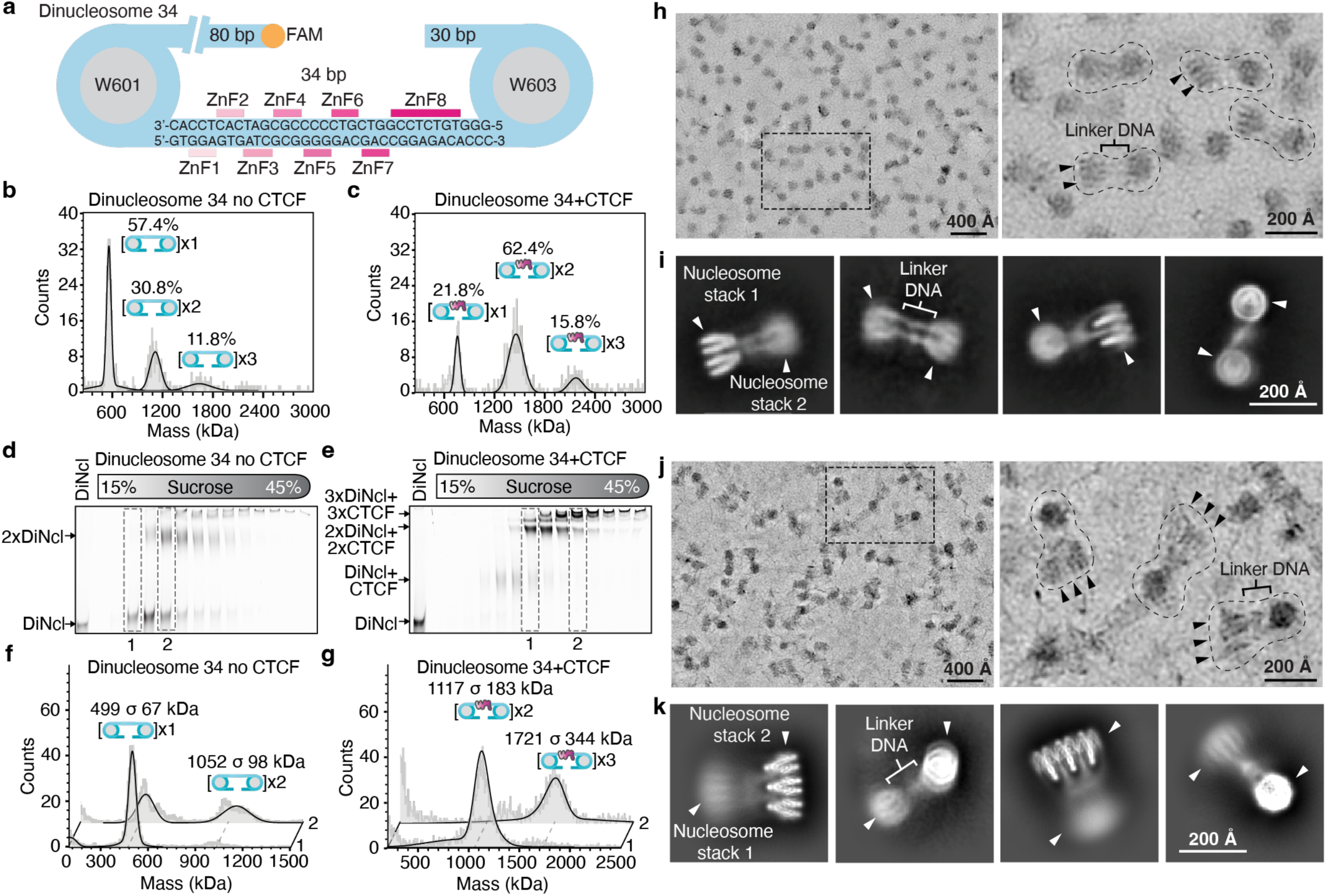
CTCF association with a dinucleosome substrate. **a,** Schematic of a dinucleosome construct (Dinucleosome 34) containing two Widom nucleosome positioning sequences (W601 and W603) connected by linker DNA comprising the first 34 bp of the CTCF binding sequence that can accommodate zinc fingers ZnF1-8. The orientation of the CTCF binding motif relative to the two nucleosomes and the positions of zinc finger contacts (ZnF1–8) are indicated. The orange circle denotes the FAM fluorophore used for detection. **b,c,** Native, non-crosslinked mass photometry analysis of Dinucleosome 34 alone (**b**) or following incubation with CTCF (**c**). Peaks corresponding to monomeric (×1), dimeric (×2), and trimeric (×3) dinucleosome-CTCF species are indicated, with the percentage of each species shown. **d,e,** Native gel analysis of fractions collected from GraFix of Dinucleosome 34 alone (**d**) or after incubation with CTCF (**e**). Gradient fractions (15–45% sucrose) were resolved on 3.5% acrylamide TBE native gels. Positions corresponding to CTCF-bound Dinucleosome 34 (DiNcl+CTCF), CTCF-Dinucleosome 34 dimers (2xDiNcl+2xCTCF), and trimers (3xDiNcl+3xCTCF) are indicated. Free Dinucleosome 34 (DiNcl) was loaded in the first lane as a control. **f,g,** Mass photometry analysis of GraFix fractions 1 and 2 from (**d**) and (**e**). Calculated masses are indicated. Peaks corresponding to monomeric (×1), dimeric (×2), and trimeric (×3) dinucleosome (**f**) or dinucleosome-CTCF complexes (**g**) are indicated, with the calculated masses shown for each peak. **h,** Representative micrograph of the CTCF-Dinucleosome 34 dimer. The dotted-line box indicates the region magnified in the right view. Arrowheads indicate nucleosome positions. Scale bars are indicated. **i**, Representative 2D classes of CTCF-Dinucleosome 34 dimers. Scale bar provided. **j,** Representative micrograph of the CTCF-Dinucleosome 34 trimer. Annotated in the same manner as (**h**). Scale bar provided. **k,** Representative 2D classes of CTCF-Dinucleosome 34 trimers. Annotated in the same manner as (**i**). Scale bar provided.

We next visualized CTCF on the 34 bp dinucleosome substrate by cryo-EM. Cryo-EM samples were prepared identically to those used for the CTCF-mononucleosome structure (Extended Data Figure 7j, Methods). Inspection of aligned micrographs and initial 2D classes revealed dinucleosome dimers with additional, fuzzy density emanating from the well-aligned nucleosome stack, corresponding to a second nucleosome pair (Figure 3h,i). Unfortunately, the flexibility in the dinucleosome complex resulted in low resolution 3D reconstructions precluding us from building a high-confidence structural model. Together, our results show that CTCF dimerizes effectively when bound to nucleosome substrates that more closely resemble the complexity of the native chromatin environment.

Multiple CTCF binding sites are often found at the base of cohesin-mediated loops and at insulator sites^48–51^. Consistent with this, prior super-resolution microscopy work has observed small clusters of several CTCF molecules in cells^52,53^. The presence of multiple CTCF binding sites has been suggested to enhance loop and insulator strength^48,50^. Extrapolation of our CTCF-mononucleosome dimer structure indicated that multiple CTCF-bound nucleosomes could interact using a repeated binding interface with neighboring or distal CTCF-nucleosome complexes (Extended Data Figure 7k). Supporting this idea, we observed the formation of higher-order CTCF-mono and dinucleosome multimers in our mass photometry analysis of non-crosslinked and GraFix-purified samples (Figure 1f and Figure 3c,g). To examine how more than two CTCF and nucleosome complexes are organized, we selected a trimeric CTCF-dinucleosome species from GraFix for cryo-EM analysis (Extended Data Figure 7j).

We observed higher-order dinucleosome assemblies in the aligned micrographs (Figure 3j). 2D classifications confirmed the presence of dinucleosome trimers with a well-aligned nucleosome stack and a fuzzy density corresponding to a second nucleosome stack connected by linker DNA (Figure 3k). Due to the high flexibility of the linker DNA, we were unable to obtain a high-resolution map to confidently build a structural model of the CTCF-dinucleosome trimer. Together, these data show that CTCF-nucleosome units can oligomerize beyond a dimer, forming rigid nucleosome stacks joined by a flexible linker DNA. Such assemblies could provide a structural basis for the enhanced loop and insulator strength associated with clustered CTCF binding sites.

### Disruption of the CTCF dimerization interface impairs growth and differentiation in mESCs

Next, we assessed the physiological relevance of CTCF dimerization. To this end, we engineered mouse embryonic stem cells (mESCs) carrying the Blimp1-mVenus and Stella-ECFP (BVSC) germline reporter transgenes^54–56^ to generate three cell lines harboring point mutations that disrupt CTCF self-interaction in the context of mononucleosomes (Figure 4a). Mirroring our *in vitro* CTCF dimerization mutants, we introduced substitutions in ZnF6 (His413Gly and Arg415Ala; hereafter “ZnF6”), in ZnF7-8 (Tyr462Gly, Glu464Val, His477Val, and Glu478Arg; “ZnF7-8”), or both sets combined (“ZnF6-7-8”) (Figure 4a, Extended Data Figure 8a). Because ZnF6 and ZnF7-8 are encoded by exons 7 and 8, respectively, the ZnF6-7-8 lines were generated by sequential editing of the ZnF7-8 background. We established at least two independent isogenic clones per genotype for downstream analyses.

**Figure 4:**
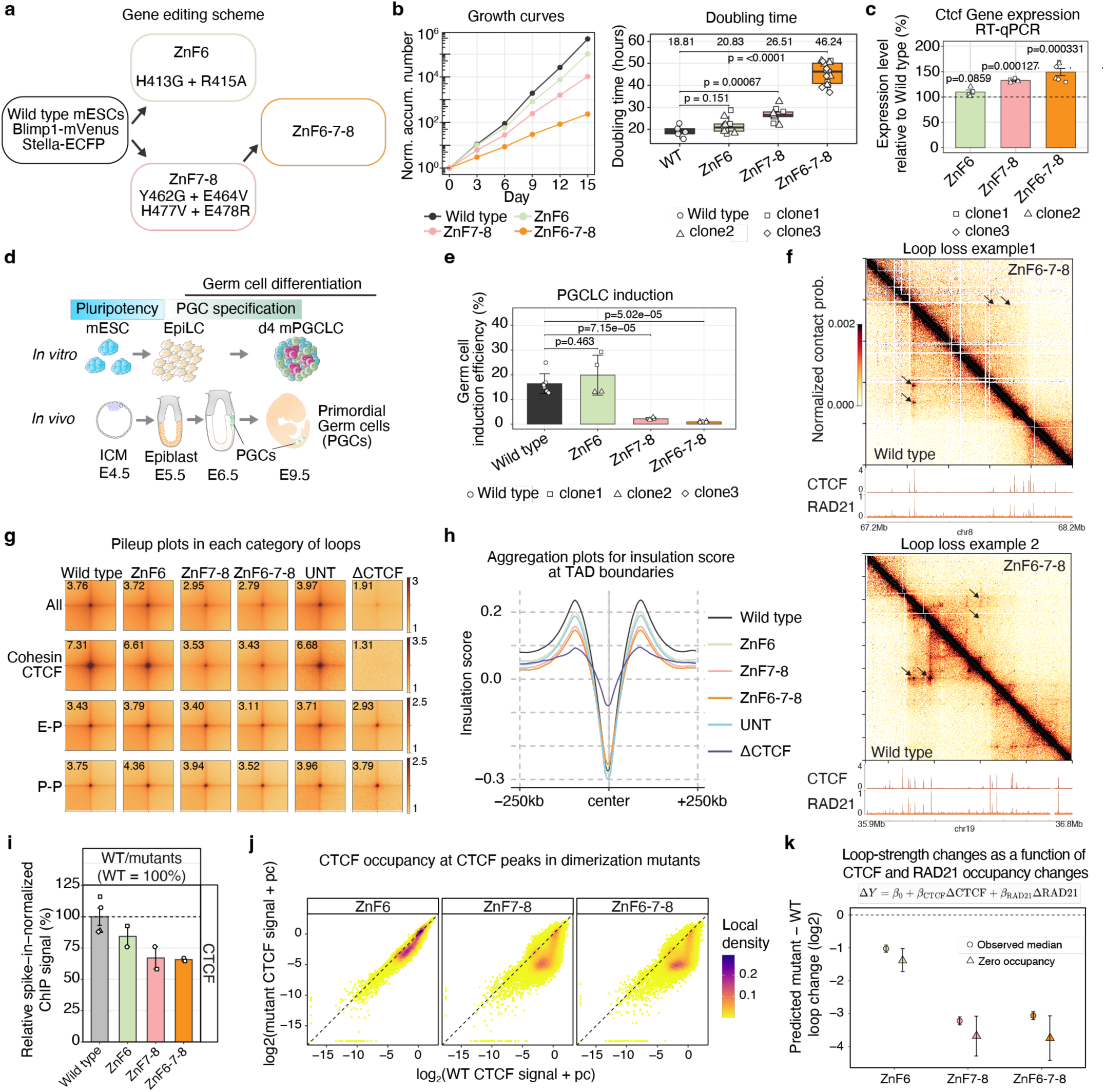
Disruption of the CTCF dimerization interface in mESCs impairs proliferation, germline differentiation, and CTCF-cohesin looping. **a,** Schematic of CRISPR/Cas9-mediated engineering of *Blimp1-mVenus; Stella-ECFP* (BVSC) mouse embryonic stem cells (mESCs) to introduce point mutations in the CTCF dimerization interface. ZnF6 mutant: His413Gly and Arg415Ala; ZnF7-8 mutant: Tyr462Gly, Glu464Val, His477Val, and Glu478Arg; ZnF6-7-8 mutant: combined ZnF6 and ZnF7-8 substitutions. **b,** Growth curves (left) and estimated doubling times (right) for wild type and mutant mESCs during serial passaging for 15 days. Growth curves show accumulated cell numbers normalized to day 0. Doubling-time box plots summarize per-genotype estimates; center line denotes the median, lower and upper hinges denote first and third quartiles, respectively, and upper and lower whiskers extend to the largest and smallest values within the 1.5× interquartile range, respectively; the shape of the points represents independent isogenic clones; P values were calculated using the two-sided Welch’s t-tests. **c,** *Ctcf* mRNA levels measured by RT-qPCR, plotted relative to wild-type (dashed line). Points represent independent clones; bars represent mean, and error bars indicate standard deviation; P values were calculated using two-sided Welch’s t-tests. **d,** Schematic of the *in vitro* reconstitution system for mouse germline development, including differentiation from mESCs to epiblast-like cells (EpiLCs) and subsequent induction of primordial germ cell-like cells (PGCLCs), alongside the corresponding *in vivo* stages. ICM: inner cell mass; PGC: primordial germ cell. Adapted with permission from Nagano M. & Saitou M. Current Opinion in Genetics & Development 94, 102383 (2025).^58,111^ © 2025 Elsevier Ltd. **e,** PGCLC induction efficiency measured by flow cytometry as the fraction of *Blimp1mVenus* (BV) reporter-positive cells after 4 days of induction. The shapes of the points represent independent clones; bars represent mean, and error bars indicate standard deviation; P values were calculated using the two-sided Welch’s t-tests. **f,** Representative Micro-C contact maps illustrating loss of CTCF-cohesin loops in ZnF6-7-8 mutant compared with wild type at two loci in chr8 (top) and chr19 (bottom). Upper triangles, ZnF6-7-8; lower triangles, wild type. Black arrows indicate the positions of loops; CTCF and RAD21 ChIP-seq tracks in wild type are shown below. **g,** Aggregate loop pileups (observed/expected) for the indicated loop classes across genotypes. Rows show all loops (“All”), CTCF-cohesin loops (“Cohesin CTCF”), enhancer-promoter loops (E-P) and promoter-promoter loops (P-P). Columns include wild type and the indicated mutants; “UNT” and “ΔCTCF” represent untreated and acute CTCF depletion conditions from Narducci 2025^62^. The top left number in each plot indicates aggregate peak enrichment score (central pixel value). **h,** Aggregate insulation profiles around TAD boundaries. Insulation scores were aggregated to called boundaries. Lines indicate the mean insulation score profile within ±250 kb of boundary centers for each condition. **i,** Relative spike-in-normalized CTCF ChIP-seq signal in wild type and CTCF dimerization-interface mutants. Signals are expressed relative to the mean wild-type signal, which was set to 100% (dashed line). Bars represent mean and error bars indicate standard error; points represent independent biological replicates. **j,** Peak-level comparison of spike-in-normalized CTCF signal in each mutant versus wild type across the common CTCF peak set. Signals are shown as log2(signal + protein-specific pseudocount). Each point represents one CTCF peak; the dashed line indicates y = x and color indicates local point density. **k,** Mutant–wild-type differences in log2-transformed loop intensity were modeled as a function of the corresponding changes in CTCF and RAD21 signals at loop anchors. Circles show model predictions at the observed median ΔCTCF and ΔRAD21 for each mutant, whereas triangles show predictions at ΔCTCF = ΔRAD21 = 0, representing matched CTCF and RAD21 occupancy. Error bars indicate 95% confidence intervals, and the dashed line indicates no change.

Mutations involving ZnF7-8 caused a pronounced growth defect. ZnF7-8 and ZnF6-7-8 cells proliferated more slowly than wild-type or ZnF6 cells, with ZnF6-7-8 showing the most pronounced growth impairment, consistent with its strongest dimerization defect *in vitro* (Figure 4b, Supplementary Table 1). *Ctcfi* mRNA levels were modestly but significantly increased in ZnF7-8 and ZnF6-7-8 cells, whereas nuclear CTCF protein levels remained largely unchanged across genotypes (Figure 4c, Extended Data Figure 8b, c). In contrast, the chromatin-bound fraction of CTCF was reduced in all mutants (Extended Data Figure 8b, c). Given that these ZnF substitutions did not measurably alter intrinsic DNA binding *in vitro* (Extended Data Figure 5c), the reduced chromatin association observed in cells may arise through several non-mutually exclusive mechanisms, including loss of dimerization-dependent stabilization of CTCF on chromatin and a consequent reduction in residence time, altered recruitment or retention by chromatin-associated factors, or other features of the native chromatin environment.

To further delineate the functional effects of these mutations, we next quantified germline differentiation efficiency using a well-established *in vitro* induction system that enables derivation of functional germ cells from mESCs^54,57–59^. In this system, mESCs are first differentiated into epiblast-like cells (EpiLCs), corresponding to the post-implantation epiblast *in vivo*, followed by cytokine-driven induction of primordial germ cell-like cells (PGCLCs), including BMP4 stimulation (Figure 4d). Whereas EpiLC induction appeared broadly comparable across genotypes, ZnF7-8 and ZnF6-7-8 cells exhibited a striking defect in PGCLC induction (Figure 4e and Extended Data Figure 8d,e). Collectively, these data indicate that the CTCF dimerization interface is physiologically important, supports proliferation, and is required for robust germ cell differentiation.

### Disruption of the CTCF dimerization interface weakens CTCF-cohesin loops

To investigate how disrupting the CTCF dimerization interface impacts 3D genome organization, we performed high-resolution Micro-C^60^ in wild type and mutant mESCs. We obtained 1.6 to 2.4 billion unique ligations per replicate for a total of 17.3 billion unique ligations across all conditions. Replicates were highly concordant across genotypes (Extended Data Figure 9a, Supplementary Table 2). The genome-wide interaction probability as a function of genomic distance was largely unchanged, consistent with observations that even acute CTCF depletion has minimal effects on global contact scaling^4,61,62^ (Extended Data Figure 9b). Likewise, A/B-compartment profiles did not show overt global changes, although each mutant exhibited mild, genotype-specific differences (Extended Data Figure 9c-e).

In contrast, CTCF-cohesin loops were markedly weakened in ZnF7-8 and ZnF6-7-8 cells, with a more moderate reduction in ZnF6 cells (Figure 4f, g and Extended Data Figure 10a). Notably, loop attenuation in these dimerization-interface mutants was substantial but less severe than that observed upon acute CTCF degradation^62^. This effect was specific to extrusion-mediated CTCF-cohesin loops. Affinity-based interactions such as promoter-promoter and enhancer-promoter loops were only weakly affected (Figure 4g, Extended Data Figure 10b)^63–65^. Loop weakening was more pronounced in euchromatic regions than in heterochromatic regions (Extended Data Figure 10c). Despite the substantial attenuation of CTCF-cohesin loops, chromatin insulation was largely retained, with only a moderate reduction of intra-domain interactions compared with acute CTCF degradation (Figure 4h, Extended Data Figure 10d). The stronger effect on looping than on insulation matches prior observations that perturbed two key amino acids required for the CTCF-cohesin interaction^35^. These results suggest that cohesin pausing or stalling at CTCF sites may be sufficient to maintain most boundary insulation, whereas stabilized looped conformations are more dependent on an intact CTCF dimerization interface.

### Quantitative dissection of CTCF occupancy and oligomerization effects

Because Western blotting showed that chromatin-bound CTCF was reduced in all dimerization-interface mutants (Extended Data Figure 8b, c), we performed quantitative CTCF and RAD21 ChIP-seq (Methods). Spike-in-normalized CTCF signal was only modestly reduced, remaining at approximately 80% of wild-type levels in ZnF6 cells and approximately 65% of wild-type levels in ZnF7–8 and ZnF6–7– 8 cells (Figure 4i, j). In contrast, RAD21 signal increased in all mutants (Extended Data Figure 11a, b). These measurements were highly reproducible across biological replicates (Extended Data Figure 11c).

To determine whether altered CTCF and RAD21 occupancy could explain the observed loop weakening, we modeled mutant–wild-type changes in loop intensity as a function of occupancy changes at the two loop anchors. At matched CTCF and RAD21 occupancy, predicted loop intensity was substantially reduced, remaining at approximately 40% of wild-type levels in ZnF6 cells and only approximately 10% of wild-type levels in ZnF7–8 and ZnF6–7–8 cells (Figure 4k). Thus, differences in CTCF and RAD21 occupancy do not account for most of the observed loop weakening, supporting a major role for the CTCF dimerization interface in stabilizing CTCF-cohesin loops.

Applying the same occupancy-adjusted analysis to CTCF/RAD21-associated TAD boundaries revealed a more modest effect, with ∼20% boundary weakening in ZnF6 cells and ∼40% weakening in ZnF7–8 and ZnF6–7–8 cells at matched CTCF and RAD21 occupancy (Extended Data Figure 11d, e).

Finally, as an orthogonal means to assess how CTCF occupancy may impact chromatin looping and insulation, we acutely degraded CTCF across a dTAG titration in an established cell-line^62^. Cells were treated for four hours across six dTAG concentrations, generating a gradient of chromatin-associated CTCF levels (Extended Data Figure 12a-c). Each dTAG condition was profiled by Micro-C and spike-in-normalized CTCF and RAD21 ChIP-seq (Extended Data Figure 12d, e). Both ChIP-seq and Western blotting showed dose-dependent loss of CTCF abundance (Extended Data Figure 12c, d), with ChIP-seq showing a stronger loss than Western blotting. High-resolution Micro-C maps comprising approximately 16.9 billion unique contacts across all conditions showed a clear dose-dependent decay of CTCF-cohesin loop signal (Extended Data Figure 12e, f) and progressive weakening of TAD boundaries (Extended Data Figure 12g).

We utilized the dTAG titration to identify a CTCF concentration that approximated the spike-in-normalized CTCF signal in ZnF7-8 and ZnF6-7-8 cells and acknowledge that the use of distinct cell lines precludes a strict one-to-one comparison. CTCF occupancy in ZnF7–8 and ZnF6–7–8 cells was modestly reduced, remaining at approximately 60– 65% of wild-type levels, which fell between the levels observed after treatment with 0.6 nM and 2 nM dTAG (Figure 4i and Extended Data Figure 12d). CTCF-cohesin loops were substantially weaker in the ZnF7-8 and ZnF6-7-8 mutants than in the 2 nM dTAG condition (Figure 4g and Extended Data Figure 12f). In contrast, boundary weakening was comparable to that observed at approximately 2 nM dTAG (Extended Data Figure 10d and 12g). Together, the occupancy-adjusted regression and dose-matched degradation analyses independently support a model in which CTCF dimerization contributes more strongly to the stabilization of CTCF-cohesin loops than to chromatin insulation generated by cohesin stalling.

### Micro-C read-level analyses favor a ‘symmetric dimer stack’ configuration

Because Micro-C is a nucleosome-centric 3D genomics method that largely analyzes ligations between pairs of nucleosomes, read pairs can be leveraged to infer nucleosome positioning and ligation junction geometries (Figure 5a)^61,66,67^. We first examined nucleosome phasing around CTCF motifs. We observed well-phased nucleosomes around CTCF motifs in agreement with prior work^30–32,62^. However, compared to wild-type, all mutants exhibited weakened amplitudes without an apparent shift in nucleosome position (Figure 5b). The effect was strongest in ZnF7-8 and ZnF6-7-8 cells, consistent with their stronger loop-loss phenotype and lower chromatin-bound CTCF fraction. Nevertheless, the amplitude of nucleosome phasing remained more ordered than in datasets following acute CTCF depletion (Extended Data Figure 13a)^62^.

**Figure 5:**
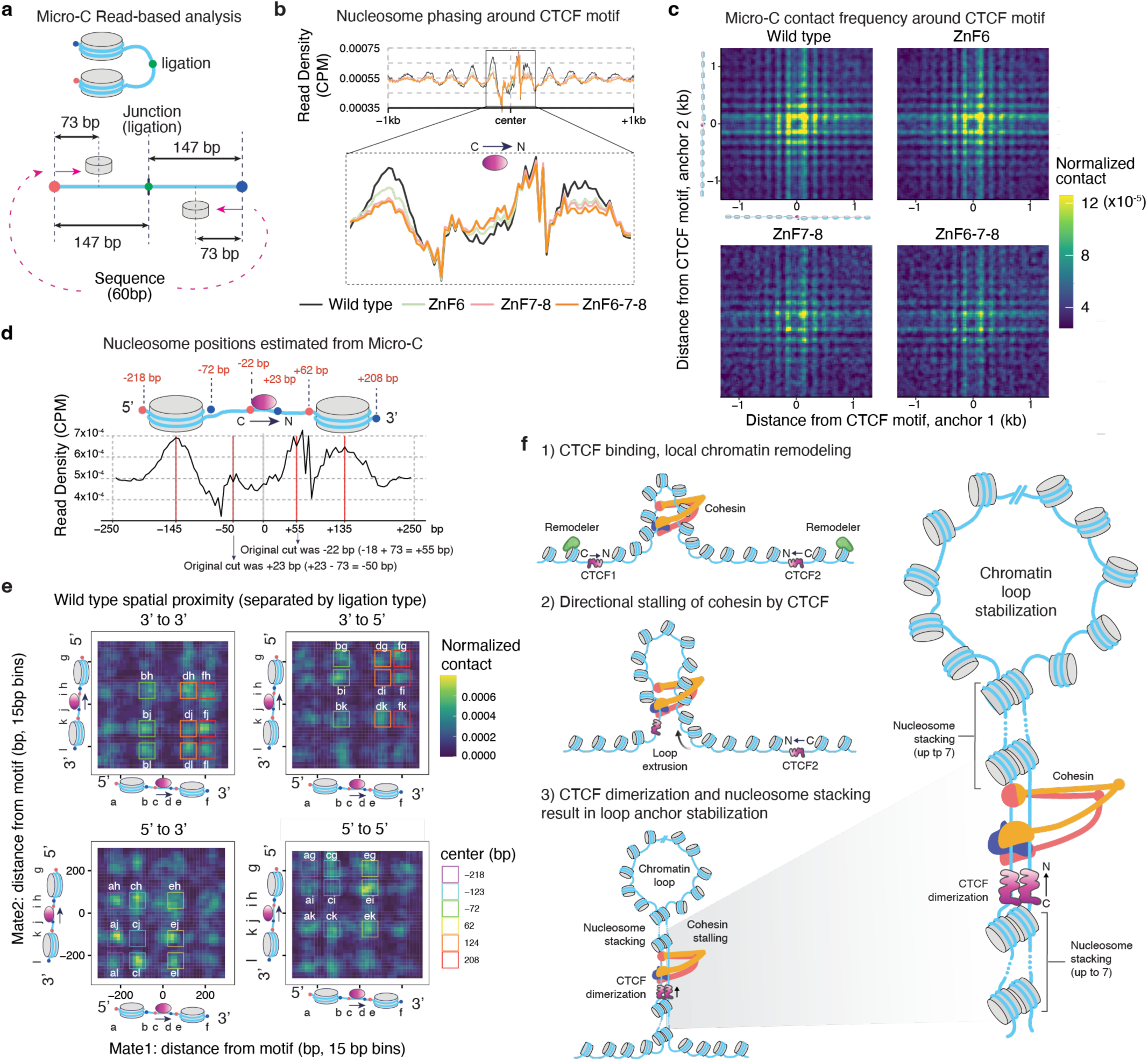
Micro-C read-level signatures around convergent CTCF-cohesin loops support a symmetric ‘dimer stack’ geometry. **a,** Schematic of read-level Micro-C analysis. According to nucleosome-centric ligation, read ends are converted into approximate nucleosome-position or ligation-junction coordinates using strand-aware offsets, enabling inference of ligation geometry around the CTCF motif. **b,** Average nucleosome read density (CPM: counts per million) around plus CTCF motif centers in each condition. The upper panel shows the profile within ±1 kb of the motif center, and the lower panel shows an expanded view of the central ±250-bp region. **c,** Loop-spanning Micro-C contact maps around convergent CTCF–CTCF loops, displayed in inferred nucleosome-dyad coordinates. Micro-C read ends were shifted by 73 bp in a strand-dependent manner to estimate nucleosome dyad positions and oriented relative to the CTCF motif at each anchor. A periodic nucleosome-scale contact pattern extends on both sides of the CTCF motif. Biological replicates were averaged and matrices were smoothed using a 3 × 3-bin window. **d,** Model-guided interpretation of motif-centered signal distribution of nucleosome density track. Top, schematic illustrating coordinate transformations used to relate mapped read positions to ligation junctions. Bottom, average nucleosome density around the CTCF motif center with annotated positions corresponding to the inferred CTCF occupancy footprint. **e,** Wild-type ligation junction interaction maps from **c** separated by ligation-orientation class. Colored boxes denote positional windows used to quantify enrichment consistent with a symmetric nucleosome dimer-stack geometry. **f,** Model for the multi-scale contributions of CTCF dimerization to chromatin organization. Step 1: CTCF binding to DNA bearing its cognate sequence motif recruits chromatin remodelers, establishing phased nucleosome arrays. Step 2: Loop extrusion and directional stalling of cohesin at an individual CTCF site generate chromatin insulation. Step 3: Loop extrusion subsequently brings two CTCF-bound regions together, facilitating CTCF dimerization, which in turn promotes stacking of the flanking nucleosomes in trans to cooperatively stabilize the loop. A zoomed-in view of step 3 is shown on the right.

Because the mutants had less chromatin-bound CTCF, we next asked whether lower CTCF occupancy explained the reduced amplitude. We divided CTCF sites into five groups based on spike-in-normalized CTCF signal and compared genotypes within matched groups (Extended Data Figure 13b, c). Only a modest residual reduction remained, with median phasing scores mostly remaining above 90% of wild-type levels at the first nucleosome on either side of the motif (Extended Data Figure 13d). Thus, most of the reduced nucleosome phasing amplitude is explained by lower CTCF occupancy, with only a modest additional effect of disrupting the dimerization interface.

Given that stabilized CTCF-cohesin loops preferentially form between convergently oriented CTCF motifs^68^, we next asked how Micro-C ligation events are arranged at such loops. We focused on a curated set of convergent CTCF-cohesin loops in which each anchor contains a single, convergently oriented CTCF motif. We then inferred interacting nucleosome positions around motif centers using a strand-aware coordinate transformation (Figure 5a). Wild-type cells displayed an ordered interaction pattern between the two anchors, reflecting ligations between well-positioned nucleosomes on both sides of both CTCF motifs, whereas this pattern appeared less defined in ZnF7-8 and ZnF6-7-8 cells (Figure 5c).

To further interpret these patterns, we examined CTCF binding profiles around the motif center. A model in which CTCF and co-factors occupy DNA approximately from −22 bp to +23 bp around the motif center explained the characteristic pair of peaks (Figure 5d). We then classified ligation events by junction orientation (that is, which ends ligate) and by distance from motif (Extended Data Figure 13e). Read classes expected to arise from the symmetric ‘dimer stack’ geometry that we observed *in vitro* were also enriched in wild-type cells but reduced in ZnF7-8 and ZnF6-7-8 mutants (Figure 5e and Extended Data Figure 13f-i). In addition, contacts between approximately corresponding nucleosome positions on the same relative side of the two CTCF loop anchors were preferentially enriched across the extended phased nucleosome arrays, including after controlling for CTCF occupancy (Figure 5c, Extended Data Figure 14a–c). This broader organization is consistent with the possibility that the extended nucleosome arrays provide additional stabilization of the dimer-stack-associated loop-anchor geometry.

Together, these read-level Micro-C signatures support the notion that the symmetric dimer-stack conformation observed in our cryo-EM reconstructions is more stable or more frequently adopted in wild-type cells, and that disrupting the dimerization interface destabilizes this configuration in cells as well as *in vitro*.

## Discussion

Nucleosomes are highly phased around CTCF-bound sites in cells. This organization is involved in gene insulation and chromatin looping^34,61,66,67,69^. CTCF also has the propensity to self-associate both *in vivo* and *in vitro*^22–25^. However, the specific role of nucleosome phasing around CTCF sites and CTCF self-association in genome organization has remained unclear^15–17,22–24,31^. Here we observe that CTCF acts cooperatively with nucleosomes to form higher-order oligomeric assemblies. The CTCF-CTCF interactions identified here are important for mediating chromatin looping in mammalian cells and are required for cellular differentiation.

Our work makes three important observations related to CTCF binding to chromatinized DNA. First, we observe that CTCF preferentially associates with itself in a parallel fashion when bound to extranucleosomal DNA (Figure 2b,c). Second, CTCF self-association involves an asymmetric binding interface between ZnF6 and ZnF7-8 on parallel CTCF molecules (Figure 2f). Third, we observe that dimerization of CTCF molecules is sensitive to the helical register of CTCF relative to the nucleosome (Extended Data Figure 6). These observations are notable because the parallel orientation of two CTCF sites is required for the appropriate formation of cohesin-driven loop anchors^9,11,19^, where the N-terminus of CTCF must face the extruding cohesin molecule to induce stable pausing of the cohesin motor^18–20^. In cells, these observations may explain why the CTCF dimerization-interface mutants showed robust weakening of loop extrusion-mediated CTCF-cohesin loops, whereas chromatin insulation was largely retained (Figure 4f-h, k, Extended Data Figure 11, 12). This is consistent with prior work mutating residues involved in direct CTCF-cohesin interactions^35^, and insulation has been shown to not require the parallel orientation of CTCF molecules^48,50^. Together, these observations support a model in which cohesin pausing/stalling at CTCF sites is largely sufficient to generate insulation, whereas stabilized loop conformations depend more strongly on an intact CTCF dimerization interface^70^.

Our CTCF dimerization-interface mutants show no DNA binding defects *in vitro* but show various degrees of proliferation and germline differentiation defects in cells that correlate with their inability to support dimerization (Figure 4b, e, Extended Data Figure 5c, 8d, e). Importantly, stabilized CTCF-cohesin loops are only present for a small fraction of time (∼2-3% on average in mESCs^72^) and are relatively short-lived (median ∼10-30 min)^70–72^. Our data therefore suggest that even transient or infrequent looped configurations can be physiologically consequential, potentially by reinforcing regulatory interactions over developmental timescales.

Finally, our read-level Micro-C analyses are consistent with the symmetric ‘dimer stack’ configuration inferred from our structural data and with a broader organization of extended phased nucleosome arrays across CTCF loop anchors (Figure 5c-e, Extended Data Figure 13e, Extended Data Figure 14a-c). Interestingly, this broader array-level organization is reminiscent of recent observations of “fiber homotypy,” in which chromatin fibers with similar nucleosome-spacing patterns preferentially interact in 3D^73^. More broadly, nucleosome spacing has also been shown to tune inter-fiber nucleosome interactions and higher-order chromatin assembly, suggesting that the regular nucleosome spacing organized around CTCF may itself contribute to interactions between opposing loop anchors^74,75^. Although these read-level signatures are modest and Micro-C cannot resolve Ångström-scale interfaces directly, the enrichment of specific motif-proximal ligation patterns in wild-type cells relative to our CTCF dimerization interface mutants suggests that an intact dimerization interface biases local nucleosome-scale assemblies. The effective ligation capture radius of Micro-C has been estimated to be ∼25-42 nm^76,77^. These distance estimates suggest that the Micro-C patterns detected here reflect local nucleosome-scale geometry^73^. Together with recent work indicating that Micro-C and related nucleosome-resolution chromosome-conformation assays can provide readouts of local chromatin geometry^61,66,67,69,78^, our results support the concept that nucleosome-centric ligation patterns can be leveraged to infer aspects of fine-scale chromatin architecture. In the future, integrating ultra-high-resolution 3D genomics with orthogonal structural approaches may enable more precise and quantitative inference of local chromatin structures that have thus far been accessible primarily through structural and biochemical analyses.

Our study provides a model for CTCF and local chromatin structure in facilitating higher-order genomic interactions. First, initial binding of CTCF results primarily in the nucleosome N-terminal to CTCF being remodeled away from the CTCF site to create sufficient space between the nucleosome and CTCF to accommodate extruding cohesin^30,79^. Further nucleosome remodeling forms an array of phased nucleosomes on both sides of CTCF. Loop extrusion by cohesin brings distal, CTCF-bound regions of the genome in proximity. Once two CTCF sites are brought together by cohesin, CTCF self-interactions and stacking interactions between the well-positioned nucleosomes flanking the two CTCF sites act cooperatively in a self-reinforcing mechanism to stabilize chromatin loop anchors, potentially further reinforced by additional interactions between the extended phased nucleosome arrays (Figure 5c, f, Extended Data Figure 13e, Extended Data Figure 14a-c). Together, our work suggests that CTCF function is integrated across multiple genomic scales, spanning from DNA sequence to chromatin organization to influence long-range folding of the genome to ultimately regulate gene expression.

## Acknowledgements

We thank all members of the Vos and Farnung labs for discussions. We thank James Jusuf and Miles Huseyin for helpful discussions, Sarah Nemsick for sharing HEK293T cells, and Domenic Narducci for sharing the CTCF-dTAG mESC line. We thank the Center for Macromolecular Interactions at Harvard Medical School, the Harvard Cryo-EM Center for Structural Biology at Harvard Medical School, the MIT.nano Cryo-EM facility, the MIT Biophysical Instrumentation Facility (BIF), and the HHMI Janelia CryoEM facility staff. We thank Elizabeth Nolan and Wei Hao Lee for their help in performing ICP-mass spectrometry experiments. M.O.V. is a Damon Runyon Philip O’Bryan Montgomery, Jr., MD, Fellow supported by the Damon Runyon Cancer Research Foundation (DRG-2480-22), and a Goldberg Fellow supported by the Cell Biology Education and Fellowship Fund, Harvard Medical School. A.C.S. received support from the Molecular Biophysics training grant (NIH/NIGMS T32 GM008313). M.N. acknowledges support from Daiichi Sankyo Foundation of Life Science Fellowship and HFSP Long-Term Fellowship (LT0022/2024-L). A.S.H. is supported by NIH grants R01EB035127, R01HG014500, and R01CA300848. L.F. is supported by the NIH Director’s New Innovator Award DP2-ES036404. S.M.V. is supported by the Smith Family Awards Program for Excellence in Biomedical Research and the NIH Director’s New Innovator Award 1DP2GM146254-01. S.M.V. and L.F. are Freeman Hrabowski Scholars of the Howard Hughes Medical Institute.

## Author contributions

S.M.V., L.F., M.O.V., and A.C.S. conceived this study with input from all authors; S.M.V., L.F., and A.S.H. supervised and secured funding; B.G.S., S.M.V., and L.C. cloned and purified wild-type and mutant CTCF proteins; B.G.S. and S.M.V. performed and analyzed FA experiments; B.G.S. performed and analyzed BLI experiments; M.O.V. and A.C.S. designed and generated all nucleosomal substrates, performed and analyzed GraFix and crosslinked mass photometry experiments, prepared cryo-EM grids, collected and processed the cryo-EM data, and built and analyzed the structure model; M.O.V. performed the non-crosslinked Fluidix mass photometry; A.C.S. performed the bulk FRET experiments; M.N. generated the genome-edited mESC lines, performed the cell-based assays, Micro-C and ChIP-seq experiments, and analyzed the corresponding data; S.M.V., M.O.V., A.C.S. and M.N. wrote the manuscript with input from all authors; L.F. and A.S.H. edited the manuscript.

## Data availability

Correspondence and requests should be addressed to Seychelle Vos or Lucas Farnung. Materials are available upon request and a completed materials transfer agreement with MIT and Harvard University.

## Materials and Methods

### Cloning and expression of CTCF

The wild-type human CTCF gene was amplified from Addgene plasmid 81789^80^ and inserted into the 438C baculovirus expression vector via ligation-independent cloning^81^. The construct contains an N-terminal 6x His tag followed by maltose-binding protein (MBP) and a tobacco etch virus (TEV) protease cleavage site. CTCF mutants were cloned using round-the-horn mutagenesis. Residues that potentiate CTCF dimerization were identified based on their relative positioning to the Cα backbone since the resolution of our structure did not allow for unambiguous side-chain modeling. ZnF6, carrying the His413Gly and Arg415Ala substitutions; ZnF7-8, carrying the Tyr462Gly, Glu464Val, His477Val, and Glu478Arg substitutions; and ZnF6-7-8, combining all substitutions present in the ZnF6 and ZnF7-8 constructs. All substitutions were designed by aligning the 11 ZnFs of CTCF and substituting the residue with an amino acid found in the same position in a different ZnF with distinct chemical properties or size from the original residue. CTCF ZnF1-11 (residues Lys263-Asp581) construct was cloned into vector 1-C (Addgene plasmid 29654) and was fused to an N-terminal 6x His tag, MBP tag, TEV and human rhinovirus 3C protease sites. Two tryptophans were inserted directly before the CTCF coding region to enable detection of the protein at 280 nm. The CTCF N-terminus (N-term, residues Met1-Pro254) construct was cloned into vector 1-C and was fused to an N-terminal 6x His tag, followed by a MBP tag, and TEV protease site. The CTCF C-terminus (C-term, residues Cys577-Arg727) construct was cloned into a modified version of vector 14-C (Addgene plasmid 48309) containing an N-terminal MBP tag and a TEV protease cleavage site. A tryptophan was inserted at the N-terminus of the construct to permit the measurement of absorption at 280 nm during chromatography. All constructs were verified by full plasmid sequencing.

Wild-type and mutant CTCF were expressed in insect cells. Briefly, purified plasmid DNA (250 ng) was electroporated into DH10EMBacY cells (Geneva Biotech) to generate bacmids. Bacmids were transfected into Sf9 cells grown in ESF921 medium (Expression Systems) with X-tremeGENE9 transfection reagent (Sigma Aldrich) to generate V0 virus. V0 virus was harvested 48–72 hours after transfection. V1 virus was produced by infecting 25 mL of Sf21 cells at 1 million cells/mL with V0 virus. SF21 cells were grown at 27 °C and 300 rpm until 48 hours after proliferation arrest, when V1 virus was harvested and stored at 4 °C. For protein expression, 600 mL of Hi5 cells at 1 million cells/mL were infected with V1 virus and grown for 60-72 hours in ESF921 at 27 °C. Cells were harvested by centrifugation (238 g, 4 °C, 30 min).

CTCF construct ZnF1-11 was expressed in BL21(DE3) RIL cells. Cells were grown in LB medium at 37 °C to OD 600 nm of 0.6. Protein expression was induced by the addition of 0.5 mM isopropyl β-D-1-thiogalactopyranoside (IPTG). Expression was conducted at 18 °C for 16 hrs. Cells were harvested by centrifugation and resuspended in ZnF1-11 lysis buffer (20 mM Tris-HCl pH 7.9, 700 mM NaCl, 25 µM ZnCl2, 30 mM imidazole pH 8.0, 5 mM beta mercaptoethanol (BME), 10% (v/v) glycerol, 0.284 mg/mL leupeptin, 1.37 mg/mL pepstatin A, 0.17 mg/mL PMSF, 0.33 mg/mL benzamidine).

Both the CTCF N-term and C-term constructs were expressed in BL21(DE3) RIL cells. Cells were grown in 2xYT medium at 37° C to OD 600 nm of 0.6. Protein expression was induced by the addition of 0.5 mM isopropyl β-D-1-thiogalactopyranoside (IPTG). Expression was conducted at 37 °C for 3 hrs. Cells were harvested by centrifugation and resuspended in CTCF N-ter/C-ter lysis buffer (20 mM Tris-HCl pH 7.9, 500 mM NaCl, 25 µM ZnCl2, 30 mM imidazole pH 8.0, 5 mM beta mercaptoethanol (BME), 10% (v/v) glycerol, 0.284 mg/mL leupeptin, 1.37 mg/mL pepstatin A, 0.17 mg/mL PMSF, 0.33 mg/mL benzamidine). Harvested cell pellets were snap frozen in liquid nitrogen and stored at -80°.

### Purification of CTCF

All steps were performed at 4 °C unless otherwise noted. Freshly harvested Hi5 cells were resuspended in lysis buffer (500 mM NaCl, 20 mM Tris-HCl pH 7.9, 10% glycerol (v/v), 0.5 mM TCEP pH 7.0, 25 µM ZnCl2, 30 mM imidazole pH 8.0, 0.284 mg/mL leupeptin, 1.37 mg/mL pepstatin A, 0.17 mg/mL PMSF, 0.33 mg/mL benzamidine) and lysed by sonication. Lysates were clarified by centrifugation. Clarified lysates were filtered through a 5 µM syringe filter (Cytiva) followed by 0.45 µM syringe filters and applied to a 5 mL HisTrap column (Cytiva) equilibrated in lysis buffer. HisTrap columns were washed with 10 column volumes (CV) of lysis buffer followed by 5 CV of high salt wash buffer (1000 mM NaCl, 20 mM Tris-HCl pH 7.9, 10% glycerol (v/v), 0.5 mM TCEP pH 7.0, 25 µM ZnCl2, 30 mM imidazole pH 8.0) and 5 CV of lysis buffer. A gradient elution from the HisTrap column onto a 10 mL amylose column equilibrated with lysis buffer (amylose resin from New England Biolabs) was performed using nickel elution buffer (500 mM NaCl, 20 mM Tris-HCl pH 7.9, 10% glycerol (v/v), 0.5 mM TCEP pH 7.0, 25 µM ZnCl2, 500 mM imidazole pH 8.0) from 0-100% over 30 minutes. The HisTrap column was detached and the amylose column was washed with 2 CV of lysis buffer before eluting with amylose elution buffer (500 mM NaCl, 20 mM Tris-HCl pH 7.9, 10% glycerol (v/v), 0.5 mM TCEP pH 7.0, 25 µM ZnCl2, 30 mM imidazole pH 8.0, 116 mM maltose). Peak fractions were analyzed by SDS-PAGE and fractions containing CTCF were pooled. The protein was dialyzed overnight at 4 °C into lysis buffer and incubated with 1.5 mg 6x His-Tobacco Etch Virus protease^82^. Protein was removed from dialysis and applied to a 5 mL HisTrap column to capture uncleaved protein, the 6x His-MBP tag, and TEV protease. Protein was concentrated in a 15 mL Amicon Ultra centrifugal filter (30 MWCO) (Millipore) to 1.0–2.0 mL. The protein was applied to a Superose 6 increase 10/300 GL column (Cytiva) equilibrated in 500 mM NaCl, 20 mM Tris-HCl pH 7.9, 10% (v/v) glycerol, 0.5 mM TCEP pH 7.0, 25 µM ZnCl2. Peak fractions were analyzed by SDS-PAGE. Pure fractions with 260/280 ratios ≤0.7 were concentrated in a 4 mL Amicon Ultra centrifugal filter (30 kDa MWCO) (Millipore), aliquoted, flash frozen in liquid nitrogen, and stored at -80 °C. Inductively coupled plasma–mass spectrometry (ICP–MS) analysis confirmed stoichiometric zinc binding consistent with full occupancy of all 11 ZnFs in the purified wild-type CTCF protein.

Cells expressing ZnF1-11 were lysed by sonication, and the resulting lysate was clarified by centrifugation and applied to a 5 mL HisTrap column (Cytiva) equilibrated in ZnF1-11 lysis buffer. HisTrap column was washed with 10 CV of ZnF1-11 lysis buffer followed by 5 CV of ZnF1-11 high-salt wash buffer (1500 mM NaCl, 20 mM Tris-HCl pH 7.9, 10% glycerol (v/v), 5 mM BME, 25 µM ZnCl2, 30 mM imidazole pH 8.0) and 5 CV of ZnF1-11 lysis buffer. Protein was eluted with ZnF1-11 nickel elution buffer (500 mM NaCl, 20 mM Tris-HCl pH 7.9, 10% glycerol (v/v), 5 mM BME, 25 µM ZnCl2, 500 mM imidazole pH 8.0) onto a 10 mL amylose column equilibrated in ZnF1-11 nickel elution buffer. After 10 CV, the HisTrap column was detached and the amylose column was washed with 2 CV of ZnF1-11 lysis buffer and 10 CV of ZnF1-11 low-salt buffer (300 mM NaCl, 20 mM Tris-HCl pH 7.9, 10% glycerol (v/v), 5 mM BME pH 7.0, 25 µM ZnCl2) before eluting with 5 CV of ZnF1-11 amylose elution buffer (300 mM NaCl, 20 mM Tris-HCl pH 7.9, 10% glycerol (v/v), 5 mM BME, 25 µM ZnCl2, 100 mM maltose) onto tandem 5 mL HiTrap Q HP and HiTrap SP columns (Cytiva) after which the amylose column was removed. The tandem column was washed with 5 CV of ZnF1-11 low-salt buffer. Finally, the HiTrap Q HP column was disconnected, and the protein was eluted from the HiTrap SP column with an NaCl linear gradient from 0.3 to 1.5 M NaCl in 20 mM Tris-HCl pH 7.9, 10% (v/v) glycerol, 5 mM BME, 25 µM ZnCl2. Peak fractions were analyzed by SDS-PAGE, and fractions containing CTCF ZnF1-11 were pooled. The protein was dialyzed overnight at 4 °C into dialysis buffer (500 mM NaCl, 20 mM Tris-HCl pH 7.9, 10% glycerol (v/v), 0.5 mM TCEP, 25 µM ZnCl2) and incubated with 1.0 mg 6x 6xHis 3C protease. Protein was removed from dialysis, diluted to 300 mM NaCl, and applied to a 5 mL HiTrap SP column to remove uncleaved protein. The HiTrap SP column was developed with a NaCl gradient from 0.3 to 1.5 M NaCl in 20 mM Tris-HCl, pH 7.9, 10% glycerol (v/v), 5 mM BME, 25 µM ZnCl2. Fractions containing cleaved protein were pooled, dialyzed in dialysis buffer, concentrated, snap-frozen in liquid nitrogen, and stored at -80 °C.

Frozen cell pellets expressing CTCF N-term or C-term were thawed in a water bath and lysed by sonication on ice. Lysates were clarified by centrifugation. Clarified lysates were filtered through a 5 µM syringe filter (Cytiva) followed by 0.45 µM syringe filters. CTCF N-term lysate was applied to a 5 mL HisTrap column (Cytiva) equilibrated in lysis buffer. The HisTrap column was washed with 10 CV of lysis buffer followed by 5 CV of high salt wash buffer and 5 CV of lysis buffer. A gradient elution from the HisTrap column onto a 10 mL amylose column equilibrated with lysis buffer (amylose resin from New England Biolabs) was performed using nickel elution buffer from 0-100%. After the elution, the HisTrap column was detached, and the amylose column was washed with 2 CV of lysis buffer before eluting with amylose elution buffer. Peak fractions were analyzed by SDS-PAGE and fractions containing CTCF N-term were pooled. CTCF C-term lysate was applied to an amylose column equilibrated in lysis buffer. The amylose column was washed with 10 column volumes (CV) of lysis buffer followed by a wash with high salt buffer and a re-equilibration in lysis buffer. The protein was eluted with amylose elution buffer. Peak fractions were analyzed by SDS-PAGE and fractions containing CTCF C-term were pooled.

CTCF C-ter was applied to a Superose 6 increase 10/300 GL column (Cytiva) and CTCF N-ter was applied to a Superdex 200 10/300 GL column (Cytiva), both equilibrated in 500 mM NaCl, 20 mM Tris-HCl pH 7.9, 10% (v/v) glycerol, 0.5 mM TCEP pH 7.0, 25 µM ZnCl2. Peak fractions were analyzed by SDS-PAGE. Pure fractions with 260/280 ratios ≤0.7 of CTCF N-term and C-term were each concentrated in a 4 mL Amicon Ultra centrifugal filter (10 kDa MWCO) (Millipore), aliquoted, flash frozen in liquid nitrogen, and stored at -80 °C.

### Cloning, expression and purification of histones

Mutant H2B (His46Val, Glu102Lys, His106Leu) was generated using round-the-horn mutagenesis. *X. laevis* histones H2A, H2B, H3, and H4 were expressed in *E. coli* and purified as previously described^83^. *H. sapiens* H4 tailless (Δ1-19) mutant was obtained from The Histone Source (Colorado State University).

### Preparation of DNA constructs for nucleosome reconstitution

DNA templates listed below were cloned into a pIDTSmart-Kan vector (IDT). DNA was prepared using large-scale PCRs as previously described^83^. Forward primers modified with 6-Carboxyfluorescein (6-FAM) and unmodified reverse primers were obtained from IDT. Constructs names correspond to y-W601/W603-CBSx, where y is the length of extranucleosomal DNA, W601/W603 are Widom 601 and 603 sequences, and CBSx is the length of the CTCF binding sequence. Unless specified otherwise, the 80-W601-CBS42 construct was used in most experiments.

### Histone octamer formation and nucleosome reconstitution

*Xenopus laevis* histone octamers containing wild-type or mutant histones were formed as previously described. Nucleosome reconstitution was performed using the salt-gradient dialysis method as previously described^83^. Reconstituted nucleosomes were quantified by measuring absorbance at 280 nm. Molar extinction coefficients for protein and DNA components were calculated and added to yield a molar extinction coefficient for each nucleosome.

### Complex formation for cryo-EM

CTCF-nucleosome complexes were formed by incubating 0.8 µM nucleosome and 1.2 µM CTCF in reaction buffer (20 mM HEPES pH 7.4, 100 mM NaCl, 1 mM MgCl2, 5% glycerol, 0.5 mM TCEP pH 7.0, 100 µM ZnCl2) at 4°C for 1 hour (final volume 250 µL). Reactions were applied to a gradient generated by a Gradient Master mixer (BioComp) lacking or containing 0-0.05% glutaraldehyde and either 10-30% sucrose (for mononucleosomal substrates) or 10-40% sucrose (for dinucleosomal substrates). Samples were centrifuged for 16 hours at 4°C for either 31000 rpm (mononucleosomal substrates) or 28000 rpm (dinucleosomal substrates) in a SW60 rotor (Beckman Coulter). Fractions were collected and analyzed on 3.5% TBE gels prerun for 30 minutes and run for 60 minutes in 0.2x TBE buffer at 150 V, 4 °C. Relevant fractions were pooled and dialyzed at 4°C for 2 hours in reaction buffer lacking glycerol. To validate formation of CTCF-nucleosome dimers or trimers, 2-10 µL of previously crosslinked samples were analyzed using a TwoMP mass photometer (Refeyn). Samples were further dialyzed overnight in fresh reaction buffer lacking glycerol and concentrated to 100 nM.

Quantifoil 2/1 200 Mesh copper grids were glow discharged for 30 seconds at 15 mA using a Pelco Easiglow plasma discharge system. 4 µL of CTCF-nucleosome complex was applied to the grid, blotted after 8 seconds for either 4 seconds (for mononucleosomal substrates) or 3.5 seconds (for dinucleosomal substrates), and vitrified by plunging into liquid ethane using an EM GP2 automatic plunge freezer (Leica Microsystems) operated at 100% humidity and 4°C.

### Cryo-EM data collection and processing of the mononucleosomal sample

Cryo-EM data for the mononucleosomal dimer sample was collected at the Center for Automated Cryogenic Electron Microscopy at MIT. Grids were imaged using a Titan Krios (ThermoFisher) equipped with a BioQuantum K3 direct electron detector (Gatan) operated at 300 kV. Data was acquired automatically using EPU software at a nominal magnification of 105,000x, corresponding to a pixel size of 0.82 Å per pixel. The dataset yielded 20818 movies with 51 frames and total exposure of 50.14 e^-^/Å^2^.

Data processing was performed in cryoSPARC (Structura Bio)^84^. Movies were aligned using patch motion correction followed by patch CTF estimation. Particles were picked using the blob picker, extracted with a box size of 448 pixels and binned 2-fold, yielding 4,105,236 particles. Initial 2D classification, ab-initio reconstruction, heterogeneous refinement and non-uniform refinement were used to select 752,811 particles containing nucleosome dimers. Three rounds of heterogeneous refinement with three copies of the same volume as input were used to enrich for 225,014 particles containing density for CTCF on the extranucleosomal DNA of each nucleosome. Four rounds of 3D classification with a focus mask on both extranucleosomal DNA strands with CTCF were performed to select 50,739 particles where features of CTCF could be visualized in greater detail. Particles were un-binned and subjected to reference-based motion correction followed by another 2D classification to remove remaining junk particles, yielding 44,141 particles. Non-uniform refinement was performed to generate the final Map B with a resolution of 3.7 Å (masked) and 6.1 Å (unmasked) (0.143 gold standard Fourier Shell Correlation (FSC)).

In parallel, two rounds of heterogeneous refinement with three copies of the same volume as input were performed on the original 752,811 particle stack to enrich for 461,738 particles with improved resolution on the stacked nucleosomes. Particles were un-binned and subjected to global and local CTF refinement followed by reference-based motion correction. A non-uniform refinement yielded the final Map A with a resolution of 2.8 Å (masked) and 3.3 Å (unmasked) (0.143 gold standard Fourier Shell Correlation (FSC)) (Extended Data Table 1).

### Cryo-EM data collection and processing of the dinucleosomal dimer sample

Cryo-EM data for the dinucleosomal dimer sample was collected at the Center for Automated Cryogenic Electron Microscopy at MIT. Grids were imaged using a Titan Krios (ThermoFisher) equipped with a BioQuantum K3 direct electron detector (Gatan) operated at 300 kV. Data was acquired automatically using EPU with a nominal magnification of 105,000x and pixel size of 0.83 Å. The dataset yielded 9577 movies with 50 frames and total exposure of 53 e^-^/Å^2^ (Extended Data Table 1). Data processing was performed in cryoSPARC. Movies were aligned using patch motion correction followed by patch CTF estimation. Particles were picked using the blob picker, extracted with a box size of 700 pixels, and binned 2-fold, yielding 1,168,103 particles. Particles were then subjected to 2D classification.

### Cryo-EM data collection and processing of the dinucleosomal trimer sample

Cryo-EM data for the dinucleosomal trimer sample was collected at the Cryo-EM Facility at Janelia Research Campus. Grids were imaged using a Titan Krios (ThermoFisher) with K3 direct electron detector (Gatan) and BioContinuum HD energy filter (Gatan) operated at 300 kV. Data was acquired with SerialEM^85^ at a nominal magnification of 64000x, corresponding to a pixel size of 0.54 Å. The dataset yielded 16008 movies with 50 frames and total exposure of 50 e^-^/Å^2^. Data processing was performed in cryoSPARC. Movie alignment was performed using patch motion correction followed by patch CTF estimation. Particles were picked with the blob picker. 1,944,962 particles were extracted at a box size of 700 pixels, binned by two, and subjected to 2D classification.

### Model building and refinement

An initial AlphaFold3 model of a CTCF-mononucleosome dimer was rigidbody docked into Map B in UCSF ChimeraX with ISOLDE ^86,87^. Individual ZnFs were replaced with the corresponding ZnFs from high-resolution structures of CTCF in complex with cognate DNA (PDB: 8SSS, 8SSQ), with the assumption that CTCF binds to the CTCF motif present in the extranucleosomal DNA in the same manner^15^. Distance restraints for the nucleosomes were generated using a high-resolution *X. laevis* nucleosome (PDB: 3LZ0)^38^ as a template, restraints for the CTCF-bound DNA were generated using a second copy of the model, and local adjustments were made using Maps A and B in ISOLDE version 1.10.1 in UCSF ChimeraX ^86,87^. The model was then subjected to real-space refinement in Phenix version 1.21.2^88,89^ (Extended Data Table 1).

### Mass photometry analysis of GraFix purified complexes

Mass photometry measurements were performed using a TwoMP mass photometer (Refeyn). Selected GraFix fractions were extensively dialyzed in fresh reaction buffer without glycerol (20 mM HEPES pH 7.4, 100 mM NaCl, 1 mM MgCl2, 0.5 mM TCEP pH 7.0, 60-100 µM ZnCl2) at 4°C. Final protein concentrations used for mass photometry measurements ranged from 10 to 50 nM. GraFix crosslinking was essential for mass photometry analysis, as equivalent non-crosslinked samples dissociated upon dilution to nanomolar concentrations required for mass photometry.

Microscope coverslips were cleaned extensively with filtered Milli-Q water and isopropanol prior to use. Measurements were carried out using the standard TwoMP optical configuration, and movies were acquired for 60 s per measurement. For each sample, two independent measurements were typically recorded.

Mass calibration was performed using protein standards (thyroglobulin and β-amylase) under identical buffer conditions, following the manufacturer’s recommended procedures. Data were processed using DiscoverMP software (Refeyn). Mass histograms were generated and analyzed to identify distinct oligomeric species, and reported masses correspond to the centers of fitted peaks.

### Mass photometry of non-crosslinked CTCF-nucleosome complexes

For the non-crosslinked samples, we employed the MassFluidix HC system (Refeyn). The MassFluidix HC system significantly expands the range of sample concentrations amenable to investigation by mass photometry by rapidly diluting the sample up to 10,000-fold on a microfluidic chip (MP-CON-21027, Refeyn), enabling the mass photometry measurement to start within 50 ms from when the molecules begin to dissociate due to dilution effects. Samples were prepared at an initial concentration of 800 nM nucleosomes and 1200 nM CTCF (1:1.5 ratio) in reaction buffer (20 mM HEPES pH 7.4, 100 mM NaCl, 1 mM MgCl2, 0.5 mM TCEP pH 7.0, 100 µM ZnCl2). Each sample was subjected to rapid dilution with a dilution factor of 1:5,000, and movies were acquired for 60 s per measurement.

Mass calibration was performed using protein standards (thyroglobulin and β-amylase) under identical buffer conditions to the static mass photometry conditions described above. Data were processed using DiscoverMP software (Refeyn). Mass histograms were generated and analyzed to identify distinct oligomeric species, and reported masses correspond to the centers of fitted peaks. All samples were measured in duplicate (Supplementary Figure 1).

### Fluorescence anisotropy (FA)

Pre-annealed 42 bp DNA oligos were synthesized by IDT and were dissolved in water to a final concentration of 100 µM. Oligos were aliquoted, snap frozen in liquid nitrogen, and stored at -80 °C prior to use. The following sequences were employed: Random sequence: 5’-6-FAM TTC CAT GTA AAC CTG TCA TAA CTT ACC TGA GAC TAG TTG GAA-3’, Shuffled random sequence: 5’-6-FAM-CGT AGG GAC CGG GGG GAG AAT CAC CGG CTT CGT GGT AGG CCA-3’, Cognate CTCF binding sequence: 5’-6-FAM-TTG CAG TGC CCA CAG AGG CCA GCA GGG GGC GCT AGT GAG GTG. The shuffled random sequence uses the exact same DNA base composition as the cognate CTCF binding sequence. Experiments performed with the CTCF N- and C-termini were conducted with the Cognate CTCF binding sequence, however, the oligos to make the dsDNA were purchased individually from Sigma and annealed in house prior to use.

CTCF was serially diluted in two-fold steps in 20 mM Tris-HCl pH 7.9, 500 mM NaCl, 10% (v/v) glycerol, 0.5 mM TCEP pH 7.0, and 25 µM ZnCl2. DNA was diluted to 25 nM in water (5 nM final concentration). 5 µL of DNA was mixed with 5 µL of CTCF on ice and incubated for 10 min. The DNA-CTCF mixture was diluted with 12.5 µL 2x assay buffer (final conditions 20 mM Tris-HCl pH 7.9, 100 mM NaCl, 10% (v/v) glycerol, 0.5 mM TCEP, 100 µM ZnCl2, 50 µg/mL salmon sperm DNA (Thermo Fisher), 1 mM MgCl2) and 2.5 µL water to a final volume of 25 µL and incubated for 20 min at room temperature in the dark. 18 µL of each solution was transferred to a Greiner 384 Flat Bottom Black Small volume plate. Salmon sperm DNA was omitted from experiments performed with the CTCF N- and C-termini.

Fluorescence anisotropy was measured at room temperature with a Tecan Spark Plate reader (Tecan) with an excitation wavelength of 470 nm (±20 nm), an emission wavelength of 518 nm (±20 nm) and a gain of 125.

### Cy3- and Cy5- H2B(Thr120Cys) labeling

Recombinant human histone H2B(Thr120Cys) was purchased from The Histone Source (Colorado State University). To produce fluorescently Cy3- and Cy5-labeled histones, 2 mg of lyophilized single-cysteine variant H2B(T120C) were dissolved in 1.0 mL of labeling buffer (20 mM Tris-HCl pH 7.0, 6 M guanidine-HCl, 5 mM EDTA), and cysteines were reduced by incubating with 2.5 mM TCEP for 2 h at 25 °C. Cy3- and Cy5-maleimide (APExBIO) were added at a concentration of 3 mM and incubated in the dark at 4 °C overnight. The labeling reactions were quenched with 80 mM beta-mercaptoethanol (BME). Excess of dye was removed by washing the histones with labeling buffer six times in a 4 mL Amicon Ultra 10,000 MWCO concentrator before bringing it to a final volume of 1.0 mL.

To calculate labeling efficiency, dye concentration was quantified using absorbance at 555 nm for Cy3, absorbance at 645 nm for Cy5 and extinction coefficients provided by the manufacturer. Histone concentration was quantified using absorbance at 280 nm and corrected using factors provided by the dye manufacturer. Labeling efficiency was calculated by dividing dye concentration by histone concentration. The measured labeling efficiencies were approximately 60%.

Each fluorescently labeled histone was mixed with equimolar concentrations of unlabeled H2A histone to generate Cy3- and Cy5-labeled H2A/B dimers and dialyzed in high-salt buffer (10 mM Tris-HCl, 2 M NaCl, 1 mM EDTA, 5 mM BME). Labeled H2A/B dimers were purified by gel filtration on a Superdex 75 Increase 10/300 gl column. Fractions containing purified H2A/B dimers were pooled and concentrated, snap-frozen in liquid nitrogen, and stored at -80 °C. The final storage buffer contained 10% (v/v) glycerol.

Cy3- and Cy5-fluorescently labeled nucleosomes were reconstituted by deposition of labeled H2A/B dimers and unlabeled H3/H4 tetramers onto unlabeled DNA by salt gradient dialysis.

### Förster resonance energy transfer (FRET) measurements

Cy3-labeled nucleosome (0.25 µM) and Cy5-labeled nucleosome (0.25 µM, 0.5 µM total nucleosome) were mixed with 0-1.5 µM CTCF wild-type or ZnF6-7-8 mutant in 20 µL reactions in reaction buffer (20 mM HEPES pH 7.4, 100 mM NaCl, 1 mM MgCl2, 5% (v/v) glycerol, 0.5 mM TCEP pH 7.0, 100 µM ZnCl2, 50 µg/mL BSA). Samples were incubated and protected from light at room temperature for 30 minutes, then transferred to a 384-well black flat-bottom small-volume plate (Greiner) and read in a Tecan Spark plate reader. Upon excitation of Cy3 at 510±10 nm, emission scans from 535±10 nm to 720±10 nm with 5 nm step size were performed. Fluorescence intensity values at the 565 nm Cy3 emission maximum (FCy3) and the 660 nm Cy5 emission maximum (FCy5) (Figure 1h and Extended Data Figure 5h) were extracted from the emission scans, averaged, and plotted. FRET ratios were calculated using the equation FCy5/(FCy3+FCy5) (Extended Data Figure 5i, j). Experiments were performed in duplicate.

### Biolayer interferometry (BLI)

Single-stranded DNA oligos were purchased from Sigma Aldrich and annealed to produce biotinylated dsDNA substrates. Sequences are Random sequence: 5’-Biotin-TTC CAT GTA AAC CTG TCA TAA CTT ACC TGA GAC TAG TTG GAA-3’, Cognate CTCF binding sequence: 5’-Biotin-TTG CAG TGC CCA CAG AGG CCA GCA GGG GGC GCT AGT GAG GTG-3’. Oligos were resuspended in water to 100 µM. Complementary ssDNAs were annealed in a final buffer containing 15 mM HEPES, 100 mM potassium acetate. Oligos were heated to 95 °C for 5 minutes, then cooled to 25 °C at a rate of 1 °C/min in a thermocycler.

Octet SA biosensors (Sartorius) coated with streptavidin were incubated in BLI buffer (20 mM Tris-HCl at pH 7.9, 250 mM NaCl, 1 mM MgCl2, 100 µM ZnCl2, 4% (v/v) glycerol, 0.5 mM TCEP pH 7.0, 0.1% (w/v) BSA, 0.05% (v/v) IGEPAL CA-630) for 10 minutes at 30 °C. Throughout the experiment, the plate was shaken at 1000 rpm and incubated at 30 °C. The biosensors were incubated with BLI buffer (250 µL per well) for 60 seconds as the first baseline, then exposed to 250 µL of 50 nM biotinylated linear DNA or a buffer control for 300 seconds. The biosensors were then washed in BLI buffer for 300 seconds. The biosensors were then moved to 250 µL of serially diluted CTCF (400 nM-0 nM in 2-fold steps) to measure CTCF association for 300 seconds. Lastly, the biosensors were returned to BLI buffer to measure CTCF dissociation for 300 seconds. The highest CTCF concentration was also applied to the biosensor with no DNA load as a control for non-specific interactions. Signal from the control biosensor with no DNA loaded and the 0 nM CTCF concentration were subtracted from each association and dissociation curves. All experiments were performed at least three times.

### Analytical density gradient fixation (GraFix)

CTCF-nucleosome complexes were formed by incubating 0.8 µM nucleosome and 1.2 µM CTCF for 30 minutes at 25°C in reaction buffer containing 20 mM HEPES pH 7.4 at 25°C, 100 mM NaCl, 1 mM MgCl2, 5% glycerol, 0.5 mM TCEP, and either 100 µM ZnCl2 (for mononucleosomal substrates) or 60 µM ZnCl2 (for dinucleosomal substrates) with a final volume of 250 µL for mass photometry experiments and 50 µL for all other applications. Reactions were applied to a gradient generated by a Gradient Master mixer (BioComp) containing 0-0.05% glutaraldehyde and either 5-20% sucrose (for linear DNA substrates), 10-30% sucrose (for mononucleosomal substrates) or 15-45% sucrose (for dinucleosomal substrates). Samples were centrifuged for 16 hours at 4°C at 40000 rpm (for linear DNA substrates), 32000 rpm (for mononucleosomal substrates) or 28000 rpm (for dinucleosomal substrates) in a TLS55 rotor (Beckman Coulter). The gradients were fractionated and were analyzed on a 5% polyacrylamide TBE gel (for linear DNA substrates) or 3.5% polyacrylamide TBE gel (for nucleosomal substrates). Gels were prerun for 30 minutes and run for 60 minutes (for linear DNA and mononucleosomal substrates) or run for 90 minutes (for dinucleosomal substrates) in 0.2x TBE buffer at 150 V and 4 °C.

The gels were imaged on an Amersham Typhoon (GE) using the 6-FAM fluorescence at 500 PMT. Full gel scans are provided in Supplementary Figure 2. Densitometry analysis of fluorescent bands was performed using ImageJ software. Briefly, based on migration behavior and mass photometry analysis of the corresponding fractions, bands were assigned as free nucleosome, CTCF-bound nucleosome, and CTCF-nucleosome dimers (2xNucleosomes+2xCTCF). Integrated band intensities were measured using FIJI’s gel analysis function following background subtraction. For nucleosome-only samples (no CTCF added), dimer abundance was calculated relative to the total signal of free nucleosome and dimer bands (2xNucleosome). For samples containing CTCF, dimer abundance was calculated relative to the combined signal of single CTCF-bound nucleosome and dimer (2xNucleosome+2xCTCF) bands. All values were normalized within each lane to control for loading variability. In preparation for mass photometry experiments, relevant fractions were dialyzed twice in reaction buffer without glycerol for at least 2 hours each. 2-10 µL of previously crosslinked samples at 10-50 nM were analyzed using a TwoMP mass photometer (Refeyn).

### Cell culture

The BDF1-2-1 mouse ES cells (mESCs) carrying the *Blimp1-mVenus* and *Stella-ECFP* (BVSC) transgenes^54–56^ were maintained under feeder-free conditions on plates pre-coated with 0.1% sterile gelatin solution (Sigma-Aldrich, cat. no. G1890-100G). Cells were cultured in KnockOut DMEM (Thermo Fisher Scientific cat. no. 10829-018) supplemented with 15% FBS (HyClone, cat. no. SH30396.03, lot AE28209315), 1000 U/mL LIF (Cell Guidance Systems, cat. no. GFM200-1000, lot 0916), 1 mM MEM non-essential amino acid solution (Thermo Fisher Scientific, cat. no. 11140-050), 2 mM GlutaMAX (Thermo Fisher Scientific, cat. no. 35050-061), 100 µg/mL penicillin–streptomycin (Thermo Fisher Scientific cat. no. 15140-122), and 0.1 mM BME (Sigma-Aldrich, M-3148). The medium was further supplemented with 2i, consisting of 10 µM MEK inhibitor (Tocris, cat. no. 4192) and 3 µM GSK inhibitor (Sigma-Aldrich, cat. no. SML1046). mESCs were passaged every two or three days using TrypLE Express (Gibco, cat. no. 12604-021). All cells were cultured at 37 °C with 5% CO2 in air in this study. Cell culture images were captured using an EVOS M5000 microscope (Invitrogen).

### Gene editing, genotyping and sequencing

Genome editing was performed in BDF1-2-1 BVSC mESCs using CRISPR/Cas9 largely according to published procedures, with minor modifications^70^. We co-transfected a Cas9 plasmid (encoding Cas9, puromycin resistance, and the sgRNA) and a repair plasmid (DNA segment to be inserted, typically flanked by 300–500 bp of homology on each side) using Lipofectamine 2000 (Thermo Fisher Scientific, cat. no. 11668019) according to the manufacturer’s protocol, using 1 µg total DNA at a molar ratio of 1:6 of Cas9 vector to repair vector per 5 × 10^4^ cells. sgRNAs were designed using the CRISPOR tool^90^ and are listed in Supplementary Table 3, 4. For each insertion, we designed an individual sgRNA, cloned it into the Cas9 plasmid, co-transfected the repair plasmid and the corresponding sgRNA/Cas9 plasmid, and plated cells onto P10 and P15 dishes at appropriate density. Twenty-four hours after transfection, puromycin was added at a concentration of 1 µg/mL. Following 24 hours of selection, puromycin was removed and cells were cultured for approximately 1 week, after which individual clones were transferred to 96-well plates and expanded. Homozygous edited clones were identified by PCR screening of genomic DNA (primers listed in Supplementary Table 4) using a two-primer strategy, with one primer pair specifically amplifying the edited sequence and the other the unedited one.

To prepare genomic DNA, cells were scraped and resuspended in buffer containing 10 mM Tris-HCl pH 8.0, 1 mM EDTA, 25 mM NaCl, and 200 µg/mL proteinase K (Viagen Biotech, cat. no. 501-PK) and incubated at 60 °C for 1 hour followed by 95 °C for 10 minutes. Following screening, edits were confirmed by sequencing using primers amplifying the entire inserted sequence. Edited clones were subsequently expanded, high-quality genomic DNA was isolated (Zymo Research, cat. no. D3024), and clones were further verified by PCR and sequencing using the same primers as in the initial screen.

### RT-qPCR

Total RNA was extracted from 1 × 10⁵–2 × 10⁵ cells using the RNeasy Micro Kit (QIAGEN, cat. no. 74004) and eluted in 40 µL of RNase-free water, according to the manufacturer’s instructions. RT-qPCR was performed using the Luna Universal One-Step RT-qPCR Kit (New England Biolabs, cat. no. E3005) following the manufacturer’s protocol. The primers used for RT-qPCR are listed in Supplementary Table 4.

### Growth assay

To estimate cell growth rates, wild-type and CTCF mutant cell lines were cultured in parallel. Briefly, cells were maintained as described in the *Cell culture* section. Cell numbers were determined using a Countess 3 automated cell counter (Thermo Fisher Scientific, cat. no. AMQAX2000), and cultures were passaged every 3 days at an appropriate seeding density. Detailed seeding densities and cell counts at each passage are described in Supplementary Table 1.

### Western blot

Western blot was performed as previously described with minor modifications^53,61^. Protein extracts were prepared from 5 × 10^6^ cells using the Subcellular Protein Fractionation Kit for Cultured Cells (Thermo Fisher Scientific, cat. no. 78840) according to the manufacturer’s instructions, assuming a packed cell pellet volume of 20 µL. Protein concentrations were determined using the Qubit Protein Assay Kit (Thermo Fisher Scientific, cat. no. Q33212).

For Western blot analysis, 20 µg of protein extract per lane was mixed with 4× Laemmli sample buffer (Bio-Rad, cat. no. 1610747), boiled at 95 °C for 10 minutes and separated by SDS–PAGE on Novex Tris–Glycine Mini Protein Gels (4–12%, 1.0 mm, WedgeWell format; Thermo Fisher Scientific, cat. no. XP04122BOX) at 200 V for 40 minutes together with Precision Plus Protein Dual Color Standards (Bio-Rad, cat. no. 1610374). Proteins were transferred to membranes (Bio-Rad, cat. no. 1620112) using an eBlot L1 fast wet transfer system (GenScript, cat. no. L00686) with the standard program (16 minutes). After transfer, membranes were blocked for 60 minutes in 2% skimmed milk (Santa Cruz Biotechnology, cat. no. sc-2325) in PBS containing 0.2% Tween-20 (0.2% PBST), and then incubated overnight at 4 °C with the following primary antibodies diluted in the same buffer: rabbit anti-CTCF (1:1,000; Cell Signaling Technology, cat. no. 3418) and mouse anti-TBP (1:3,000; Abcam, clone 1TBP18, cat. no. ab818). Membranes were washed three times for 15 minutes each in 0.2% PBST and incubated for 90 minutes at room temperature with HRP-conjugated secondary antibodies in blocking buffer: goat anti-rabbit IgG (1:5,000; Cytiva, cat. no. NA934-1ML) and sheep anti-mouse IgG (1:5,000; Cytiva, cat. no. NA931-1ML). Signals were detected using Pierce ECL Western Blotting Substrate (Thermo Fisher Scientific, cat. no. 32209) and imaged with a ChemiDoc MP system (Bio-Rad). Band intensities were quantified using ImageJ (v.2.16.0).

### Primordial germ cell-like cell (PGCLC) induction and analysis

PGCLC induction was performed largely as previously described^54,58^. For preconditioning of mESCs, cells were cultured in N2B27 medium supplemented with 0.4 µM PD0325901 (Stemgent, cat. no. 04-2006), 3 µM CHIR99021 (BioVision, cat. no. 4423) and 1,000 U/mL leukemia inhibitory factor (LIF; Merck Millipore, cat. no. ESG1107). Cells were seeded on 12-well plates that had been sequentially coated with 0.01% poly-L-ornithine (Sigma-Aldrich, cat. no. P3655) and 100 ng/mL laminin (BD Biosciences, cat. no. 354232) and maintained under these conditions for 3 days before PGCLC induction.

EpiLC and PGCLC induction was carried out as previously described with minor modifications^54,58^. Briefly, for EpiLC induction, 8 × 10^4^ mouse ES cells were plated per well of a 12-well plate coated with human plasma fibronectin (16.7 µg/mL; Merck Millipore, cat. no. FC010) and cultured in N2B27 medium supplemented with activin A (20 ng/mL; Peprotech, cat. no. 120-14), bFGF (12 ng/mL; Invitrogen, cat. no. 13256029) and 1% v/v KSR (Gibco, cat. no. 10828028).

To generate mPGCLCs, 4 × 10^3^ EpiLCs after 2 days of induction were transferred into each well of a low-cell-binding U-bottom 96-well plate (Thermo Fisher Scientific, cat. no. 174925) and cultured under floating conditions in Glasgow’s minimal essential medium (Thermo Fisher Scientific, cat. no. 11710035) containing 15% KSR, 0.1 mM non-essential amino acids (NEAA; Thermo Fisher Scientific, cat. no. 11140-050), 1 mM sodium pyruvate (Thermo Fisher Scientific, cat. no. 11360-070), 0.1 mM BME (Thermo Fisher Scientific, cat. no. 21985023), 100 U/mL penicillin, 0.1 mg/mL streptomycin (Thermo Fisher Scientific, cat. no. 15140122) and 2 mM L-glutamine (Thermo Fisher Scientific, cat. no. 25030149). The medium was further supplemented with BMP4 (500 ng/mL; R&D Systems, cat. no. 314BP01M), LIF (1,000 U/mL; Merck Millipore, cat. no. ESG1107), SCF (100 ng/mL; R&D Systems, cat. no. 455MC), and EGF (50 ng/mL; R&D Systems, cat. no. 2028EG). Subsequently, aggregates of day 4 mPGCLCs (PGCLCs induced for 4 days) were incubated in TrypLE Express (Gibco, cat. no. 12604-021) for 10 minutes and dissociated by vigorous pipetting, after which BV-positive cells were analyzed using a FACSAria IV cell sorter (BD Biosciences) with FACS Diva v9.0.1 and FlowJo v11.1.0. (Supplementary Figure 4).

### Acute CTCF degradation and dTAG titration

CTCF-dTAG mESCs carrying an endogenous FLAG-SNAPfTag-FKBP12(F36V)-CTCF fusion (cloneB) were described previously^62^. For the graded depletion experiment, cells were treated for 4 h with dTAG-13 (Tocris, cat. no. 6605) at 0, 0.6, 2, 5, 15 or 500 nM. Two independent biological replicates were prepared for each dTAG concentration, with approximately 6.0 × 10⁷ cells collected per condition and replicate. At the end of treatment, matched cultures were harvested in parallel for immunoblotting (5 × 10^6^) (Supplementary Figure 3), quantitative HEK293T spike-in CTCF and RAD21 ChIP-seq (1.2 × 10⁷), and Micro-C (3.5 × 10⁷).

### Quantitative HEK293T spike-in ChIP-seq

Quantitative CTCF and RAD21 ChIP-seq was performed largely according to the spike-in ChIP-seq workflow as described^91^. mESC and HEK293T chromatin were crosslinked, sonicated and quantified separately. Chromatin concentration was estimated by reverse-crosslinking a 20 µL aliquot of each sonicated lysate. A common HEK293T reference was generated by pooling six independently prepared HEK293T chromatin batches and was used for all samples.

For each mESC sample, 15.75 µg mESC chromatin was combined with 10.5 µg pooled HEK293T chromatin, corresponding to 40% HEK293T chromatin by mass, and adjusted to 26.25 µg total chromatin in 700 µL (37.5 ng µL⁻¹). From this mixture, 7.5 µg chromatin in 200 µL was used for CTCF (Cell Signaling Technology, cat. no. #3418) immunoprecipitation and 15 µg chromatin in 400 µL was used for RAD21 (abcam, cat. no. ab992) immunoprecipitation. Immunoprecipitations used 40 µL Protein A Dynabeads (Invitrogen, cat. no. 10001D) per sample. Chromatin was precleared using 5 µL normal rabbit IgG (Cell Signaling Technology, cat. no. 2729S), followed by immunoprecipitation with 5 µL anti-CTCF (Cell Signaling Technology, cat. no. 3418S) or 5 µL anti-RAD21 (Abcam, cat. no. ab992). An aliquot of each mixed chromatin sample was retained as the matched input. Subsequent antibody incubation, bead washes, elution, reverse crosslinking, DNA purification, library preparation and sequencing followed the previously described workflow.

### Spike-in ChIP-seq data processing and normalization

Paired-end reads were adapter-trimmed and aligned with Bowtie2 v2.5.5 to a concatenated mouse(mm10)–human(hg38) reference genome containing species-prefixed chromosomes^92^. Alignments with mapping quality <20 were removed, coordinate-sorted, and PCR duplicates were removed with Picard v3.4.0. Deduplicated alignments were separated by species, and mouse and human read or fragment counts were recorded independently.

For each ChIP and matched input library, a spike-in scale factor was calculated as the number of deduplicated mouse fragments divided by the number of deduplicated human fragments. For each ChIP sample, an input-adjusted spike-in normalization factor was then calculated as the ChIP spike-in scale factor divided by that of the corresponding input library. Mouse-only bigWig tracks used for quantitative analyses were generated from deduplicated mouse BAM files using deepTools v3.5.2 bamCoverage with CPM normalization, 25-bp bins, exclusion of the mm10 ChIP-seq blacklist, and the input-adjusted spike-in normalization factor supplied as the scale factor. For sample-level abundance summaries, an input-adjusted normalization factor was calculated as the ChIP spike-in scale factor divided by the matched input spike-in scale factor. CTCF and RAD21 values in the dTAG series were expressed relative to the target-specific mean of the 0 nM samples, whereas values in the genotype comparison were expressed relative to the target-specific mean of the wild-type samples.

### Peak-level CTCF and RAD21 quantification and ChIP-seq reproducibility

A pooled wild-type mESC CTCF peak^54^ set was used for all peak-level analyses. For each spike-in-normalized CTCF or RAD21 bigWig^93^, signal was summarized across the full width of every reference peak. The mean signal across the complete peak was multiplied by peak width to obtain an integrated peak signal. For genotype-level scatter plots, biological-replicate signals were averaged within each genotype. A single protein-specific pseudocount, defined as one-half of the smallest positive integrated signal observed across all relevant peaks and samples, was added before log₂ transformation. Peak-level comparisons were therefore plotted as log₂(integrated signal + pseudocount).

Reproducibility between individual biological replicates was evaluated from the vectors of log₂-transformed peak signals at the common CTCF peak set. Pairwise Pearson correlation coefficients were calculated, and samples were hierarchically clustered using 1 − Pearson correlation as the distance metric and average linkage. ChIP-seq enrichment at ESC CTCF peaks was calculated independently as the percentage of deduplicated mm10-aligned reads overlapping the CTCF peak set.

### Micro-C library preparation

Micro-C library preparations were performed as described previously^65^. Cells were dissociated with trypsin and then crosslinked in two steps. Cell suspensions were first rotated at room temperature for 35 minutes in 3 mM disuccinimidyl glutarate (DSG; Thermo Fisher Scientific, cat. no. 20593). Formaldehyde (Thermo Fisher Scientific, cat. no. 28908) was then added to a final concentration of 1% v/v and incubation was continued for another 10 minutes. Crosslinking was quenched by adding Tris-HCl (pH 7.5; KD Medical, cat. no. RGE-3370) to 0.375 M. Cells were collected by centrifugation (950 × g, 7 min), washed once with PBS containing 1% (w/v) BSA (Thermo Fisher Scientific, cat. no. 15260037), aliquoted at 5 × 10⁶ cells per tube, and stored at −80 °C until use.

Frozen crosslinked pellets were thawed on ice and resuspended in ice-cold Micro-C buffer MB#1 (50 mM NaCl, 10 mM Tris-HCl pH 7.5, 5 mM MgCl₂, 1 mM CaCl₂, 0.2% NP-40 alternative (Millipore, cat. no. 492018), 1× protease inhibitor cocktail (Sigma, cat. no. 5056489001)). After incubation on ice for 20 minutes, nuclei were pelleted by centrifugation at 1,750 × g for 5 minutes. The nuclei were washed once in the same buffer, and chromatin was digested with 25 U micrococcal nuclease (MNase; Worthington, cat. no. LS004798) at 37 °C for 20 minutes with shaking at 1000 rpm on a heat block. Digestion was stopped by adding EGTA to a final concentration of 4 mM, followed by incubation at 65 °C for 10 minutes with shaking at 1000 rpm to inactivate MNase. Samples were washed twice with MB#2 (50 mM NaCl, 10 mM Tris-HCl pH 7.5, 10 mM MgCl₂, 100 µg/mL BSA (Sigma, cat. no. B8667)).

MNase-digested nuclei were then phosphorylated with T4 polynucleotide kinase (NEB, cat. no. M0201) in NEBuffer 2.1 supplemented with 2 mM ATP and 5 mM DTT at 37 °C for 15 minutes. To generate blunt ends, Klenow fragment (NEB, cat. no. M0210) was added to a final concentration of 0.5 U/µL, and incubation was continued for an additional 15 minutes at 37 °C. End repair was followed by a fill-in reaction using biotin-dATP and biotin-dCTP (Jena Bioscience, cat. no. NU-835-BIO14 and NU-809-BIOX) together with dGTP and dTTP (Jena Bioscience, cat. no. NU1003 and NU1004), each at 66 µM, for 45 minutes at 25 °C. Reactions were terminated by adding EDTA to a final concentration of 30 mM and incubating at 65 °C for 20 minutes. Nuclei were pelleted at 2,000 × g for 5 minutes and washed once with MB#3 (50 mM Tris-HCl pH 7.5, 10 mM MgCl₂).

Proximity ligation was carried out in 1× T4 DNA ligase buffer supplemented with 100 µg/mL BSA using T4 DNA ligase (NEB, cat. no. M0202; 10,000 U per tube) at 25 °C overnight. Residual biotin on unligated DNA ends was removed by treatment with exonuclease III (NEB, cat. no. M0206) at 37 °C for 15 minutes. Crosslinks were reversed by overnight incubation at 65 °C in 1% (w/v) SDS and 250 mM NaCl in the presence of proteinase K (final 2 mg/mL ; Viagen Biotech, cat. no. 501-PK) and RNase A (final 0.1 mg/mL; Thermo Fisher Scientific, cat. no. EN0531).

DNA was purified using a Zymo DNA Clean & Concentrator-25 kit (Zymo Research, cat. no. D4034) and separated on a 1% (w/v) agarose gel (120 V, 60 minutes). The 200–400 bp dinucleosome-sized DNA was excised and recovered using a Zymo gel extraction kit (Zymo Research, cat. no. D4008), and biotinylated ligation products were captured on T1 streptavidin beads (Invitrogen, cat. no. 65601). Sequencing libraries were then prepared directly on bead-bound DNA using the NEBNext Ultra II DNA Library Prep kit (NEB, cat. no. E7645S) according to the manufacturer’s protocol. Libraries were amplified for 8 PCR cycles and purified with a 0.95× volume of AMPure XP beads (Beckman Coulter, cat. no. A63881).

### Micro-C data processing

Micro-C paired-end reads generated on the Illumina NovaSeq platform were processed using a Snakemake v9.1.1 workflow^94^. Raw FASTQ files were aligned to the mouse reference genome (mm10) with bwa-mem2 v.2.2.1^95^ in paired-end mode with the -SP option. The alignments were streamed directly into pairtools v1.1.0 parse2^96^, which converted them to pairs format while annotating mapping quality with the --add-columns mapq option and genomic coordinates based on the mm10 chromosome sizes file with the -c chrom.sizes option, together with the --expand, --report-position outer, and --assembly mouse options. During this step we retained only read pairs with mapping quality ≥30 and an insert size ≤150 bp, using the --min-mapq 30 and --max-insert-size 150 options, and discarded the intermediate SAM output with the --drop-sam option.

For each sequencing lane, mapping statistics were summarized using pairtools v1.1.0 stats, and lane-level pairs files were sorted by genomic coordinates with pairtools v1.1.0 sort. Sorted pairs from all lanes belonging to the same biological replicate were then combined with pairtools v1.1.0 merge with the --max-nmerge 4 option, followed by removal of PCR/optical duplicates with pairtools v1.1.0 dedup with the --max-mismatch 1 and -- mark-dups options. De-duplicated replicate-level pairs were further merged per sample using pairtools v1.1.0 merge, re-sorted with pairtools v1.1.0 sort, and indexed with pairix v0.3.8^97^.

Contact matrices were generated and stored with cooler v0.10.3^98^. Briefly, per-sample sorted pairs were ingested at 500-bp resolution using cooler v0.10.3 cload pairs with the -c1 2, -p1 3, -c2 4, and -p2 5 options together with the mm10 chromosome sizes file, producing single-resolution .cool files. These were subsequently converted into multi-resolution .mcool containers using cooler v0.10.3 zoomify with the --balance option and the following set of resolutions: 10,000,000; 5,000,000; 2,500,000; 1,000,000; 500,000; 250,000; 100,000; 50,000; 25,000; 10,000; 5,000; 2,000; 1,000 and 500 bp. Unless otherwise stated, downstream analyses were performed on the balanced matrices from these .mcool files.

### Micro-C reproducibility analysis

Reproducibility between Micro-C datasets was assessed using the HiCRep.py v0.1.0 stratum-adjusted correlation coefficient (SCC)^99^. Briefly, balanced contact matrices were accessed from mcool files at 50-kb resolution and pairwise SCC values were computed using the hicreppy scc command with the default smoothing parameters. Pairwise SCC values were computed for all sample pairs and summarized as an SCC matrix. The SCC matrix was hierarchically clustered using average linkage on a distance metric of 1 − SCC and visualized as a clustered heatmap with SCC values annotated in each comparison.

### Contact probability decay analysis

Contact probability decay, P(s), defined as the mean contact frequency as a function of genomic separation, was quantified from balanced Micro-C contact maps using cooltools v0.7.1^100^. For each sample, cis contacts at 1-kb resolution were analyzed with the expected_cis module with the smooth and aggregate_smoothed options and smooth_sigma set to 0.1. The resulting distance bins were converted to base-pair units, and separations shorter than 2 kb were excluded from further analysis. Genome-wide decay curves for each sample were obtained from the smoothed and aggregated expectation track. P(s) was plotted on log–log axes, and the local slope P′(s) was computed as the numerical derivative of log P(s) with respect to log genomic distance using the gradient function from numpy v1.26.4^101^.

### Compartment score and saddle analysis

Compartment scores were calculated from Micro-C contact maps using cooltools v0.7.1. Briefly, balanced cis contact matrices at 25-kb resolution were subjected to eigenvector decomposition with the eigs-cis command of cooltools v0.7.1, using the --phasing-track option with a 25-kb GC-content track for mm10 as the phasing track. For each sample, the first eigenvector (E1) at 25-kb resolution was taken as the compartment score track and used for downstream analyses.

Quantile-binned compartment saddle plots were generated from these 25-kb compartment scores using the saddle function of cooltools v0.7.1. For each Micro-C sample, cis expected contact frequencies were computed at 25-kb resolution with the expected_cis function and used for observed/expected normalization in the saddle analysis. Genomic bins were then stratified according to their 25-kb E1 values into 38 equal-occupancy groups between the 2.5th and 97.5th percentiles of the genome-wide E1 distribution, with additional outer bins capturing more extreme values (i.e. qrange = (0.025, 0.975) and n_bins = 38). The saddle function was then used to aggregate and normalize contacts between all pairs of E1 groups, yielding an (n + 2) × (n + 2) matrix of average observed/expected contact frequencies. For visualization and saddle strength quantification, outlier bins were excluded and the inner n × n matrix (38 x 38) was used. The saddle score was defined as the ratio of same-compartment to opposite-compartment contacts using the most negative and most positive E1 bins, (AA + BB)/(AB + BA).

### Insulation score calculation and TAD detection

Insulation profiles and TAD boundary calls were derived from Micro-C contact maps using the insulation functions in cooltools v0.7.1. Briefly, balanced cis contact matrices at 5-kb resolution were loaded from mcool files for each sample, and genome-wide insulation scores were computed with the insulation function in cooltools v0.7.1. Calculations were performed with a 100-kb sliding window, ignoring the two closest diagonals, and bins were considered scorable only when at least two valid pixels were present within the window. TAD boundaries were then identified from the same 5-kb / 100-kb insulation profiles. Boundary calling was carried out with ignore_diags = 2, min_dist_bad_bin = 0, min_frac_valid_pixels = 0.66 and threshold = "Li", and the resulting boundary set was further filtered by applying a boundary strength threshold of 0.3 (i.e. retaining regions with boundary strength > 0.3).

To visualize average contact patterns around TAD boundaries, we performed aggregate “pileup” analyses using the coolpup.py v1.1.0. For each Micro-C map, we used the 5-kb resolution balanced matrices and ran cool-pup.py in local mode with a ±200-kb padding (--local --pad 200000), yielding per-sample local aggregate contact maps centered on TAD boundaries.

In addition, we examined how insulation scores behave around TAD boundaries using deepTools v3.5.2^102^. One-dimensional bigWig tracks of insulation scores at 5-kb resolution (computed with the 100-kb window) were used to the computeMatrix command in reference-point mode. For each sample, insulation scores were aggregated in a 500-kb window centered on the boundary (--referencePoint center, -b 250000, -a 250000, --binSize 5000), while excluding blacklisted regions of the mm10 genome. The resulting matrices were summarized with plotProfile command to generate the files containing average insulation score profiles centered on the TAD boundaries, which were subsequently visualized in R v4.5.0.

### Loop analysis

Chromatin loop sets were generated largely following Jusuf et al. (2025) with minor adaptations^103^. Briefly, loops were called on a mega-merged mouse ESC Micro-C dataset at 1-kb, 2-kb and 5-kb bin sizes using the Mustache loop caller v1.3.3^104^. For each resolution, Mustache was run on autosomes and chrX with a q-value threshold of 0.1, a sigma-zero parameter of 1.6, a maximum genomic separation of 2 Mb, and sparsity thresholds of 0.7 for 1-kb matrices and 0.88 for 2-kb and 5-kb matrices. To compare resolution-specific and shared loops, BEDPE files from the 1-kb, 2-kb and 5-kb calls were intersected pairwise using bedtools v2.30.0 slop with a ±5-kb extension on loop anchors, followed by bedtools v2.30.0 pairToPair to partition loops into categories detected exclusively at one resolution (1-kb only, 2-kb only or 5-kb only), shared between exactly two resolutions (1-kb and 2-kb, 1-kb and 5-kb or 2-kb and 5-kb) or common to all three resolutions^105^. For loops detected at multiple resolutions, coordinates from the finest bin size were kept as the final loop position.

Loop anchors from the union loop set were further classified according to their local chromatin state in mESCs. ChIP–seq peaks for H3K27ac, H3K4me1, H3K4me3, CTCF and RAD21 from a public dataset^54^, as well as promoter intervals defined as ±1 kb around annotated transcription start sites (mm10), were used for classification. Using GenomicRanges v1.60.0^106^, loop anchors were converted to GRanges objects and intersected with each feature set, requiring a minimum overlap of 1 bp. For each anchor we recorded binary overlap flags for promoter, H3K27ac, H3K4me1, H3K4me3, CTCF and RAD21. Based on these overlap profiles, each anchor was assigned both an “inclusive” and a “mutually exclusive” chromatin annotation. In the inclusive scheme, anchors were hierarchically classified as Active_Promoter (overlapping both a promoter interval and H3K4me3), Active_Enhancer (non-promoter anchors overlapping H3K27ac), Poised_Enhancer (non-promoter anchors overlapping H3K4me1) or CTCF_Cohesin (overlapping both CTCF and RAD21), with “Other” reserved for anchors lacking these features. In the exclusive scheme, the same four categories were defined with additional requirements that exclude overlapping categorization (for example, Active_Promoter and enhancer classes were required to be CTCF-negative, and CTCF_Cohesin sites were required to lack H3K27ac, H3K4me1 and H3K4me3), ensuring mutually exclusive assignments.

For loop-level analyses, anchor annotations were collapsed into three broad classes—promoter (P; Active_Promoter), enhancer (E; Active_Enhancer and Poised_Enhancer) and CTCF–cohesin (C; CTCF_Cohesin)— and each loop was assigned to one of four categories (E–P, P–P, C–C, or E– E) according to the combination of annotations at its two anchors. This classification was carried out separately for the inclusive and exclusive annotation schemes, and the resulting category-specific BEDPE files were exported for downstream analyses.

To visualize average contact patterns around loop anchors, we performed loop pileup analysis using cooltools v0.7.1 and coolpup.py^107^. First, cis expected contact frequencies were computed at 2-kb resolution for each Micro-C sample from the balanced mcool matrices using the expected-cis command from cooltools v0.7.1. The resulting genome-wide expected files were used as normalization for all subsequent pileup analyses. Loop pileups were then computed at 2-kb resolution with coolpup.py, using the loop sets described above. For each sample, we aggregated contacts within a ±200-kb window around the loop center (–-flank 200000), restricting the analysis to loops with genomic separations between 10 kb and 2 Mb (–-mindist 10000, –-maxdist 2000000) and using the weight column from the cooler files (–-clr_weight_name weight) together with the precomputed expected contact frequencies (–-expected). Pileups were calculated for all loops combined as well as for loop subsets (E–P, P–P, C–C, or E–E). Subsequently, pileup matrices were center-cropped to a 51 × 51-pixel window (corresponding to ±50 kb at 2-kb resolution) and visualized as observed/expected heatmaps. Aggregate peak enrichment scores were computed from a 1 × 1 central window.

### Loop quantification

Loop strength was quantified directly for individual loops using balanced Micro-C contact maps. For each loop set (inclusive or exclusive, stratified by anchor classes as described above), balanced 1-kb contact matrices were accessed from mcool files using the cooler v0.10.3. For each loop, the sub-matrix spanning the two anchor intervals was fetched from the balanced matrix, and the loop contact intensity was defined as the mean of all pixels.

To assess whether loop strength changes depend on the compartment, loop anchors were additionally annotated by the A/B compartment scores computed as described in the compartment score and saddle analysis section (E: euchromatin (E1 > 0) or H: heterochromatin (E1 < 0)). Loops were then categorized as E–E, H–H or mixed according to the combination of anchor classification. Loop strength differences between genotypes were quantified as log2 ratios of contact intensities.

To compare loop strengths across genotypes and loop classes, per-loop contact intensities were also visualized in log space. For selected loop classes, contact intensities in each condition were plotted against the wild-type condition in log10 space. A small pseudocount (1 × 10^-6^) was added before log transformation to accommodate zero values.

### Nucleosome position analysis around CTCF motif from Micro-C

High-confidence CTCF motif positions in mESCs were defined by intersecting CTCF ChIP–seq peaks^54^ with public CTCF motif datasets^108^. These sites were then used as reference points for nucleosome profiling. Briefly, MACS2 narrowPeak calls from pooled mESC CTCF ChIP–seq were intersected with CTCF motif instances from Jolma, CTCFBSDB, JASPAR MA0139 and SwissRegulon^109^. Motif orientation was assigned separately for each motif source, and peaks for which all sources agreed on orientation and that contained exactly one reduced motif interval (that is, a single interval obtained by merging overlapping motif hits within the peak) were retained as consensus CTCF sites. For each consensus site, the center of the reduced motif interval was computed, and motif centers were exported as separate BED files for plus- and minus-oriented sites. These motif-center BED files were used as reference regions in the nucleosome occupancy analyses.

To derive nucleosome dyad density from Micro-C data, mapped read pairs were converted into dyad positions on the genome. For each Micro-C sample, pairs files containing uniquely mapped “UU” pairs were used. For each read end, a putative nucleosome dyad coordinate was defined by shifting the mapped position by ±73 bp in a strand-specific manner (adding 73 bp for reads on the plus strand and subtracting 73 bp for reads on the minus strand). The resulting dyad positions were written as BED intervals, sorted by genomic coordinate and converted to coverage profiles using bedtools v2.30.0 genomecov to obtain base-pair resolution bedGraph tracks^105^. These bedGraph files were then converted to dyad bigWig files with bedGraph-ToBigWig v2.10 and further normalized to counts per million (CPM) to facilitate comparison across samples.

Average nucleosome occupancy profiles around CTCF motifs were generated with deepTools. For each orientation (plus and minus), motif-center BED files and CPM-normalized dyad bigWigs were supplied to computeMatrix in reference-point mode, using the motif center as the reference point, a 1-kb upstream and 1-kb downstream window (-b 1000, -a 1000) and a bin size of 5 bp. Regions overlapping the mm10 blacklist were masked by providing the blacklist file as an exclusion track, and bins with missing signal were treated as zero (--skipZeros, --missingDataAsZero). The resulting matrices were summarized with plotProfile command to generate the files containing average nucleosome occupancy profiles centered on CTCF motifs for each sample, which were subsequently visualized in R v4.5.0.

### Micro-C analysis following titrated CTCF degradation

Micro-C libraries from the 4 hr CTCF-dTAG titration were prepared and processed as described above, with two independent biological replicates at each dTAG concentration. Replicate-level and pooled contact maps were generated for 0, 0.6, 2, 5, 15 and 500 nM dTAG. For all dTAG titration conditions, we used the same loop sets and TAD boundary set defined for the genotype analyses, and generated loop pileups and quantified loop intensity and boundary strength using the same procedures described above.

### Occupancy-adjusted modeling of CTCF–cohesin loop strength

For occupancy-adjusted loop analyses, only loops with at least one peak from the CTCF peak set overlapping each anchor were retained. Spike-in-normalized CTCF and RAD21 signals were averaged across biological ChIP-seq replicates within each condition. For each loop and condition, the CTCF signal at an anchor was defined as the sum of integrated CTCF signal across all reference CTCF peaks overlapping that anchor. RAD21 anchor signal was quantified over the same CTCF peak intervals. Protein-specific pseudocounts were defined as one-half of the smallest positive anchor signal across the analyzed dataset.

For each CTCF–cohesin loop, spike-in-normalized CTCF and RAD21 signals at the two anchors were log2-transformed with protein-specific pseudocounts and summed across anchors. Loop intensity was log2-transformed after addition of a pseudocount of 1 × 10⁻⁶. For each mutant, paired mutant–wild type changes in loop intensity were modeled by ordinary least squares as a function of the corresponding changes in CTCF and RAD21 occupancy (ΔL ∼ ΔCTCF + ΔRAD21). The occupancy-adjusted loop effect was obtained from the model prediction at ΔCTCF = ΔRAD21 = 0, with 95% confidence intervals. The latter estimates the mutant–wild-type loop change at matched CTCF and RAD21 occupancy; because relatively few loops had both occupancy changes close to zero, this estimate can involve extrapolation. Models were fitted in R v4.5.0 using lm and predict^110^.

### CTCF/RAD21-rich boundary classification and occupancy-adjusted boundary modeling

To identify boundaries most strongly associated with CTCF and cohesin, H3K4me3, H3K27ac, CTCF and RAD21 signals were integrated across each fixed 5-kb boundary bin, log-transformed, standardized across boundaries and subjected to k-means clustering with four centers. The cluster with the highest combined standardized CTCF and RAD21 signal (cluster 3) was designated the CTCF/RAD21-rich boundary class and used for occupancy-adjusted modeling. For each boundary in this class, CTCF peaks overlapping the boundary bin were identified, CTCF signal was defined as the sum of the integrated signals across these peaks, and RAD21 signal was quantified over the same CTCF peak intervals. Mutant and wild-type measurements were then paired at each boundary, and for each mutant, the change in boundary-strength score was modeled as Δboundary strength = β₀ + βCTCFΔCTCF + βRAD21ΔRAD21 + ε, where ΔCTCF and ΔRAD21 represent the corresponding mutant–wild-type changes in log₂-transformed ChIP-seq signal. Separate models were fitted for ZnF6, ZnF7–8 and ZnF6–7–8. Predictions and 95% confidence intervals were calculated at the observed median occupancy changes and at ΔCTCF = ΔRAD21 = 0 using the same R lm/predict framework as for loops.

### CTCF-occupancy-matched nucleosome phasing analysis

To distinguish effects of the dimerization-interface mutations from those caused by lower CTCF occupancy, consensus motif-centered CTCF sites were linked to their corresponding reference CTCF peaks and assigned spike-in-normalized integrated CTCF signal for each genotype.

To compare nucleosome phasing at sites with similar CTCF binding strength across genotypes, CTCF sites were stratified into five binding-strength groups (Q1–Q5) using common quantile boundaries defined from spike-in-normalized CTCF signals pooled across wild-type and all three mutant genotypes. This restricted comparisons within each group to a common range of CTCF signal across genotypes. Plus- and minus-oriented motif sites were analyzed separately. Nucleosome-phasing profiles were compared between genotypes within the same CTCF-strength groups, and the quantitative comparison shown in Extended Data Fig. 13d was performed for plus-oriented Q3–Q5 sites. Q1 and Q2 were excluded from the final phasing-score comparison because the wild-type and mutant CTCF-signal distributions could not be matched sufficiently in these low-signal groups.

For each retained group (Q3–Q5) and Micro-C replicate, CPM-normalized dyad profiles were generated as described above. For the plus-oriented sites used for quantitative comparison, the phasing score was calculated as the sum of dyad CPM values within ±30 bp of the wild-type-defined −1 and +1 nucleosome peaks (−145 and +135 bp), corresponding to windows of −175 to −115 bp and +105 to +165 bp relative to the motif center. Within each CTCF-intensity group, replicate-level scores were normalized to the wild-type median, which was set to 100%. Bars show the genotype median and points denote independent Micro-C replicates.

### Micro-C read-based analysis

Read-level Micro-C analyses were performed for contacts spanning convergent CTCF–cohesin loops. To define motif-centered loop anchors, MACS2 narrowPeak calls for mESC CTCF ChIP–seq were first filtered to peaks overlapping RAD21 narrowPeak calls^54^. CTCF peak summits were then assigned from the corresponding summit BED file; when multiple summits were present within a peak, the summit closest to the peak center was retained. CTCF motif instances were collated from four public motif collections (Jolma 2013, CTCFBSDB, JASPAR MA0139.1 and SwissRegulon)^108^, keeping strand orientation (plus/minus). For each RAD21-filtered CTCF peak, motif hits within ±100 bp of the assigned summit were considered and the motif instance closest to the summit was selected. Motif centers were recorded as 1-bp intervals with the associated orientation.

To identify convergent motif configurations at loop anchors, an exclusive CTCF–cohesin loop set (C–C loops; Defined in “*Loop analysis*” section) was used as input. Motif centers were searched within loop anchors expanded by ±1 kb, and all combinations of left- and right-anchor motifs were evaluated. Loops containing at least one convergent motif configuration (left anchor “plus” and right anchor “minus”) were retained. For downstream read-based filtering, each anchor was refined to a motif-centered “narrow anchor” interval defined as motif center ±600 bp.

Pairs supporting convergent loops were extracted from deduplicated pairtools-formatted.pairs.gz files. Pairs were reformatted as BEDPE and intersected with the narrow-anchor BED using bedtools v2.30.0 pairtobed with the -type both option to require that both read ends overlap anchor intervals^105^. Intersected pairs were further filtered with a custom AWK step to retain only read pairs whose two ends overlapped the two opposite anchors of the same loop (anchor side 1 and anchor side 2), yielding loop-spanning read pairs for downstream analyses.

Filtered loop-spanning read pairs were analyzed in R v4.5.0^110^. For each read end, an approximate ligation-junction coordinate was inferred by shifting the mapped position by 146 bp in a strand-aware manner (read_pos + 146 for plus-strand reads and read_pos − 146 for minus-strand reads). Distances from the corresponding anchor motif center were computed for each read end, and distances were placed in a common orientation by sign-flipping values for minus-oriented motifs. For each readID, oriented distances for anchor 1 and anchor 2 were collected as (dist1, dist2) and restricted to pairs where both distances fell within ±300 bp. Distances were then binned in 15-bp increments and accumulated into 2D contact matrices stratified by ligation-orientation class (3’to3’, 3’to5’, 5’to3’ and 5’to5’). Matrices were globally normalized such that the total signal summed to 1 across all bins and ligation classes and smoothed using a 3 × 3 moving-average filter. Contact enrichment in predefined positional windows (centered on expected ligation coordinates) was quantified as the summed normalized signal within a ±2-bin neighborhood (i.e. a 5 × 5-bin window).

### Extended nucleosome-array contact analysis across CTCF loop anchors

Loop-spanning read pairs were re-extracted for the 3,592 convergent CTCF– cohesin loops with the motif-centered anchors widened to ±2.5 kb, using the filters described above. For each read end a putative nucleosome dyad coordinate was inferred by shifting the mapped position by 73 bp in a strand-specific manner (adding 73 bp for plus-strand reads and subtracting 73 bp for minus-strand reads), distances from the anchor motif center were computed and placed in a common orientation by sign-flipping values for minus-oriented motifs. Wild-type-defined phased nucleosome positions were determined as described above and numbered −7 to −1 and +1 to +7 relative to the CTCF motif; there is no position 0, and the interval between the −1 and +1 nucleosomes contains the motif and remains unassigned. Each inferred dyad was assigned to the nearest such position if it fell within 90 bp of that peak center, giving a 180-bp assignment window per nucleosome; dyads outside these windows were discarded.

Because the two loop anchors are equivalent after orientation normalization, every read pair was entered in both anchor orders in all matrices below. For the dyad-coordinate contact maps, read pairs with both dyad positions between −1,325 and +1,325 bp, the range spanned by the indexed array, were binned in 25-bp increments, smoothed with a 3 × 3 moving-average filter as described above and scaled so that the total signal summed to 1; biological replicates were processed separately and averaged for display. For the nucleosome-indexed analysis, read pairs with both ends assigned were tabulated into a 14 × 14 nucleosome-index contact matrix. For each pair of nucleosome indices, the expected contact frequency was calculated from the product of the independent frequencies of the two nucleosome positions at the two anchors, such that positions contributing more reads were expected to contribute proportionally more contacts. Observed contact frequencies were divided by these expected frequencies to obtain the observed/expected (O/E) ratio, with O/E = 1 indicating the expected contact frequency.

Same-side nucleosome-array enrichment was quantified from the 38 entries in which the two positions lie on the same side of the CTCF motif and their indices differ by no more than one. The (+1, −1) and (−1, +1) entries are excluded, because the −1 and +1 nucleosomes are separated by the CTCF motif rather than by a single nucleosome step. For each biological replicate the O/E values of these 38 entries were averaged with equal weight, enrichment above expectation was taken as this mean minus 1, and enrichment was expressed as a percentage of the corresponding wild-type value within each comparison.

Loop-level CTCF occupancy was defined as the sum of the log2-transformed spike-in-normalized CTCF ChIP–seq signals at the two loop anchors, as in the occupancy-adjusted loop analysis above. Occupancy values from all four genotypes were pooled across these loops, five groups (Q1–Q5) were defined using common quantile boundaries from the pooled distribution, and the same boundaries were applied to every genotype. The metric was calculated independently within Q3, Q4 and Q5, the three groups with sufficient loops and read pairs in all genotypes, and the three estimates were combined with equal weight. Separately, the 182 loops falling in the highest occupancy group in both the ZnF7-8 and ZnF6-7-8 mutants were selected, and this same set of genomic loci was evaluated in all four genotypes, with the matrix, expected frequencies and O/E values recomputed from those loci alone.

## Figure generation

Figures were generated using Adobe Illustrator 2026, GraphPad Prism 11.0.2 (100), and UCSF ChimeraX version 1.11.1

---

**Extended Data Table 1.**
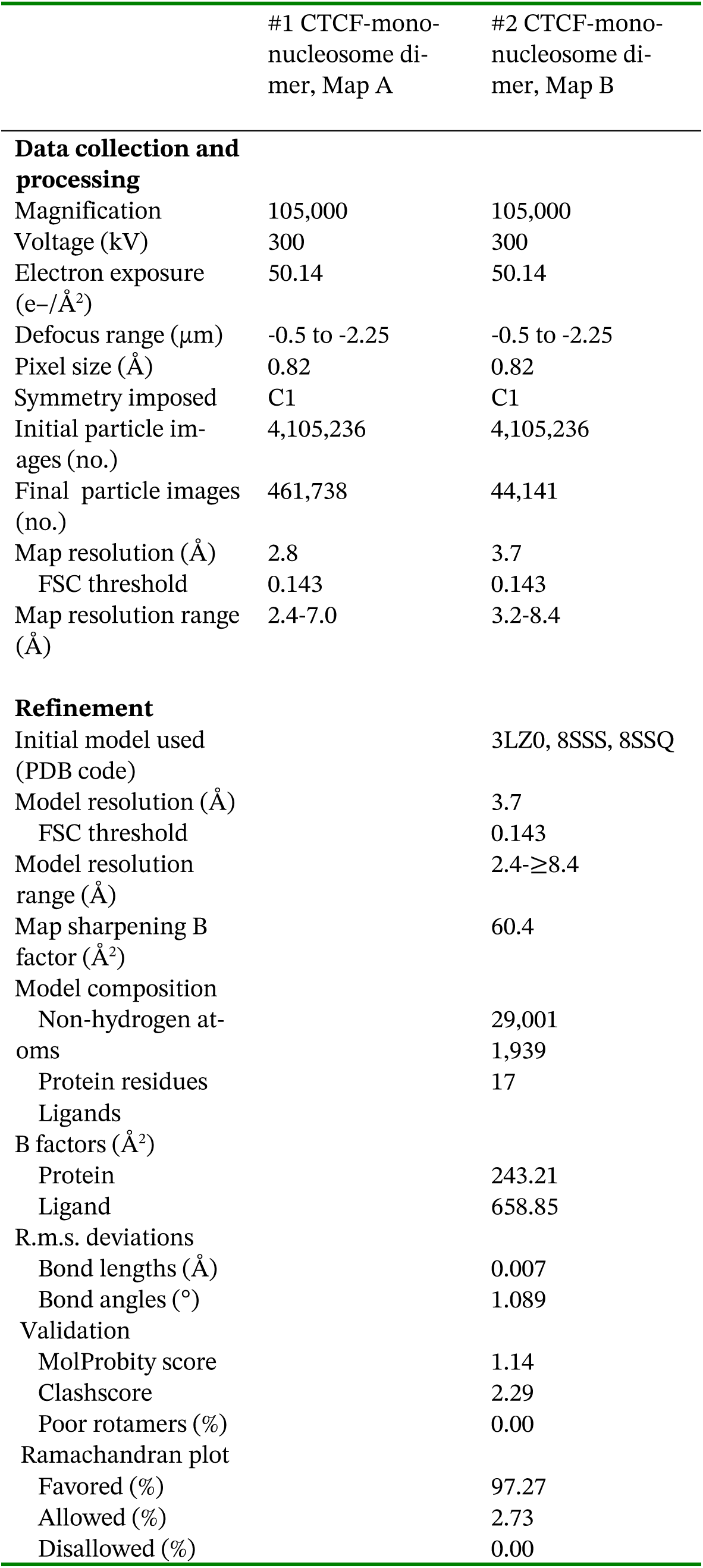
Cryo-EM data collection, refinement, and validation statistics.

## Supplementary Information

**Supplementary Figure 1. Native non-crosslinked mass photometry**

**Supplementary Figure 2. Full gel scans**

**Supplementary Figure 3. Western blot summary**

**Supplementary Figure 4. FACS gating**

**Supplementary Tables: see the separate Excel documents**

**Supplementary Table 1. Growth assay cell counts**

**Supplementary Table 2. Micro-C mapping stats**

**Supplementary Table 3. Oligonucleotides used in this study**

**Supplementary Table 4. CRISPR guide RNA sequences used in this study**

