## Extended Data Figures for "Cooperative CTCF-nucleosome oligomerization stabilizes chromatin loop anchors"

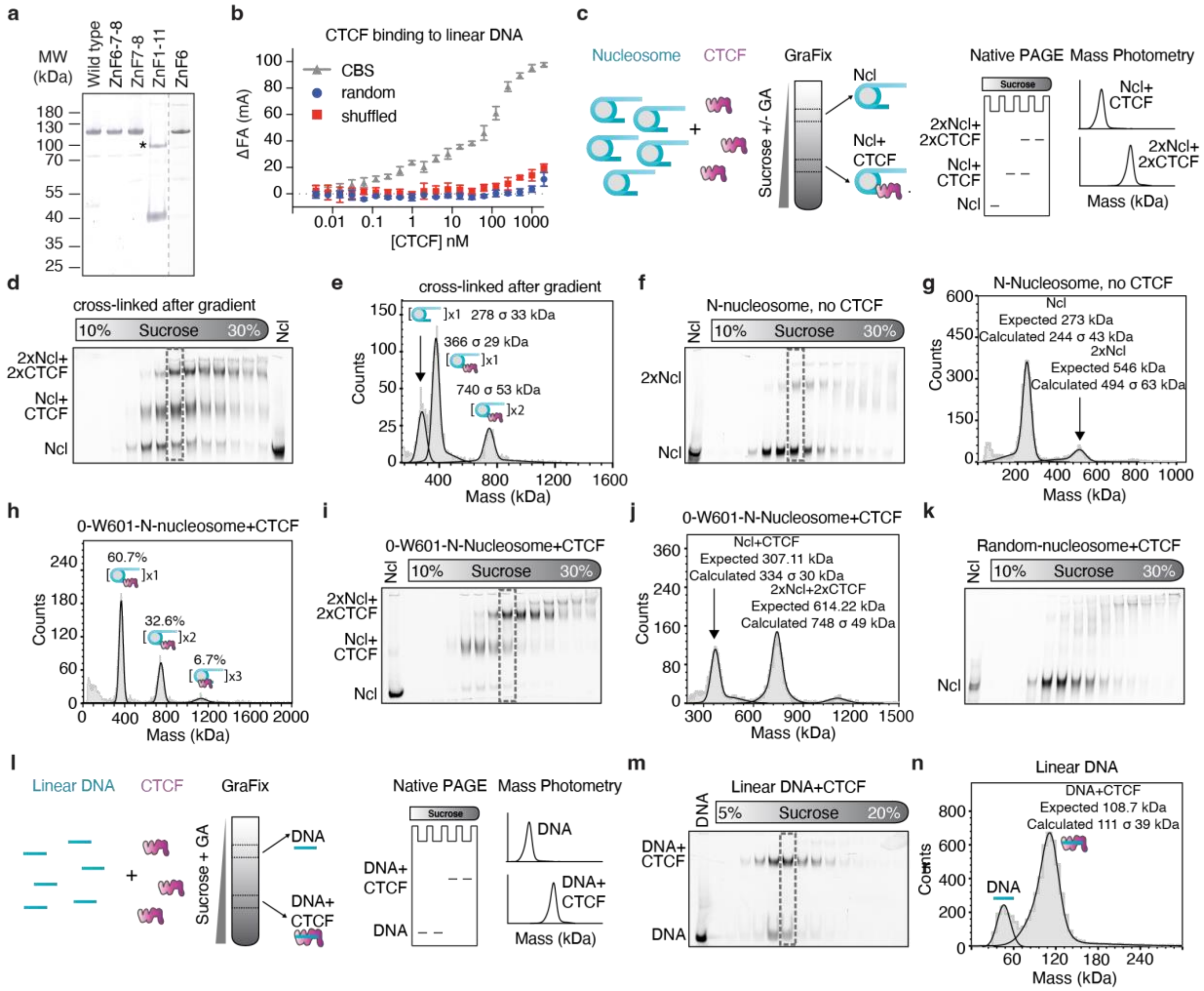


**Extended Data Figure 1: Biochemical and biophysical characterization of CTCF-DNA and CTCF-nucleosome complexes.**

**a,** Coomassie-stained SDS-PAGE analysis of purified wild-type CTCF and CTCF mutants used in this study. Molecular weight markers are shown on the left. The asterisk denotes a minor undigested fraction observed for the CTCF truncated mutant comprising ZnF1 to ZnF11.

**b,** Binding of full-length CTCF to 42 bp linear DNA measured by fluorescence anisotropy (FA). Shown are binding responses to a cognate CTCF binding sequence (CBS), a random DNA sequence, and a shuffled CTCF motif. All measurements were performed in the presence of salmon sperm DNA. Data are shown as mean ± standard deviation from 3 replicates.

**c,** Schematic of the experimental workflow for sucrose gradient centrifugation analysis of nucleosome-CTCF complexes. Samples were mixed and incubated before gradient centrifugation, followed by analysis of gradient fractions by native PAGE and mass photometry to assess oligomeric state.

**d,** Native PAGE analysis of sucrose gradient fractions of FAM-labeled N-nucleosomes-CTCF complexes run in the absence of glutaraldehyde crosslinker. Samples were only crosslinked before being loaded onto the gel. Gradient fractions were resolved on 3.5% acrylamide TBE native gels. Positions corresponding to free nucleosome (Ncl), CTCF-bound nucleosomes (Ncl+CTCF), and CTCF-nucleosome dimers (2×Ncl+2×CTCF) are indicated. A control nucleosome (Ncl) was included in the last lane.

**e,** Mass photometry analysis of N-nucleosome-CTCF complex crosslinked after sucrose gradient centrifugation. The fraction analyzed is highlighted in (**d**). Free nucleosome (Ncl), CTCF-bound nucleosome (Ncl+CTCF), and nucleosome dimer (2×Ncl) species are indicated. Calculated masses of each species are shown.

**f**, Native PAGE analysis of GraFix fractions of the N-nucleosome in the absence of CTCF. The bands corresponding to the N-nucleosome (Ncl) and a faint higher molecular weight band presumably corresponding to a nucleosome dimer (2xNcl) are indicated. The dotted gray box indicates the fraction used in mass photometry analysis.

**g,** Mass photometry analysis of the GraFix fraction highlighted in (**f**), showing populations corresponding to free nucleosome (Ncl) and nucleosome dimers (2×Ncl). Expected and calculated masses are indicated.

**h,** Native, non-crosslinked mass photometry analysis of N-nucleosome lacking the 80 bp extranucleosomal DNA at the entry side (0-W601-N-nucleosome), incubated with CTCF. Distinct populations corresponding to monomeric (x1), dimeric (x2), and trimeric (x3) nucleosome-CTCF species were resolved, with the percentage of each species indicated.

**i,** Native PAGE analysis of GraFix fractions of 0-W601-N-nucleosome incubated with CTCF, showing formation of CTCF-bound nucleosome (Ncl+CTCF) complexes and CTCF-nucleosome dimers (2×Ncl+2×CTCF).

**j**, Mass photometry analysis of the GraFix fraction highlighted in (**i**), showing populations corresponding to CTCF-bound nucleosome complexes (Ncl+CTCF) and CTCF-nucleosome dimers (2×Ncl+2×CTCF). Expected and calculated masses are indicated.

**k,** Native PAGE analysis of GraFix fractions of a FAM-labeled nucleosome containing a 42 bp random DNA sequence incubated with CTCF.

**l,** Schematic of the experimental workflow for GraFix analysis of linear DNA-CTCF complexes. Samples were mixed and incubated prior to sucrose gradient centrifugation with glutaraldehyde, followed by analysis of gradient fractions by native PAGE and mass photometry.

**m,** Native PAGE analysis of GraFix fractions of a 42 bp linear DNA fragment comprising the CTCF-binding motif, incubated with CTCF. Gradient fractions were analyzed on a 5% acrylamide TBE native gel.

**n,** Mass photometry analysis of GraFix fraction highlighted in (**m**) from linear DNA incubated with CTCF, showing formation of DNA-CTCF complexes, with no detectable multimeric species.

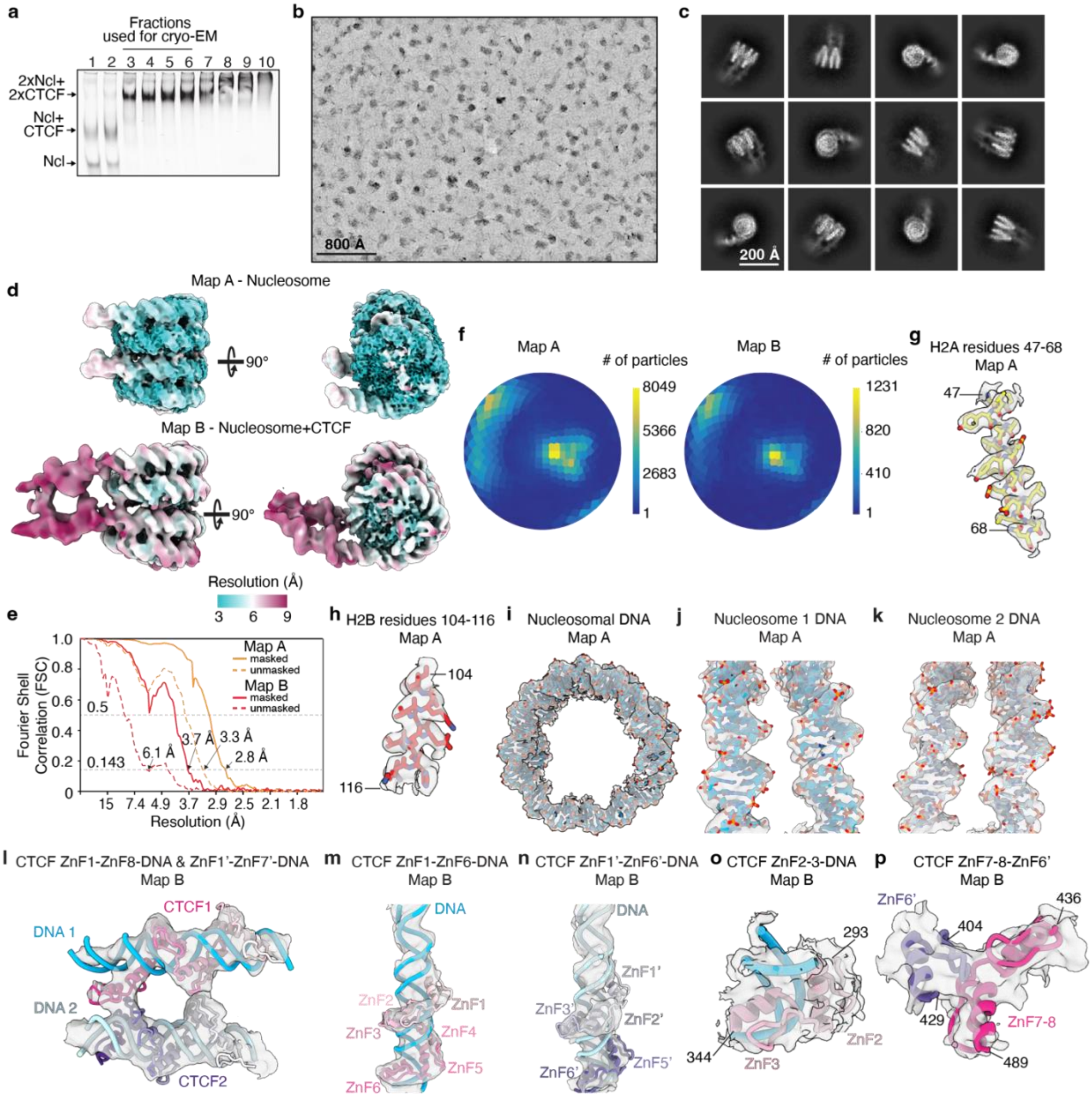


**Extended Data Figure 2. CTCF-mononucleosome dimer complex formation, cryo-EM data collection and map analysis.**

**a,** Native PAGE analysis of GraFix fractions using N-nucleosome and wild-type CTCF. Gradient fractions were resolved on 3.5% acrylamide TBE native gels. Fractions pooled for cryo-EM analysis are indicated.

**b,** Representative micrograph from data collection with scale bar (800 Å).

**c,** Representative 2D classes of the CTCF-nucleosome dimer.

**d,** Maps A and B colored by local resolution.

**e,** Fourier Shell Correlation FSC curves of Maps A and B. The 0.143 and 0.5 FSC thresholds are indicated.

**f,** Angular distribution plots of Map A and B.

**g,h,** Representative fit for model showing histones H2A residues 47-68 (**g**), H2B residues 104-116 (**h**) in map A.

**i,j,k,** Representative fit for model showing nucleosomal DNA of nucleosome 1 (**i**), and zoomed-in views of DNA from Nucleosome 1 (**j**) and Nucleosome 2 (**k**), in density from Map A.

**l,m,n,** Representative fit for model showing CTCF 1 (ZnF1-8) and CTCF 2 (ZnF1'-7') together with their respective bound DNA (DNA 1 and DNA 2) (**l**), zoomed-in views of CTCF 1 ZnF1-6 and corresponding DNA (**m**), and CTCF 2 ZnF1'-6' and corresponding DNA (**n**), in density from Map B. Helical features of the DNA and clear CTCF densities are visible.

**o,p,** Representative fit for model showing CTCF 1 ZnF2-3 (residues 293-344) and corresponding DNA (**o**), and CTCF 1 ZnF7-8 (residues 436-489) interacting with CTCF 2 ZnF6 (residues 404-429) (**p**) in density from Map B.

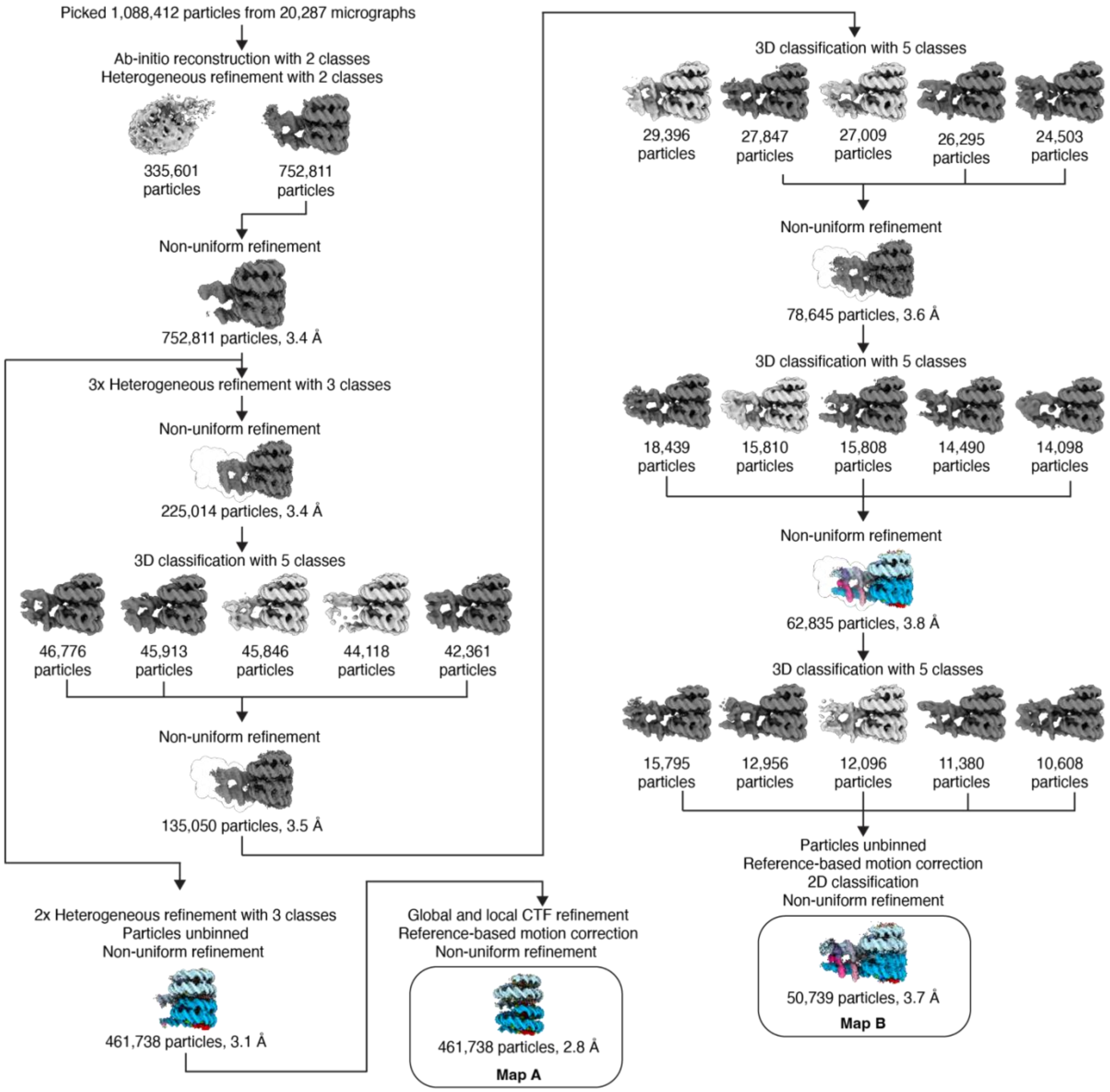


**Extended Data Figure 3. Cryo-EM data processing and reconstruction of CTCF–nucleosome dimer assemblies.**

Cryo-EM image processing workflow for the CTCF-mononucleosome dimer dataset. A total of 1,088,412 particles were picked from 20,287 micrographs and subjected to *ab-initio* reconstruction, heterogeneous refinement, three-dimensional (3D) classification, and non-uniform refinement. Particle numbers and nominal resolutions are indicated for each step. Two independently refined maps (Maps A and B) were obtained.

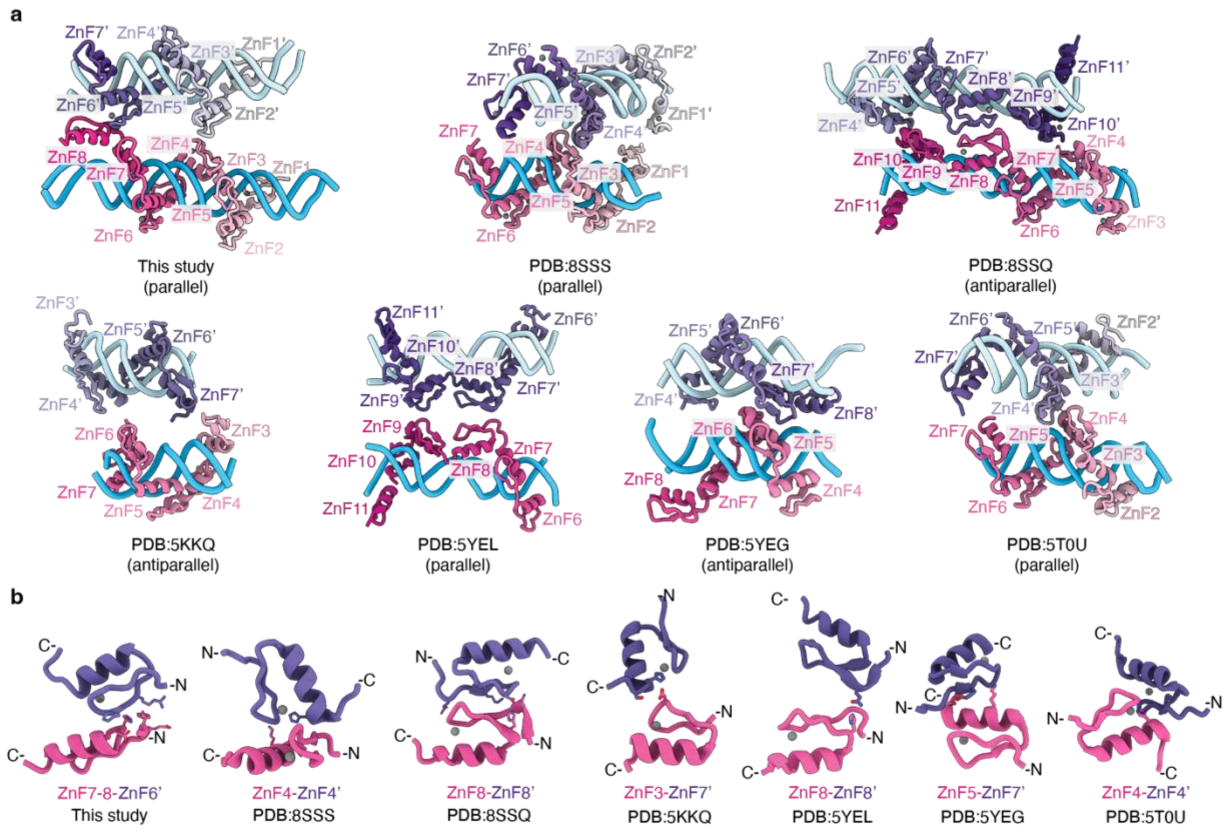


**Extended Data Figure 4. Structural comparison of reported CTCF zinc finger interactions.**

**a,** Structural overview of zinc-finger (ZnF) interactions mediating CTCF dimerization. CTCF zinc finger interactions observed in this study and from published CTCF dimeric structures in complex with cognate DNA. PDBs: 8SSS (ZnF1-ZnF7), 8SSQ (ZnF3-ZnF11), 5KKQ (ZnF3-ZnF7), 5YEL (ZnF6-ZnF11), 5YEG (ZnF4-ZnF8), 5T0U (ZnF2-ZnF7), illustrating variation in ZnF interactions mediating CTCF dimerization.

**b,** Representative views of paired ZnF interfaces comparing interactions observed in this study (ZnF7-8-ZnF6') with those in existing structures shown in (**a**).

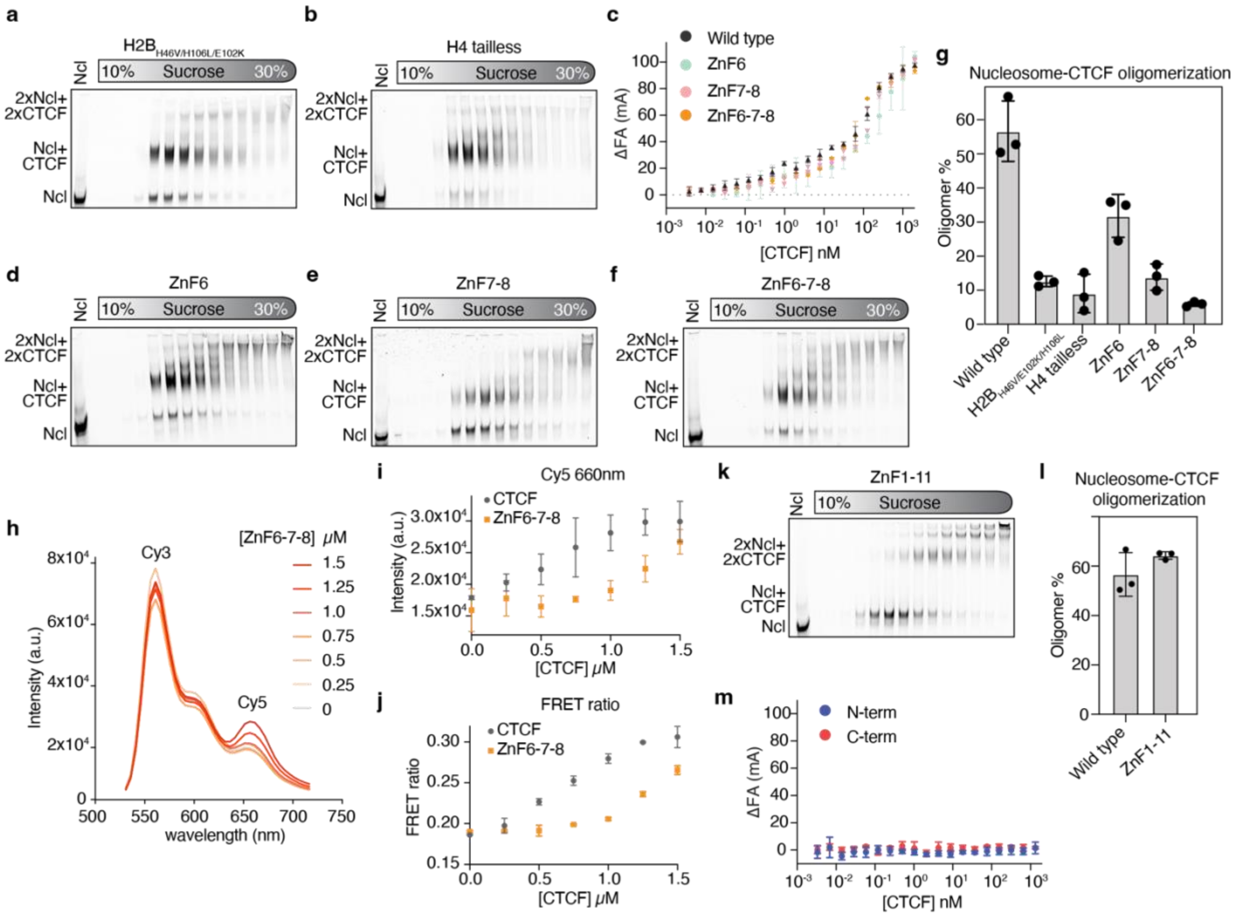


**Extended Data Figure 5. Biochemical characterization of the CTCF ZnF6-ZnF7-8 dimerization interface.**

**a,b,** Native PAGE analysis of GraFix fractions of nucleosome mutants without CTCF. H2B (His46Val, Glu102Lys, His106Leu) (**a**) or H4 tailless (Δ1-19) (**b**). Positions corresponding to free nucleosome (Ncl), CTCF-bound nucleosome (Ncl+CTCF), and CTCF-nucleosome dimers (2×Ncl+2×CTCF) are indicated. A free nucleosome is loaded in the first lane as control.

**c,** Fluorescence anisotropy (FA) analysis of DNA binding by wild-type CTCF and zinc-finger mutants. All measurements were performed in the presence of salmon sperm DNA. Data are shown as mean ± standard deviation from 3 independent measurements.

**d,e,f,** Native PAGE analysis of GraFix fractions of FAM-labeled N-nucleosome incubated with CTCF zinc finger mutants ZnF6 (**d**), ZnF7-8 (**e**), ZnF6-7-8 (**f**). The gels are labeled as in panels (**a,b**).

**g,** Quantification of CTCF-induced nucleosome oligomerization for wild-type CTCF and the indicated CTCF and nucleosome mutants, based on native PAGE analyses shown in (**a,b**), (**d-f**), and **Figure 1i**. Bars represent mean ± standard deviation from 3 independent experiments. Individual data points are shown.

**h,** Fluorescence emission spectra of Cy3/Cy5-labeled nucleosomes upon titration with increasing concentrations of the ZnF6-7-8 mutant (0–1.5 μM), plotted as in **Figure 1g**.

**i,j,** Cy5 emission intensity at 660 nm (**i**) and FRET ratio (**j**) as a function of CTCF concentration, for the ZnF6-7-8 mutant and wild-type CTCF, derived from the spectra shown in (**h**) and **Figure 1g**, respectively. Data are shown as mean ± standard deviation from two independent experiments.

**k,** Native gel analysis of GraFix fractions of N-nucleosome incubated with a CTCF construct comprising only the central 11 zinc finger DNA-binding domain (ZnF1-11). Positions corresponding to free nucleosome (Ncl), CTCF-bound nucleosome (Ncl+CTCF) complexes, and CTCF-nucleosome dimers (2xNcl+2xCTCF) are indicated. Free nucleosome was included in the first lane as a control.

**l,** Quantification of CTCF-dependent nucleosome oligomerization for wild-type CTCF and the ZnF1-11 construct based on native gel analysis shown in (**k**) and **Figure 1i**. Bars represent mean ± standard deviation from 3 independent experiments.

**m,** Fluorescence anisotropy (FA) analysis of isolated N-terminal and C-terminal fragments of CTCF, showing negligible binding over the tested concentration range. No salmon sperm DNA was included in these experiments. Data are shown as mean ± standard deviation.

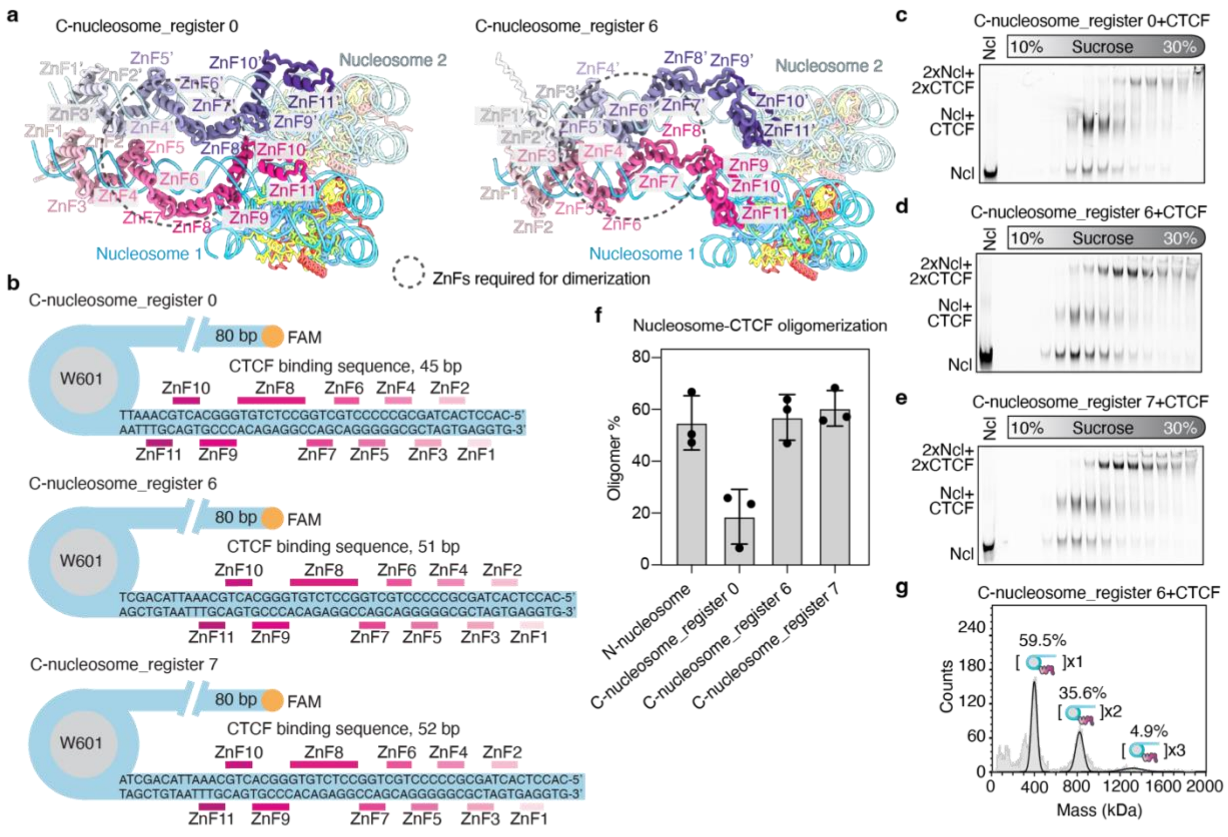

**Extended Data Figure 6. CTCF-induced nucleosome oligomerization is orientation independent but sensitive to the helical register.**

**a,** Predicted structural models of CTCF bound to the C-nucleosome_register 0 and C-nucleosome_register 6 constructs, illustrating the differences in helical register of the CTCF zinc fingers relative to the nucleosome between the two constructs.

**b**, Schematic of reconstituted mononucleosomes containing a CTCF binding sequence to position CTCF with the C-terminus facing the Widom 601 (W601) sequence with 0, 6, and 7 bp between the nucleosome and the CTCF motif (helical registers 0, 6, and 7, respectively). The nucleotide sequence of the extranucleosomal DNA and the relative position of the individual zinc finger contacts (ZnF1-11) are shown. The orange circle represents the FAM fluorophore attached to the nucleosomes for detection.

**c,d,e,** Native PAGE analysis of GraFix fractions of FAM-labeled C-nucleosome_register 0 (**c**), C-nucleosome_register 6 (**d**), and C-nucleosome_register 7 (**e**) incubated with CTCF. Positions corresponding to free nucleosome (Ncl), CTCF-bound nucleosome

(Ncl+CTCF) complexes, and CTCF-nucleosome dimers (2xNcl+2xCTCF) are indicated. Free nucleosome was included in the first lane as a control.

**f**, Quantification of CTCF-induced oligomerization of C-nucleosome substrates based on native PAGE analyses shown in **c-e**. Bars represent mean ± standard deviation from 3 independent experiments. Individual data points are shown.

**g**, Native, non-crosslinked mass photometry analysis of C-nucleosome_register 6 in the presence of CTCF. Distinct populations corresponding to monomeric (x1), dimeric (x2), and trimeric (x3) nucleosome-CTCF species are resolved, with the percentage of each species indicated.

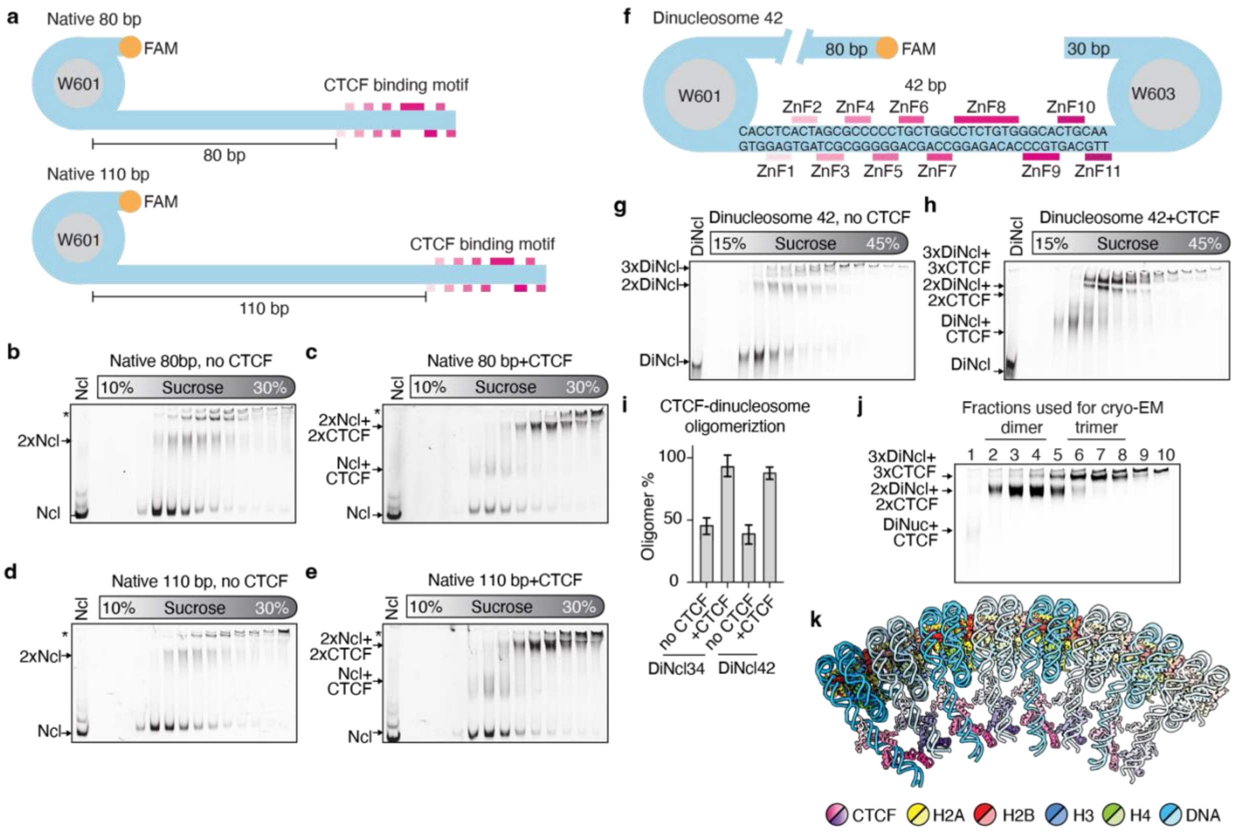


**Extended Data Figure 7. CTCF-dependent oligomerization of native linker-length mononucleosomes and dinucleosomes.**

**a,** Schematic of native linker-length mononucleosome constructs containing 80 bp or 110 bp linker DNA between the Widom 601 (W601) nucleosome positioning sequence and the 42 bp CTCF binding sequence. The position of the CTCF binding motif relative to the nucleosome and linker DNA length is indicated.

**b,c,d,e** Native gel analysis of GraFix fractions of the native 80 nucleosome alone (**b**) and after incubation with CTCF (**c**), and Native 110 nucleosome alone (**d**) and after incubation with CTCF (**e**). Positions corresponding to free nucleosome (Ncl), nucleosome dimers (2xNcl), CTCF-bound nucleosome (Ncl+CTCF) complexes, and CTCF-nucleosome dimers (2xNcl+2xCTCF) are indicated. Asterisks denote higher molecular weight species. Free nucleosome was included in the first lane as a control.

**f,** Schematic of the Dinucleosome 42 containing two Widom positioning sequences (W601 and W603) connected by linker DNA comprising a complete 42 bp CTCF binding motif. The orientation of the CTCF binding motif and the positions of zinc finger contacts (ZnF1-11) are indicated. The linker and extranucleosomal DNA lengths are shown. The orange circle denotes the FAM fluorophore used for detection.

**g,h,** Native gel analysis of GraFix fractions of Dinucleosome 42 in the absence (**g**) or presence of CTCF (**h**). Gradient fractions (15-45% sucrose) were resolved on 3.5% acrylamide TBE native gels. Positions corresponding to free Dinucleosome 42 (DiNcl), Dinucleosome 42 dimers (DiNclx2), CTCF-bound Dinucleosome 42 (DiNcl+CTCF), CTCF-Dinucleosome 42 dimers (2xDiNcl+2xCTCF), and trimers (3xDiNcl+3xCTCF) are indicated.

**i,** Quantifications of CTCF-dinucleosome oligomerization for Dinucleosome 34 and Dinucleosome 42, based on native PAGE analyses in (**g,h**) and in **Figure 3d, e**. Bars represent mean ± standard deviation from 3 independent experiments.

**j,** Native gel analysis of selected fractions collected following complex formation and gradient fixation (GraFix) using Dinucleosome 34 and wild-type CTCF. Fractions pooled for cryo-EM analysis are indicated.

**k,** Extrapolation of the CTCF-mononucleosome dimer model containing multiple interacting CTCF-nucleosome complexes. This is a model and does not necessarily reflect formation of this structure in cells.

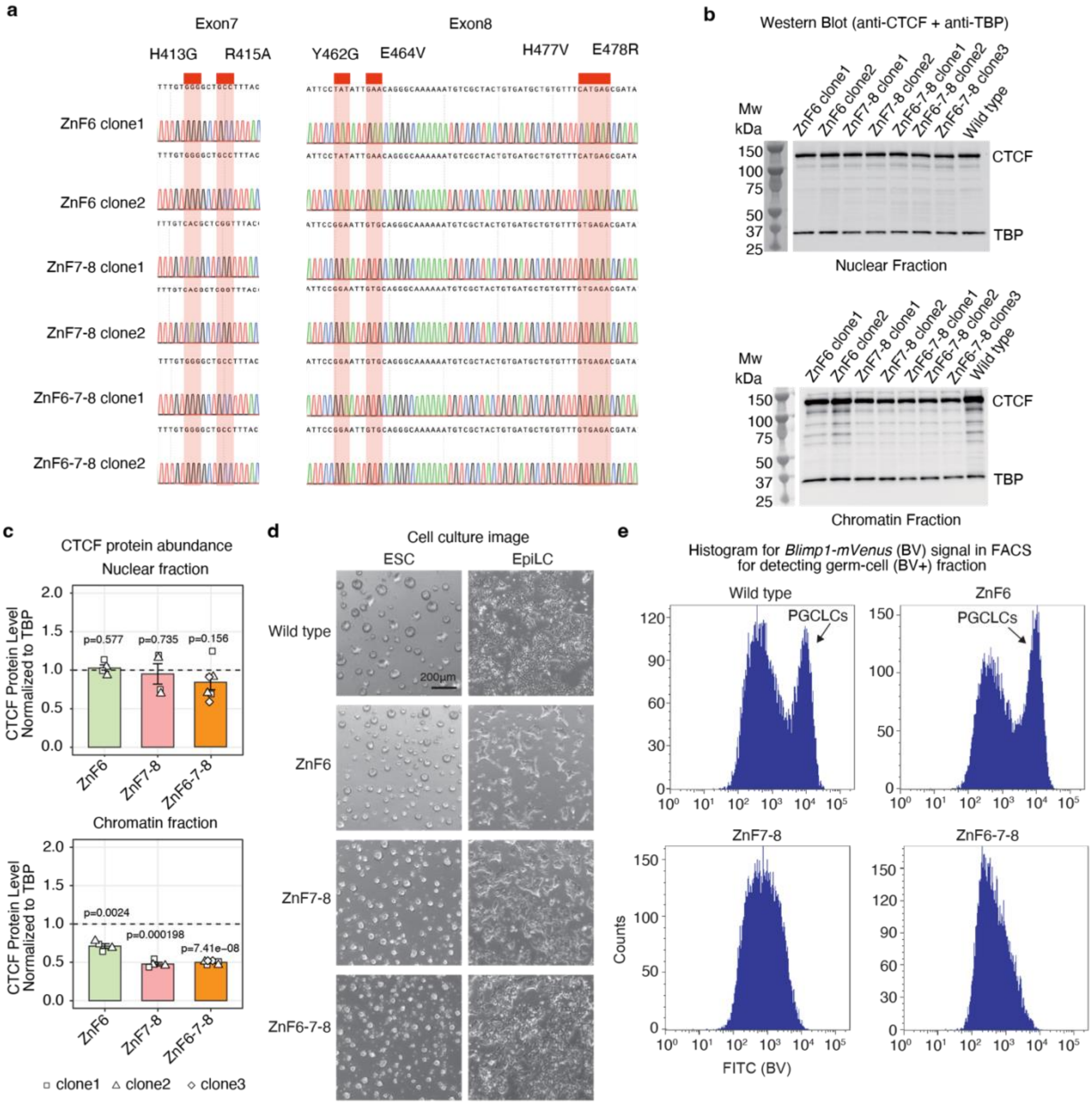


**Extended Data Figure 8. Validation of CTCF dimerization-interface mutants and germline differentiation phenotypes.**

**a,** Sanger sequencing traces confirming the intended point substitutions in exon 7 (ZnF6) and exon 8 (ZnF7-8) regions across independent edited clones.

**b,** Western blots of nuclear and chromatin fractions from wild-type and mutants against CTCF and TBP. TBP is used as a loading control. Samples are indicated.

**c,** Quantification of CTCF abundance from Western blots in **b.**, shown separately for chromatin and nuclear fractions (normalized to TBP; wild-type set to 1). The shapes of points show independent clones; bars represent mean and error bars indicate standard deviation; P values were calculated using two-sided Welch's t-tests.

**d,** Representative brightfield images of wild-type and mutants during mESC culture ("ESC") and after EpiLC induction ("EpiLC"). Scale bar, 200 µm.

**e,** Representative flow cytometry histograms of BV reporter signal (FITC channel) used to quantify BV-positive PGCLCs.

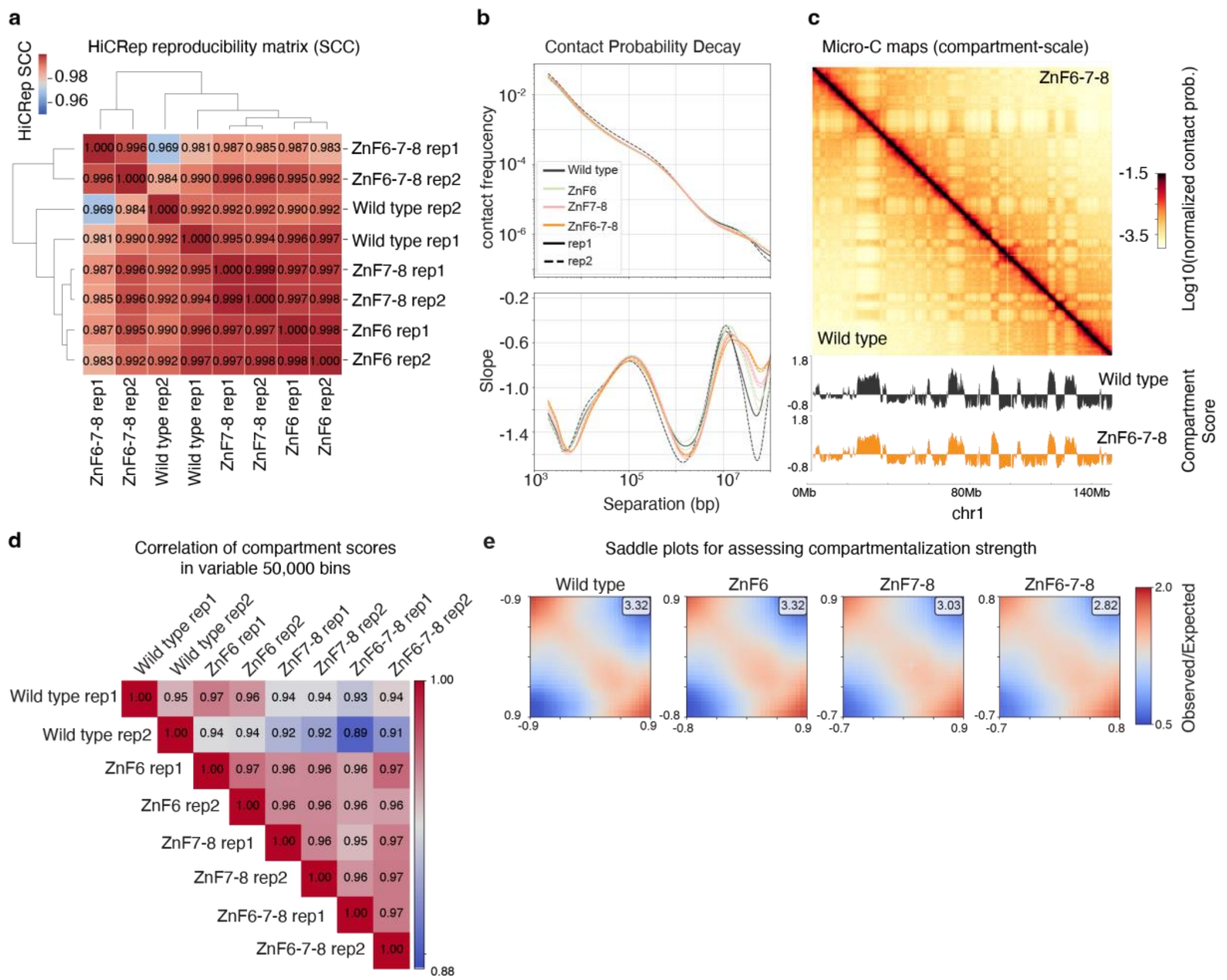


**Extended Data Figure 9. Micro-C reproducibility and compartment-scale genome organization in CTCF dimerization-interface mutants.**

**a,** HiCRep reproducibility matrix (stratum-adjusted correlation coefficient, SCC) across Micro-C replicates and genotypes, with hierarchical clustering.

**b,** Genome-wide contact probability decay P(s) (top) and local slope P′(s) (bottom) as a function of genomic separation for each genotype and replicate.

**c,** Example compartment-scale Micro-C maps on chromosome 1 for wild-type and ZnF6-7-8, together with compartment score tracks shown below.

**d,** Pairwise correlations of compartment scores computed in top variable 50,000 bins across all replicates and genotypes.

**e,** Compartment saddle plots (observed/expected) derived from 25-kb compartment scores for the indicated genotypes. Numbers indicate saddle scores summarizing A/B compartment segregation strength.

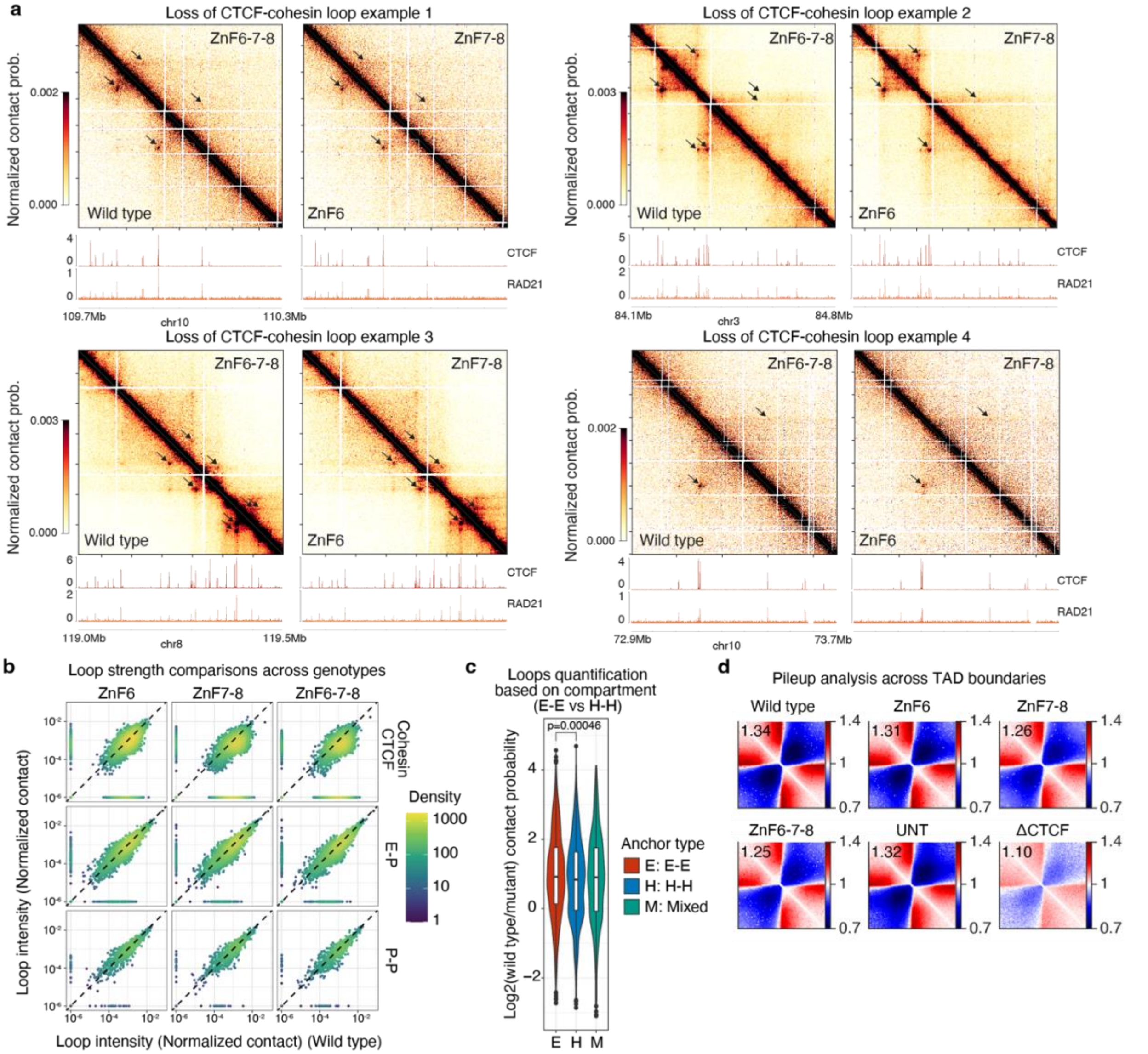


**Extended Data Figure 10. Quantification of CTCF-cohesin loop weakening upon disruption of the CTCF dimerization interface.**

**a,** Representative loci illustrating reduced CTCF-cohesin loop signal across genotypes. Balanced Micro-C contact maps are shown with corresponding CTCF and RAD21 ChIP-seq tracks in wild-type. Black arrows indicate the positions of loops.

**b,** Comparison of loop intensity in each mutant versus wild type for the indicated loop classes (CTCF-cohesin; enhancer-promoter (E-P); promoter-promoter (P-P)). Each point corresponds to one loop; the dashed line indicates y = x; color indicates point density.

**c,** Loop intensity quantification stratified by loop-anchor classification by compartment. Violin plots show distributions of loop intensity enrichment ($Log_2$ wild type/mutant) for CTCF-cohesin loops grouped as euchromatin-euchromatin (E-E), heterochromatin–heterochromatin (H-H), or mixed anchors (Mixed). In box plots, center line denotes the median, lower and upper hinges denote first and third quartiles, respectively, and upper and lower whiskers extend to the largest and lowest values within the 1.5× interquartile range, respectively. P values were calculated using two-sided Welch's t-tests.

**d,** Aggregate contact maps ("pileups") around called TAD boundaries for wild type, mutants, and the samples (untreated (UNT) and ΔCTCF) from Narducci 2025[62]. Numbers indicate intra-domain contact enrichment.

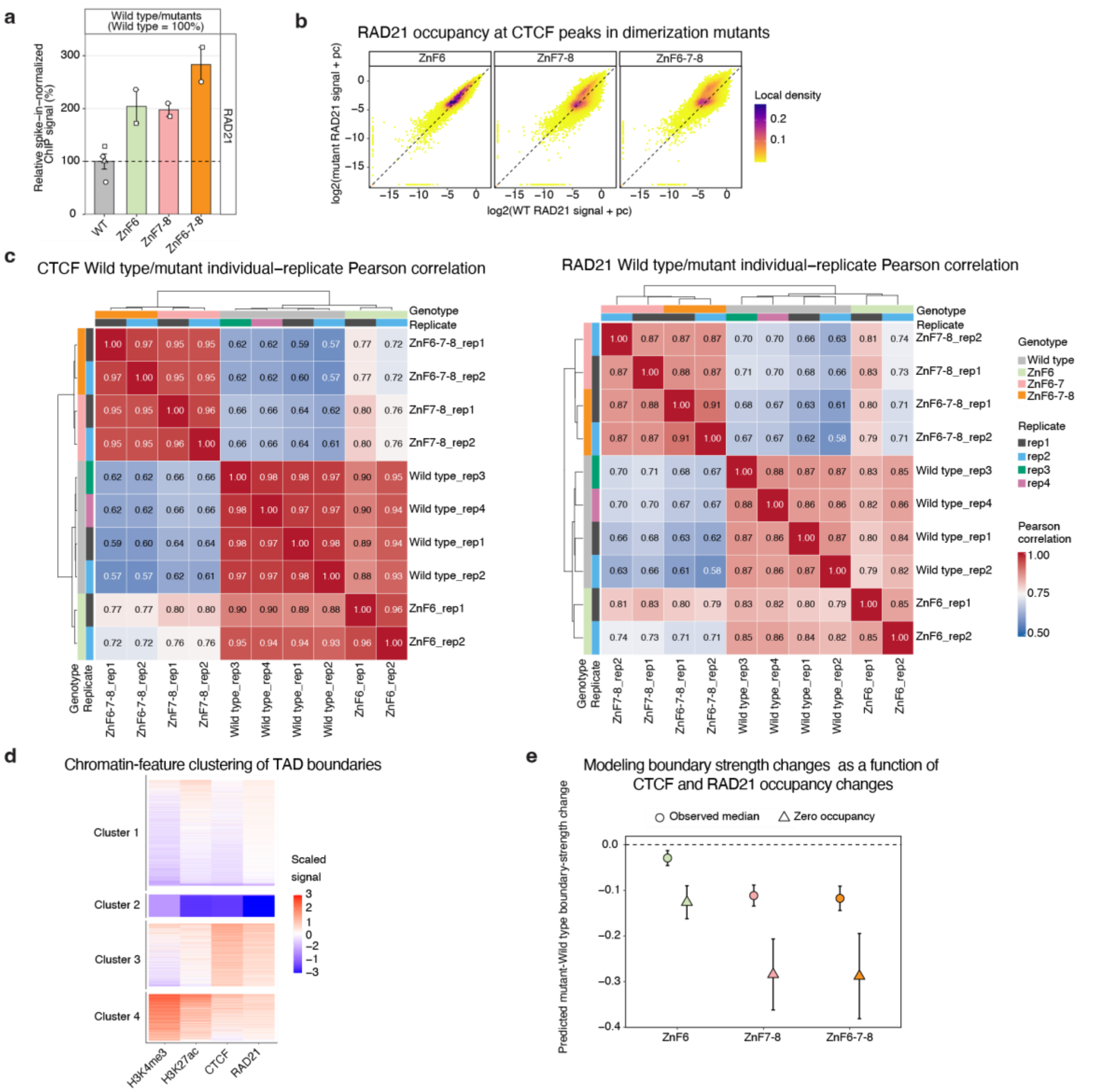

**Extended Data Figure 11. Spike-in-normalized CTCF and RAD21 occupancy and occupancy-adjusted boundary analysis in CTCF dimerization-interface mutants.**

**a,** Relative spike-in-normalized RAD21 ChIP-seq signal in wild-type and the indicated CTCF dimerization-interface mutants. Signals are expressed relative to the mean wild-type signal, which was set to 100% (dashed line). Bars represent mean and error bars indicate standard error; points represent independent biological replicates.

**b,** Peak-level comparison of spike-in-normalized RAD21 signal in each mutant versus wild type across the common CTCF peak set. Signals are shown as $\log_2$(signal +

protein-specific pseudocount). Each point represents one CTCF peak; the dashed line indicates y = x and color indicates local point density.

**c,** Pearson correlation matrices of CTCF (left) and RAD21 (right) ChIP–seq signals across individual biological replicates. Correlations were calculated from $\log_2$(signal + protein-specific pseudocount) values at the common CTCF peak set. Samples were hierarchically clustered using 1 − Pearson correlation as the distance metric and average linkage. Annotation bars indicate genotype and replicate.

**d,** K-means clustering of called TAD boundaries according to H3K4me3, H3K27ac, CTCF and RAD21 ChIP-seq signals. Signals were log-transformed and standardized across boundaries before clustering into four groups. Each row represents one boundary; boundaries are grouped by cluster and ordered by mean scaled signal within each cluster. Cluster 3 represents the CTCF/RAD21-rich boundary class used for the analysis in (**e**).

**e,** Occupancy-adjusted change in boundary-strength score within the CTCF/RAD21-rich boundary class. Mutant-wild-type changes in boundary strength were modeled as a function of the corresponding changes in CTCF and RAD21 signal. Circles show model predictions at the observed median ΔCTCF and ΔRAD21 for each mutant, whereas triangles show predictions at ΔCTCF = ΔRAD21 = 0, representing matched CTCF and RAD21 occupancy; error bars indicate 95% confidence intervals, and the dashed line indicates no change. The y-axis represents an absolute difference in boundary-strength score. The median wild-type boundary-strength score in this boundary class was approximately 0.76.

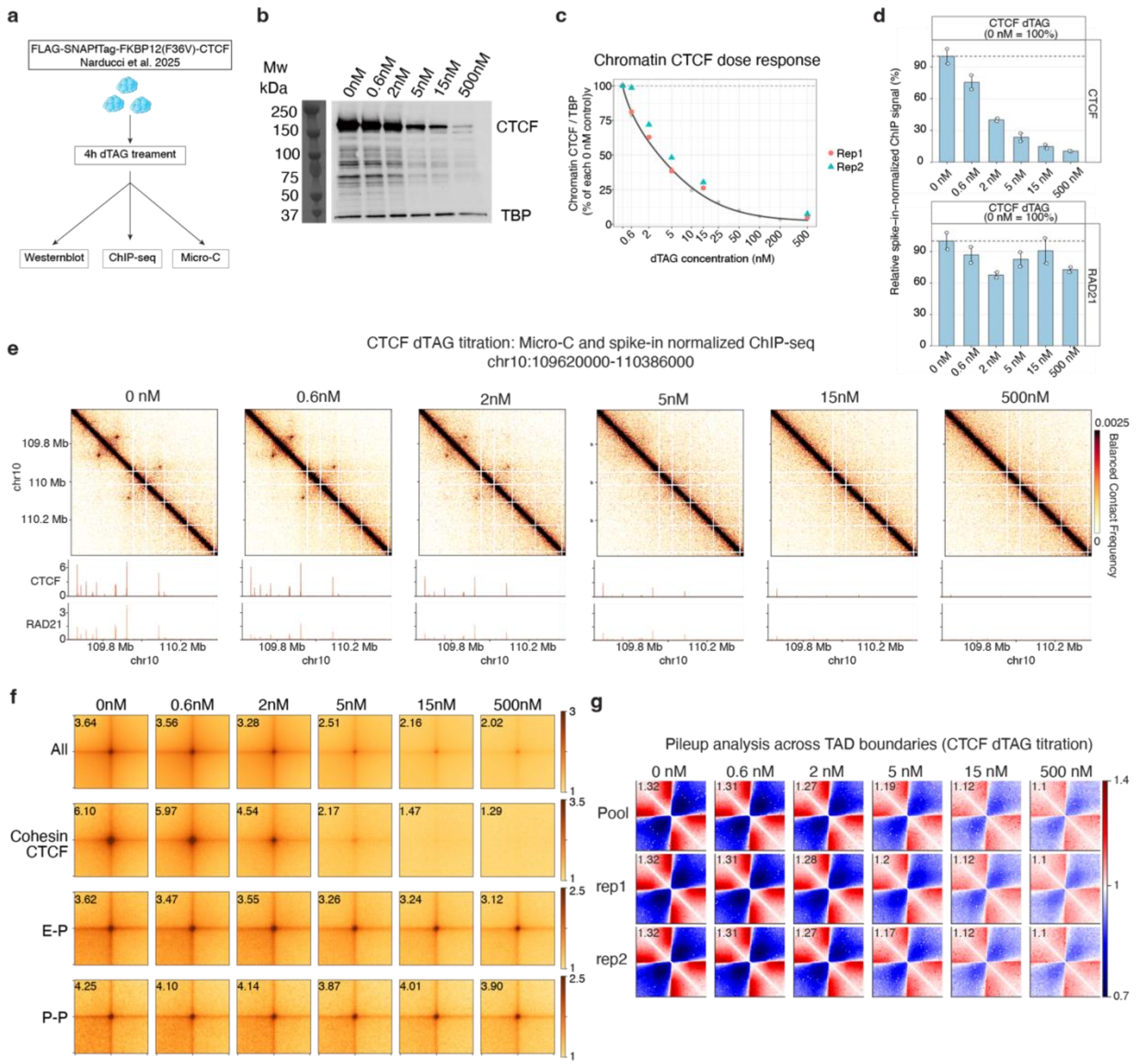


**Extended Data Figure 12. Graded acute CTCF depletion produces dose-dependent weakening of CTCF-cohesin loops and TAD boundaries.**

**a,** Schematic of the CTCF-dTAG titration experiment. FLAG-SNAPfTag-FKBP12(F36V)-CTCF mESCs[62] were treated for 4 h with the indicated dTAG concentrations and analyzed by Western blotting, spike-in-normalized CTCF and RAD21 ChIP-seq, and Micro-C.

**b,** Representative Western blots of chromatin fractions from CTCF-dTAG mESCs treated with the indicated dTAG concentrations. Blots were probed against CTCF and TBP; TBP was used as a loading control.

**c,** Quantification of chromatin-associated CTCF across the dTAG titration. CTCF abundance was normalized to TBP and expressed relative to the 0 nM control. Gray points indicate measurements from a preliminary titration experiment and the dark gray line indicates the fitted dose-response curve; colored points indicate two independent chromatin-fraction titration experiments.

**d,** Relative spike-in-normalized CTCF (top) and RAD21 (bottom) ChIP-seq signals following treatment with 0, 0.6, 2, 5, 15 or 500 nM dTAG. Signals are expressed relative to the mean 0 nM signal, which was set to 100% (dashed line). Bars represent mean and error bars indicate standard error, points represent independent biological replicates.

**e,** Representative balanced Micro-C contact maps across chr10:109,620,000-110,386,000 following treatment with the indicated dTAG concentrations. Contact maps are shown at 2-kb resolution. Corresponding spike-in-normalized CTCF and RAD21 ChIP-seq tracks are shown below each map.

**f,** Aggregate loop pileups for the indicated loop classes across the CTCF-dTAG titration. Rows show all loops (“All”), CTCF-cohesin loops (“Cohesin CTCF”), enhancer-promoter loops (E-P) and promoter-promoter loops (P-P). Columns show the indicated dTAG concentrations. The number in the upper left of each pileup indicates aggregate peak enrichment measured at the central pixel.

**g,** Aggregate contact maps across called TAD boundaries following treatment with the indicated dTAG concentrations. Rows show pooled data (Pool) and the two independent Micro-C replicates (rep1 and rep2). Numbers indicate intra-domain contact enrichment.

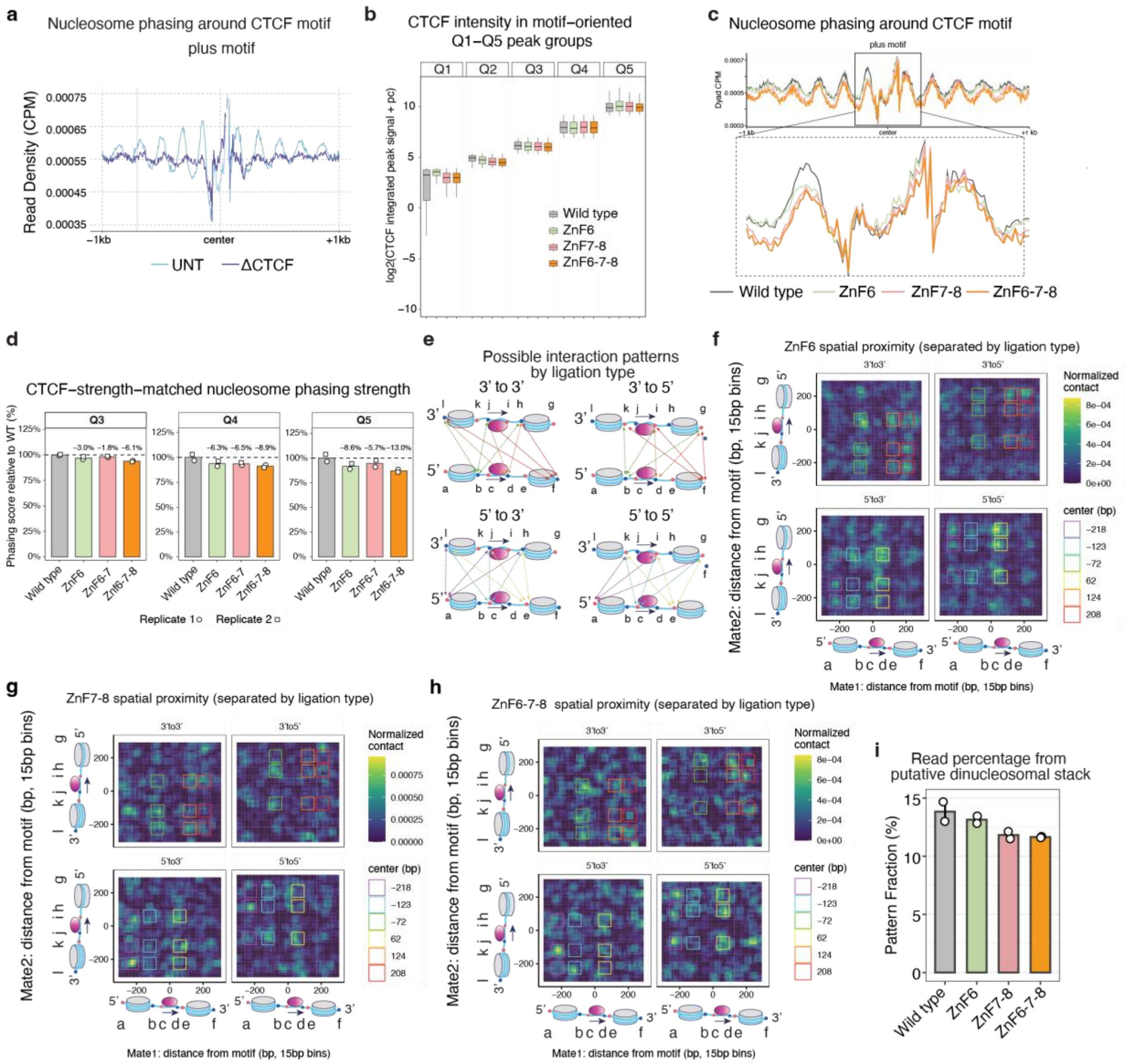


**Extended Data Figure 13. Effects of dimerization-interface disruption and acute CTCF depletion on nucleosome phasing and read-level ligation geometry.**

**a,** Nucleosome phasing around oriented CTCF motifs in UNT and ΔCTCF, shown separately for plus- and minus-oriented motifs.

**b**, Distribution of spike-in-normalized CTCF signal across five CTCF binding-strength groups (Q1-Q5) at plus-oriented motifs. Within each quantile, distributions are shown for wild type, ZnF6, ZnF7-8 and ZnF6-7-8. In box plots, center lines denote medians, hinges denote first and third quartiles, and whiskers extend to the most extreme values within 1.5× the interquartile range.

**c,** Nucleosome phasing around plus-oriented CTCF motifs at CTCF-strength-matched Q4 sites. Lines show the mean dyad CPM profiles for wild-type, ZnF6, ZnF7-8 and ZnF6-7-8 cells. The upper panel shows profiles within ±1 kb of the motif center, and the lower panel shows an expanded view of the central ±250-bp region indicated by the box in the upper panel.

**d,** Quantification of nucleosome phasing at plus-oriented Q3-Q5 CTCF sites after matching CTCF binding strength. Q1 and Q2 were excluded because CTCF signal distributions were not adequately matched between wild-type and the mutants in these groups (**b**). The phasing score was calculated as the summed dyad CPM within ±30 bp of the wild type-defined −1 and +1 nucleosome peaks, corresponding to −175 to −115 bp and +105 to +165 bp relative to the motif center. Within each quantile, scores were normalized to the wild type median, set to 100%. Bars show median values and points represent independent Micro-C replicates. Labels indicate the median percentage change relative to wild type.

**e,** Schematic of ligation-orientation classes (3′-3′, 3′-5′, 5′-3′, and 5′-5′) used to stratify read pairs by junction orientation and likely ligation events, assuming nucleosome dimer-stack geometry.

**f,g,h** Read-level spatial proximity maps separated by ligation-orientation class for mutants ZnF6 (**f**), ZnF7-8 (**g**), and ZnF6-7-8 (**h**). Heatmaps show motif-centered distance-distance distributions (15-bp bins), with colored boxes indicating the positional windows used for enrichment quantification in (**i**).

**i,** Fraction of read pairs assigned to the expected positional windows defined in **f-h** and **Figure 5e**, plotted for wild type and mutants. Points indicate independent Micro-C replicates and dark red dots represent the mean.

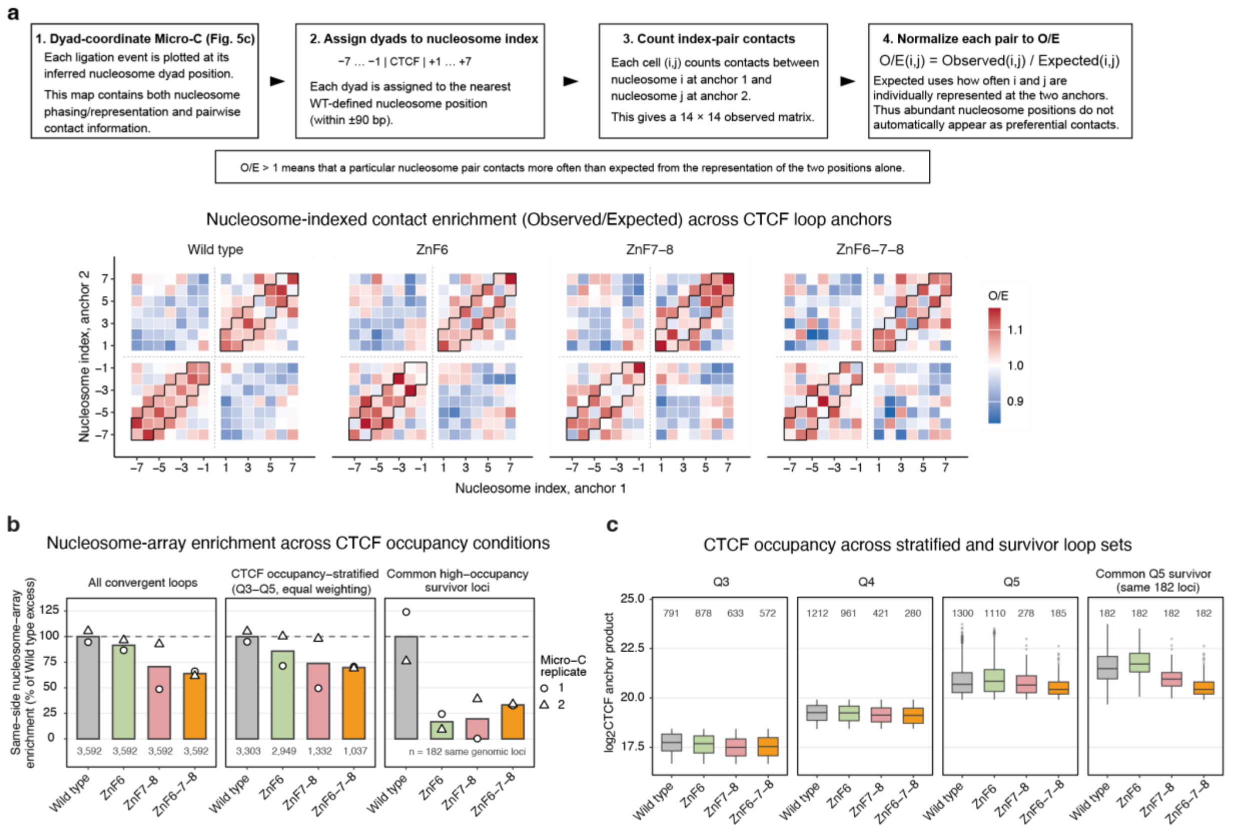


**Extended Data Figure 14. Extended nucleosome-array organization across CTCF loop anchors and its relationship to CTCF occupancy**.

**a,** Nucleosome-indexed contact enrichment across CTCF loop anchors. Inferred dyads were assigned to the nearest wild-type–defined phased nucleosome position within a 180-bp window centered on each peak, spanning nucleosome indices -7 to -1 and +1 to +7. As illustrated in the schematic, dyad-coordinate contacts were first collapsed into nucleosome-index pairs, and the observed frequency of each pair was then divided by the frequency expected from the independent representation of the two nucleosome positions. This normalization accounts for position-dependent differences in read abundance, including the higher representation of nucleosomes proximal to CTCF, and therefore highlights nucleosome pairs that contact more frequently than expected from their individual representation. Black outlines indicate the same-side contacts used for quantification in **b**, allowing the two nucleosome indices to differ by at most one. Contacts spanning the central −1/+1 CTCF interval were excluded from this definition.

**b,** Quantification of same-side nucleosome-array enrichment. Observed/expected (O/E) values were averaged over the outlined cells in **a**, and enrichment above the expected value (O/E = 1) was expressed relative to the corresponding wild-type enrichment. Left, all 3,592 convergent loops. **Middle,** loops stratified by the product of CTCF occupancy at the two loop anchors; O/E was calculated independently within globally defined Q3–Q5 occupancy strata and the three strata were combined with equal weighting. This comparison therefore matches CTCF-occupancy ranges across genotypes while allowing different genomic loci to contribute. **Right,** a more stringent same-locus comparison: 182 rare loops that remained in the highest CTCF-product stratum in both ZnF7-8 and ZnF6-7-8 were selected, and these exact same genomic loci were evaluated in all four genotypes. Unlike the Q3–Q5 analysis, which allows different loci to contribute within similar CTCF-occupancy ranges, this comparison fixes the genomic loci across genotypes while focusing on a rare subset that retains high CTCF occupancy in ZnF7-8 and ZnF6-7-8 mutants. Bars indicate genotype means and symbols indicate the two independent Micro-C biological replicates.

**c,** CTCF occupancy distributions for the loop sets used in the occupancy-controlled analyses in **b**. Loop-level CTCF occupancy was defined as the product of the spike-in-normalized CTCF signals at the two anchors and is shown on a $\log_2$ scale. Q3-Q5 were defined using a single set of quantile boundaries derived from the pooled CTCF-product distribution across all four genotypes. The final facet shows CTCF occupancy at the same 182 common high-occupancy survivor loci used in **b**. Numbers above boxes indicate the number of loops.
