## Supplementary Information Figures for "Cooperative CTCF-nucleosome oligomerization stabilizes chromatin loop anchors"

#### Supplementary Figure 1

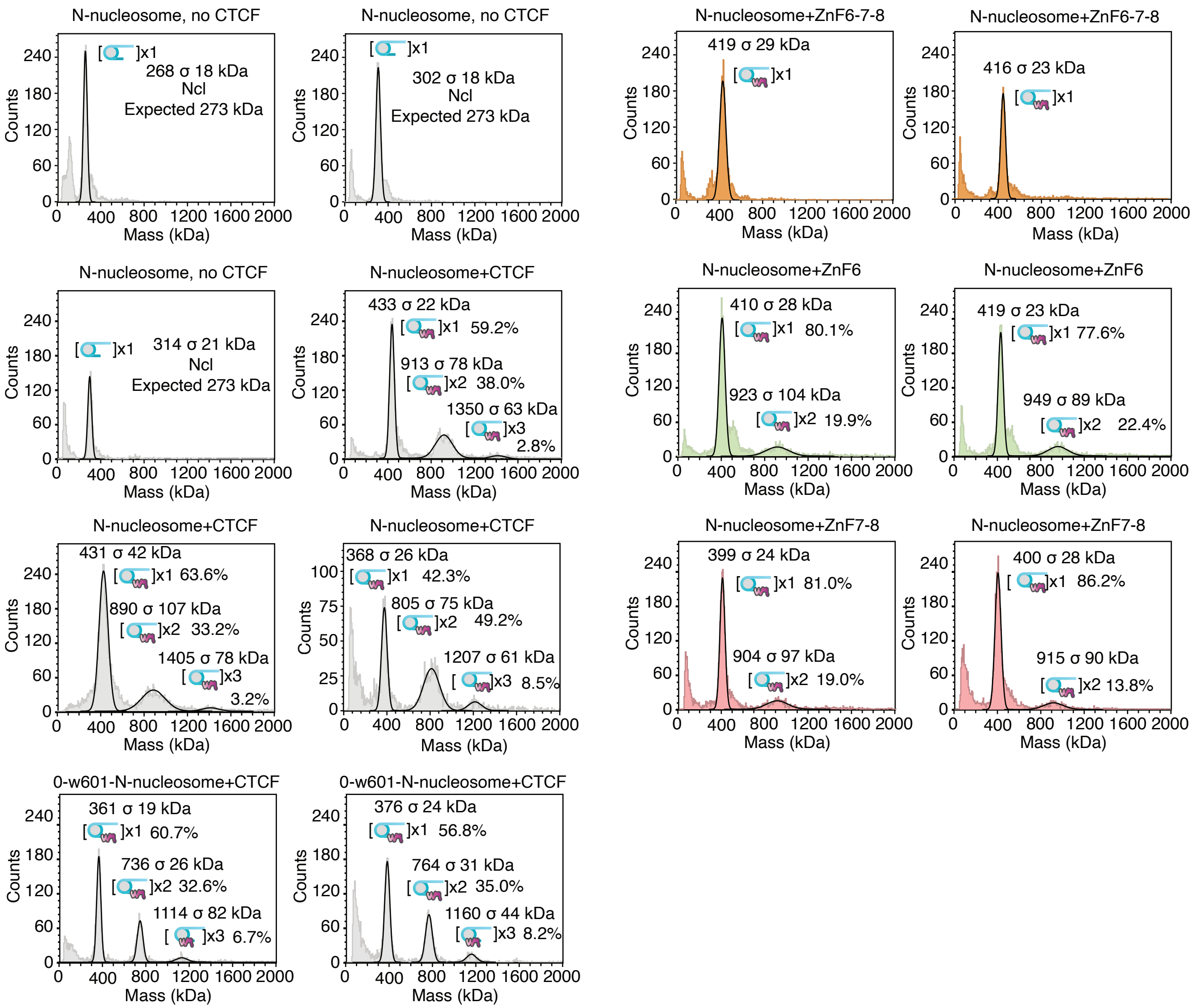

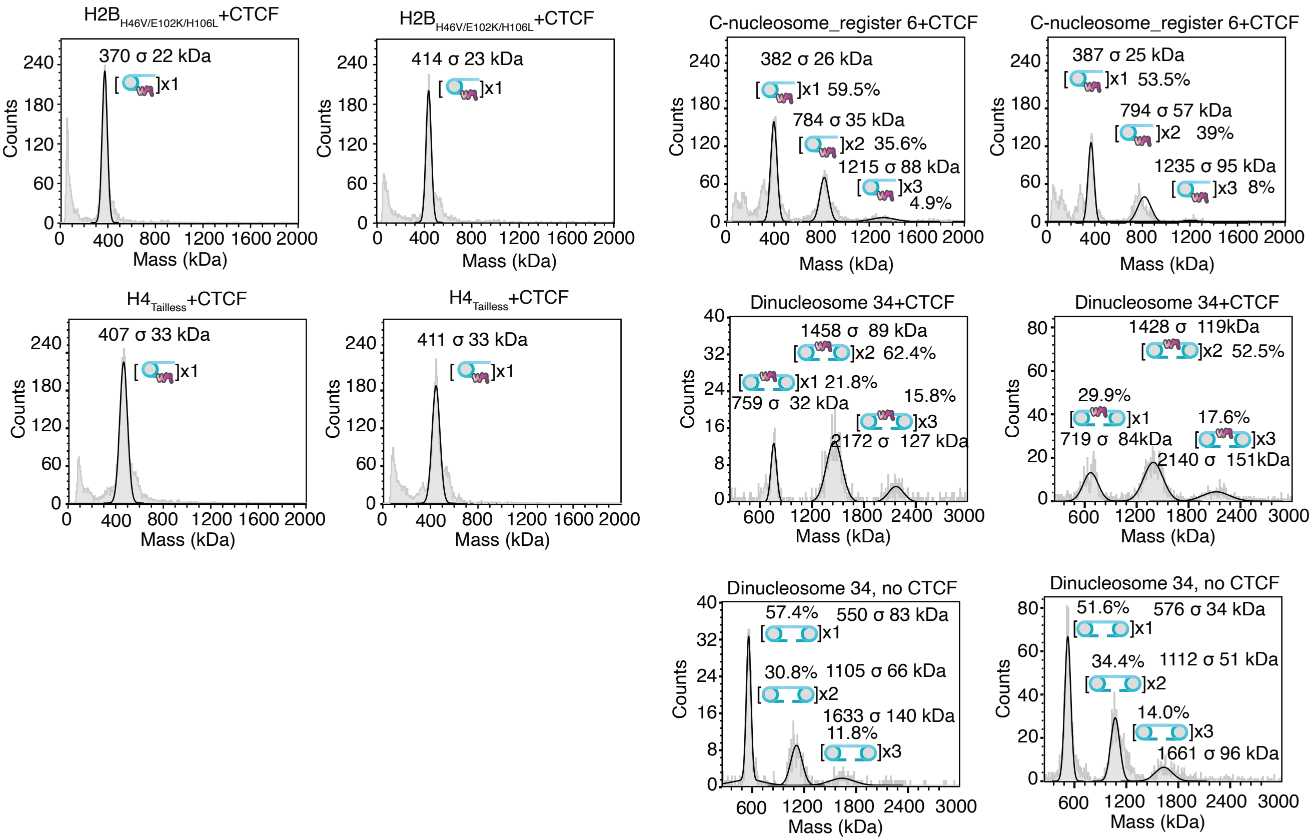


**Supplementary Figure 1**. Native non-crosslinked mass photometry measurements. Calculated masses corresponding to free nucleosome and nucleosome-CTCF complexes are shown. Percentages of different species are indicated.

### Supplementary Figure 2

Extended Data Figure 1a
purified proteins

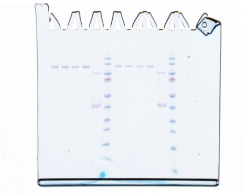

Extended Data Figure 1d
crosslinked after gradient

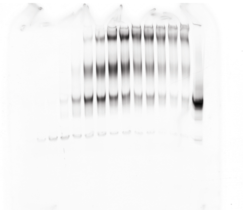

Figure 1i. N-nucleosome+CTCF

Replicate 1 Replicate 2 Replicate 3

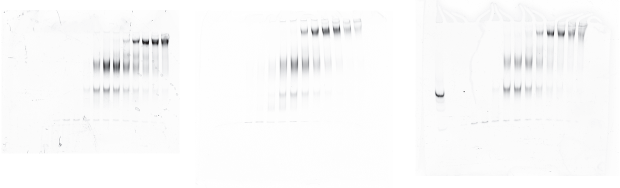

Extended Data Figure 1f. N-nucleosome, no CTCF

Replicate 1 Replicate 2 Replicate 3

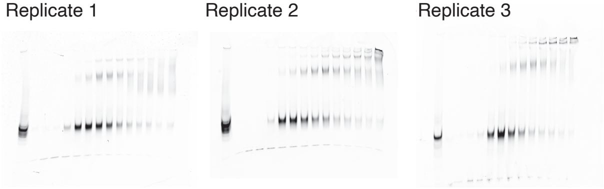

Extended Data Figure 1i. 0-W601-N-nucleosome+CTCF

Replicate 1 Replicate 2 Replicate 3

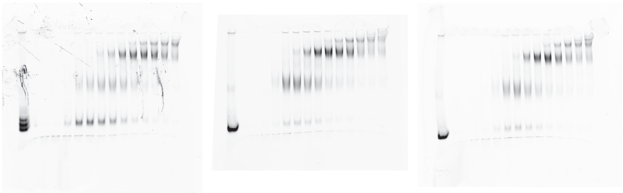

Figure 1k. Random-nucleosome+CTCF

Replicate 1 Replicate 2 Replicate 3

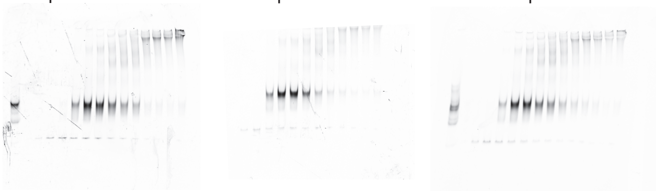

Figure 1m. Linear DNA+CTCF

Replicate 1 Replicate 2 Replicate 3

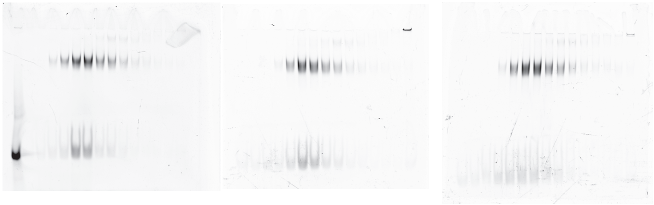

Extended Data Figure 5a. H2B$_{H46V/H106L/E102K}$+CTCF

Replicate 1

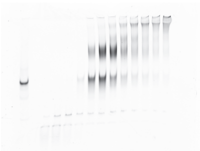

Replicate 2

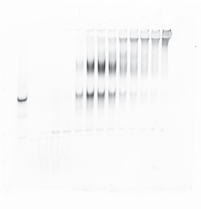

Replicate 3

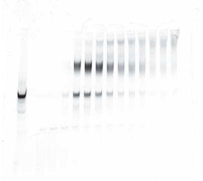

Figure 5b. H4 tailless+CTCF

Replicate 1

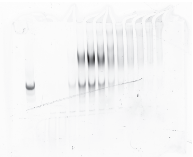

Replicate 2

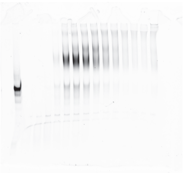

Replicate 3

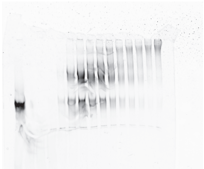

Extended Data Figure 5d. N-nucleosome+ZnF6mut

Replicate 1

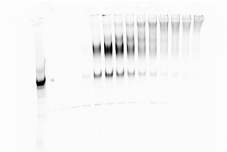

Replicate 2

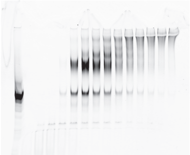

Replicate 3

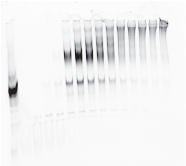

Extended Data Figure 5e. N-nucleosome+ZnF7-8mut

Replicate 1

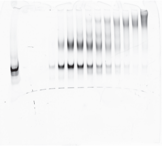

Replicate 2

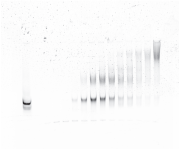

Replicate 3

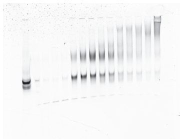

Figure 5f. N-nucleosome+ZnF6-7-8mut

Replicate 1

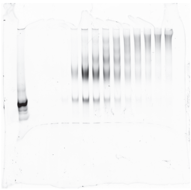

Replicate 2

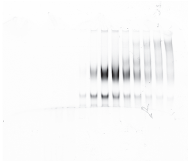

Replicate 3

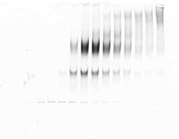

Extended Data Figure 5l. N-nucleosome+ZnF1-11

Replicate 1

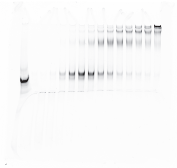

Replicate 2

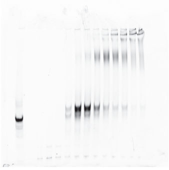

Replicate 3

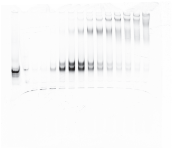

Extended Data Figure 6c. C-nucleosome_register 0+CTCF

Replicate 1

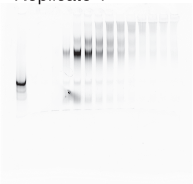

Replicate 2

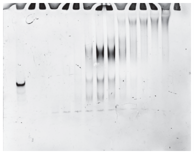

Replicate 3

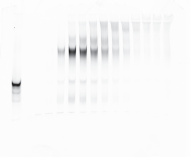

Extended Data Figure 6d. C-nucleosome_register 6+CTCF

Replicate 1

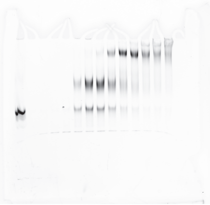

Replicate 2

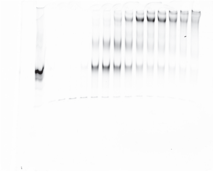

Replicate 3

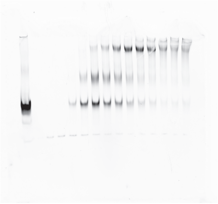

Extended Data Figure 6e. C-nucleosome_register 7 + wildtype CTCF

Replicate 1

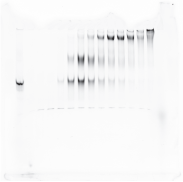

Replicate 2

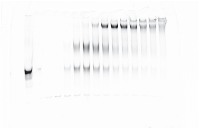

Replicate 3

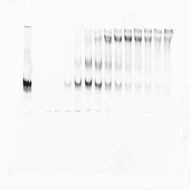

Figure 3d. Dinucleosome 34, no CTCF

Replicate 1 Replicate 2 Replicate 3

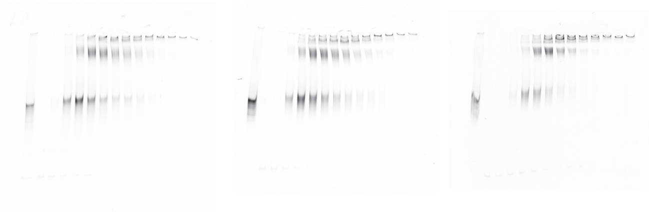

Figure 3e. Dinucleosome 34+CTCF wild type

Replicate 1 Replicate 2 Replicate 3

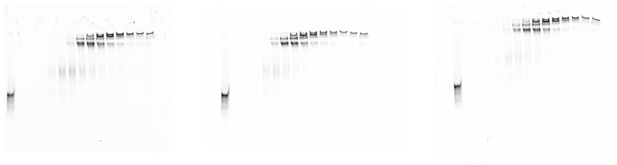

Extended Data Figure 7b. Native 80, no CTCF

Replicate 1 Replicate 2 Replicate 3

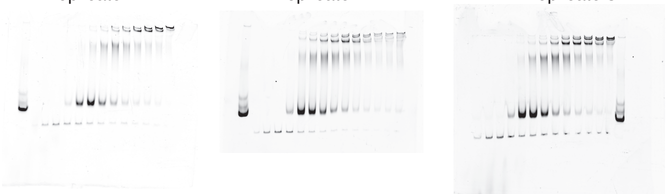

Extended Data Figure 7c. Native 80+CTCF wild type

Replicate 1 Replicate 2 Replicate 3

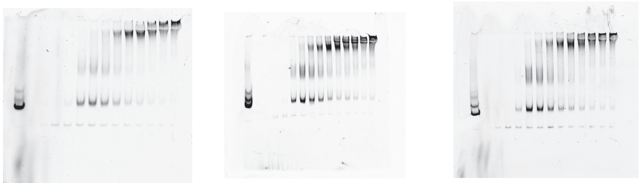

Extended Data Figure 7d. Native 110, no CTCF

Replicate 1 Replicate 2 Replicate 3

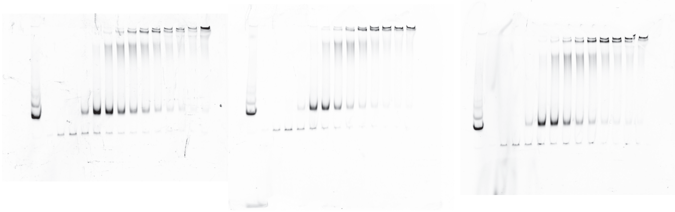

Extended Data Figure 7e. Native 110+CTCF wild type

Replicate 1 Replicate 2 Replicate 3

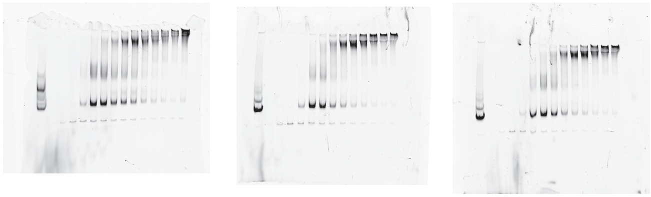

Extended Data Figure 7g. Dinucleosome 42, no CTCF

Replicate 1 Replicate 2 Replicate 3

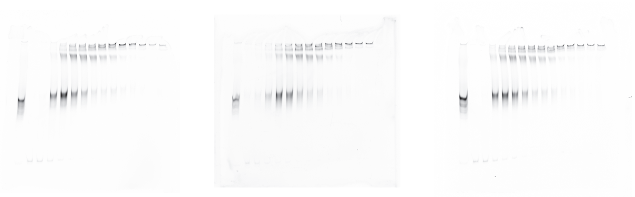

Extended Data Figure 7h. Dinucleosome 42+wildtype CTCF

Replicate 1 Replicate 2 Replicate 3

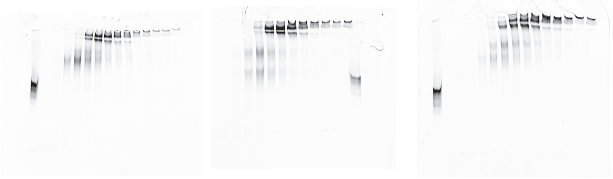

**Supplementary Figure 2**. Uncropped gel images from SDS and native PAGE

### Supplementary Figure 3

Raw images of Western blot

Nuclear Fraction
Biological rep1 251023

Membrane1

Marker

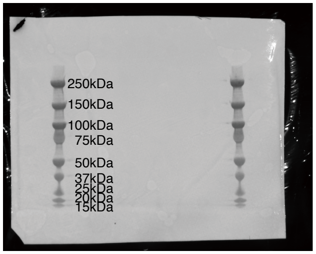


Signal

anti-CTCF + anti-TBP

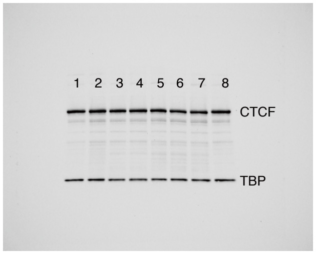


From Left to Right
1. e7mut_G1
2. e7mut_A4
3. e8mut_F1
4. e8mut_H8
5. e7e8mut_F1D11
6. e7e8mut_F1F11
7. e7e8mut_H8H8
8. WT

Chromatin Fraction
Biological rep1 251027

Membrane2

Marker

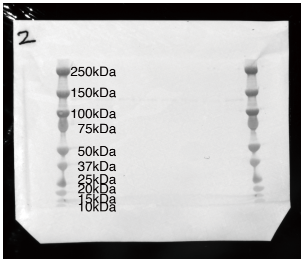


Signal

anti-CTCF + anti-TBP

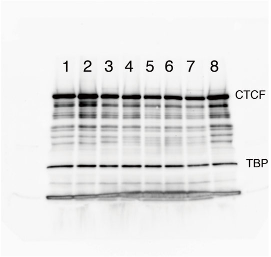


From Left to Right
1. e7mut_G1
2. e7mut_A4
3. e8mut_F1
4. e8mut_H8
5. e7e8mut_F1D11
6. e7e8mut_F1F11
7. e7e8mut_H8H8
8. WT

Nuclear Fraction
Biological rep2 251112

Membrane1

Marker

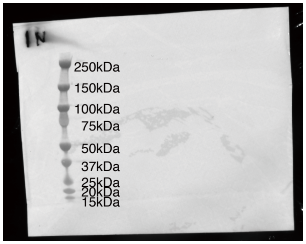


Signal

anti-CTCF + anti-TBP

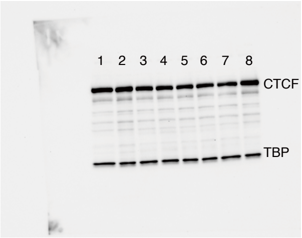


From Left to Right
1. e7mut_G1
2. e7mut_A4
3. e8mut_F1
4. e8mut_H8
5. e7e8mut_F1D11
6. e7e8mut_F1F11
7. e7e8mut_H8H8
8. WT

Chromatin Fraction
Biological rep2 251112

Membrane2

Marker

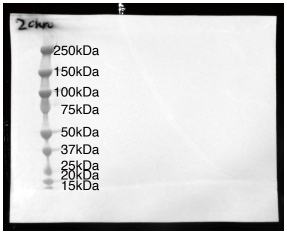


Signal

anti-CTCF + anti-TBP

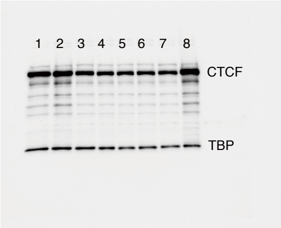


From Left to Right
1. e7mut_G1
2. e7mut_A4
3. e8mut_F1
4. e8mut_H8
5. e7e8mut_F1D11
6. e7e8mut_F1F11
7. e7e8mut_H8H8
8. WT

Raw images of Westernblot for dTAG titration experiments

Chromatin Fraction
260505

Marker

Signal
anti-CTCF + anti-TBP

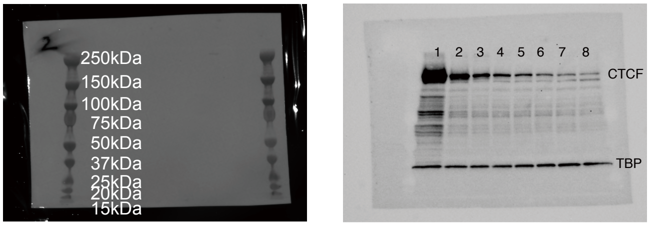


From Left to Right
1. ESC_Ctcf_dTAG_4h_0nM_Chromatin_1
2. ESC_Ctcf_dTAG_4h_5nM_Chromatin_1
3. ESC_Ctcf_dTAG_4h_10nM_Chromatin_1
4. ESC_Ctcf_dTAG_4h_25nM_Chromatin_1
5. ESC_Ctcf_dTAG_4h_50nM_Chromatin_1
6. ESC_Ctcf_dTAG_4h_100nM_Chromatin_1
7. ESC_Ctcf_dTAG_4h_200nM_Chromatin_1
8. ESC_Ctcf_dTAG_4h_500nM_Chromatin_1

Chromatin Fraction
260513

Marker

Signal
anti-CTCF + anti-TBP

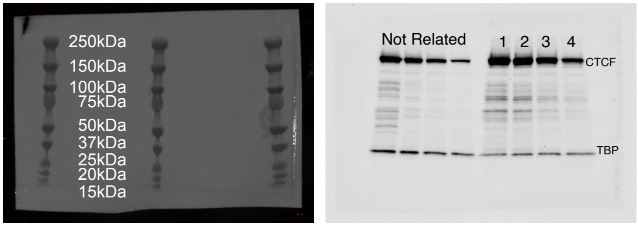


From Left to Right
1. ESC_Ctcf_dTAG_4h_0nM_Chromatin_1
2. ESC_Ctcf_dTAG_4h_0.6nM_Chromatin_1
3. ESC_Ctcf_dTAG_4h_2nM_Chromatin_1
4. ESC_Ctcf_dTAG_4h_5nM_Chromatin_1

Chromatin Fraction
Biological rep1 260527

Marker

Signal
anti-CTCF + anti-TBP

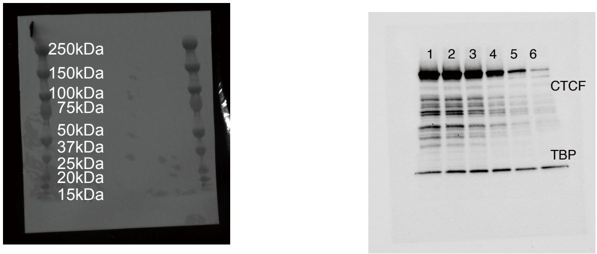


From Left to Right
1. ESC_Ctcf_dTAG_4h_0nM_Chromatin_1
2. ESC_Ctcf_dTAG_4h_0.6nM_Chromatin_1
3. ESC_Ctcf_dTAG_4h_2nM_Chromatin_1
4. ESC_Ctcf_dTAG_4h_5nM_Chromatin_1
5. ESC_Ctcf_dTAG_4h_15nM_Chromatin_1
6. ESC_Ctcf_dTAG_4h_500nM_Chromatin_1

Chromatin Fraction
Biological rep2 260528

Membrane2

Marker

Signal
anti-CTCF + anti-TBP

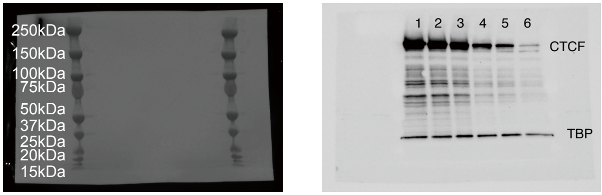


From Left to Right
1. ESC_Ctcf_dTAG_4h_0nM_Chromatin_2
2. ESC_Ctcf_dTAG_4h_0.6nM_Chromatin_2
3. ESC_Ctcf_dTAG_4h_2nM_Chromatin_2
4. ESC_Ctcf_dTAG_4h_5nM_Chromatin_2
5. ESC_Ctcf_dTAG_4h_15nM_Chromatin_2
6. ESC_Ctcf_dTAG_4h_500nM_Chromatin_2

**Supplementary Figure 3**. Raw images of Western blot analyses. Samples, antibodies used, and molecular weight markers are indicated.

#### Supplementary Figure 4

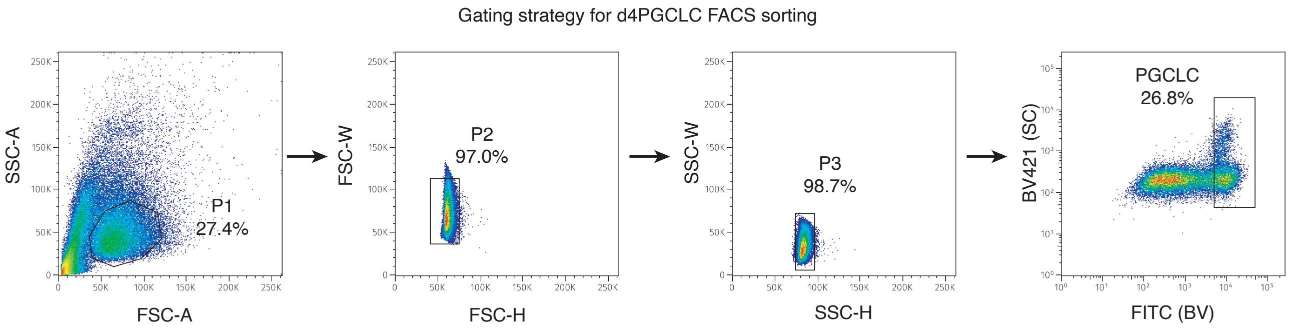


**Supplementary Figure 4**. Fluorescence-activated cell sorting (FACS) analysis of primordial germ cell-like cells induction.
